# Brain networks in human conscious visual perception

**DOI:** 10.1101/2021.10.04.462661

**Authors:** Sharif I. Kronemer, Mark Aksen, Julia Ding, Jun Hwan Ryu, Qilong Xin, Zhaoxiong Ding, Jacob S. Prince, Hunki Kwon, Aya Khalaf, Sarit Forman, David Jin, Kevin Wang, Kaylie Chen, Claire Hu, Akshar Agarwal, Erik Saberski, Syed Mohammad Adil Wafa, Owen P. Morgan, Jia Wu, Kate L. Christison-Lagay, Nicholas Hasulak, Martha Morrell, Alexandra Urban, R. Todd Constable, Michael Pitts, R. Mark Richardson, Michael J. Crowley, Hal Blumenfeld

## Abstract

Consciousness is not explained by a single mechanism, rather it involves multiple specialized neural systems overlapping in space and time. We hypothesize that synergistic, large-scale subcortical and cortical attention and signal processing networks encode conscious experiences. To identify brain activity in conscious perception without overt report, we classified visual stimuli as perceived or not using eye measurements. Report-independent event-related potentials and functional magnetic resonance imaging (fMRI) signals both occurred at early times after stimuli. Direct recordings revealed a novel thalamic awareness potential linked to conscious visual perception based on report. fMRI showed thalamic and cortical detection, arousal, attentional salience, task-positive, and default mode networks were involved independent of overt report. These findings identify a specific sequence of neural mechanisms in human conscious visual perception.

**One-Sentence Summary:** Human conscious visual perception engages large-scale subcortical and cortical networks even without overt report.

## Main Text

Consciousness is central to human experience yet is not easily explained. Theories of consciousness typically emphasize a single mechanism hoping to solve the mystery linking brain activity to conscious experience. However, single-mechanism theories or models have so far not achieved convincing success. Instead, we posit that consciousness is best understood through a synergistic combination of multiple mechanisms overlapping in space and time. Just as several processes in biology together distinguish living from non-living things, multiple mechanisms in neuroscience combine to separate conscious from non-conscious neural activity. Specifically, conscious awareness of discrete events lies at the temporal nexus of attention and memory, both major fields in modern neuroscience that contribute to understanding consciousness. We hypothesize that systems crucial for consciousness include: (1) attention mechanism mediating signal detection, dynamic modulation of arousal, and bottom-up plus top-down attentional control, overlapping in space and time with; (2) systems that limit competing signals (e.g., through reduced default mode network activity); and, finally, (3) hierarchically organized systems that fully process signals for memory encoding and subsequent report (*1*).

To investigate these multiple systems, a comprehensive approach is needed to identify activity throughout the brain at high spatial and temporal resolution specifically related to consciousness. We aimed to overcome several limitations of prior studies. For example, subcortical regions are poorly understood in human conscious perception and are often relegated to preconscious state-based precursors of consciousness (*2–4*). We therefore investigated dynamic changes in both subcortical and cortical systems using techniques with complementary strengths. These included large sample scalp electroencephalography (EEG) and depth recordings from the human thalamus providing direct measurements of neural activity at high time resolution, and large sample functional magnetic resonance imaging (fMRI) analyzed with data-driven approaches providing comprehensive mapping of the whole brain. Most importantly for linking these neural measures with consciousness, we used a threshold visual perception task to measure brain signals produced by physically identical stimuli that are either perceived versus not perceived, coupled with a unique innovation to remove the confound of overt report. When participants are asked to overtly report whether they perceived a stimulus this introduces post-perceptual processes (e.g., decision making and motor planning) that can confound signals linked to consciousness, even when the report is delayed by several seconds after the stimulus (*5–7*).

To address the confound of post-perceptual processing we developed a novel no-report paradigm using transient changes in pupillometry and eye tracking to classify stimuli as perceived or not perceived without overt report (*8*) (fig. S1-9; Tables S4-6). In a previously established Report Paradigm (*9*), participants were repeatedly shown identical faces at 50% perceptual threshold (Fig. 1A). This resulted in approximately equal numbers of perceived and not perceived stimuli based on overt report of stimulus presence and location (fig. S6, 7). We formed a novel combined Report + No-Report Paradigm by maintaining the report, task-relevant stimuli from the Report Paradigm and introducing identical no-report, task-irrelevant faces (i.e., stimuli that did not require overt report) interleaved with task-relevant stimuli, distinguished by different screen locations (four quadrant locations and four central locations; Fig. 1B). For example, in one session participants were told to report only on stimuli appearing in the screen quadrant locations and not in the screen central locations (fig. S1A; see (*8*) for counterbalancing and details). The perception of no-report stimuli was determined by classification of pupillometry and eye tracking during the task. Specifically, we found pupil dilation, blink rate increases, and microsaccade rate decreases for consciously perceived visual stimuli, present irrespective of task relevance (Fig. 1C, D). Pupillometry and eye tracking have been previously implemented as covert measures of consciousness (*10, 11*), however, the dynamics reported here are unique because they do not rely on a task sequence (e.g., no-report stimuli always first), perceptual switching (e.g., binocular rivalry), nor stimulus type (e.g., stimulus modality), as we have found similar pupil, blink, and microsaccade responses for perceived auditory and tactile stimuli (*8, 12, 13*).

**Fig. 1.**
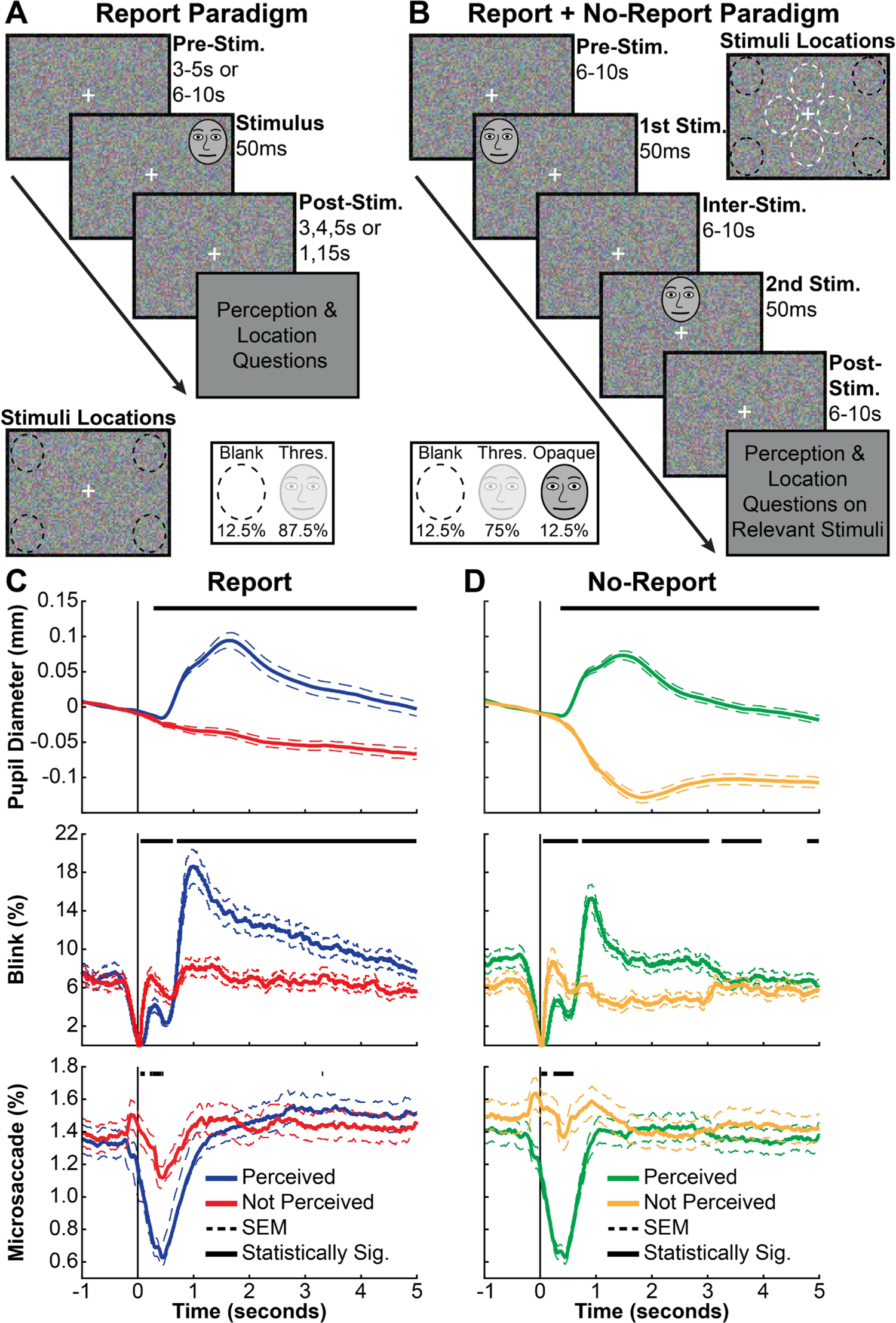
Report-dependent and report-independent behavioral paradigms for conscious perception; similar pupillometry and eye tracking responses. (A) Report Paradigm trial sequence, stimulus locations, and opacity conditions. (B) Report + No-Report Paradigm trial sequence, stimulus locations, and opacity conditions. (C) Report (task-relevant) data with perception based on overt participant responses and (D) no-report (task-irrelevant) data with perception classified by machine learning yield similar pupil, blink, and microsaccade dynamics (*8*). Data in (C) and (D) are from stimuli at perceptual threshold (see inset) in the Report + No-Report Paradigm (N = 68 participants). Stimulus onset was at time = 0. Classification score threshold for (D) was 0 (fig. S9) (*8*). Statistically significant (Statistically Sig.) different time points between perceived and not perceived data are indicated (black bars above pupil, blink, and microsaccade timecourses) based on cluster-based permutation tests (*p* < 0.05) (*8*). Standard error of the mean (SEM). Note that the face photograph used as a visual stimulus in this study was removed from Figure 1 per *bioRxiv* policy and replaced by a sketch version. The actual face stimulus used can be found in (*9*).

Our first goal was to investigate report-dependent and report-independent brain signals at high time resolution. We recorded the following well established event-related potentials (ERPs) for perceived stimuli in the report data, in temporal sequence: (1) N100, (2) VAN (visual awareness negativity), (3) P2/N2, and (4) P3/LP (late potentials) (Fig. 2A; fig. S10A, 11A) (*14–17*). The earliest ERP (N100) is thought to be related to visual cortical activation, and was present for all stimuli regardless of perceived, not perceived, report or no-report, although the magnitude varied depending on conditions (fig. S10, 11). The VAN was significantly greater for perceived versus not perceived stimuli, both for report and no-report data (Fig. 2A, B; fig. S10A, C). Importantly, the VAN was not significantly different for report versus no-report data, whereas the later ERPs (P2/N2 and P3/LP) were significantly larger for perceived stimuli in the report data (Fig. 2C; fig. S10E). Our novel paradigm thus strengthens previous findings that early ERPs, particularly the VAN, are seen even under no-report or task-irrelevant conditions.

**Fig. 2.**
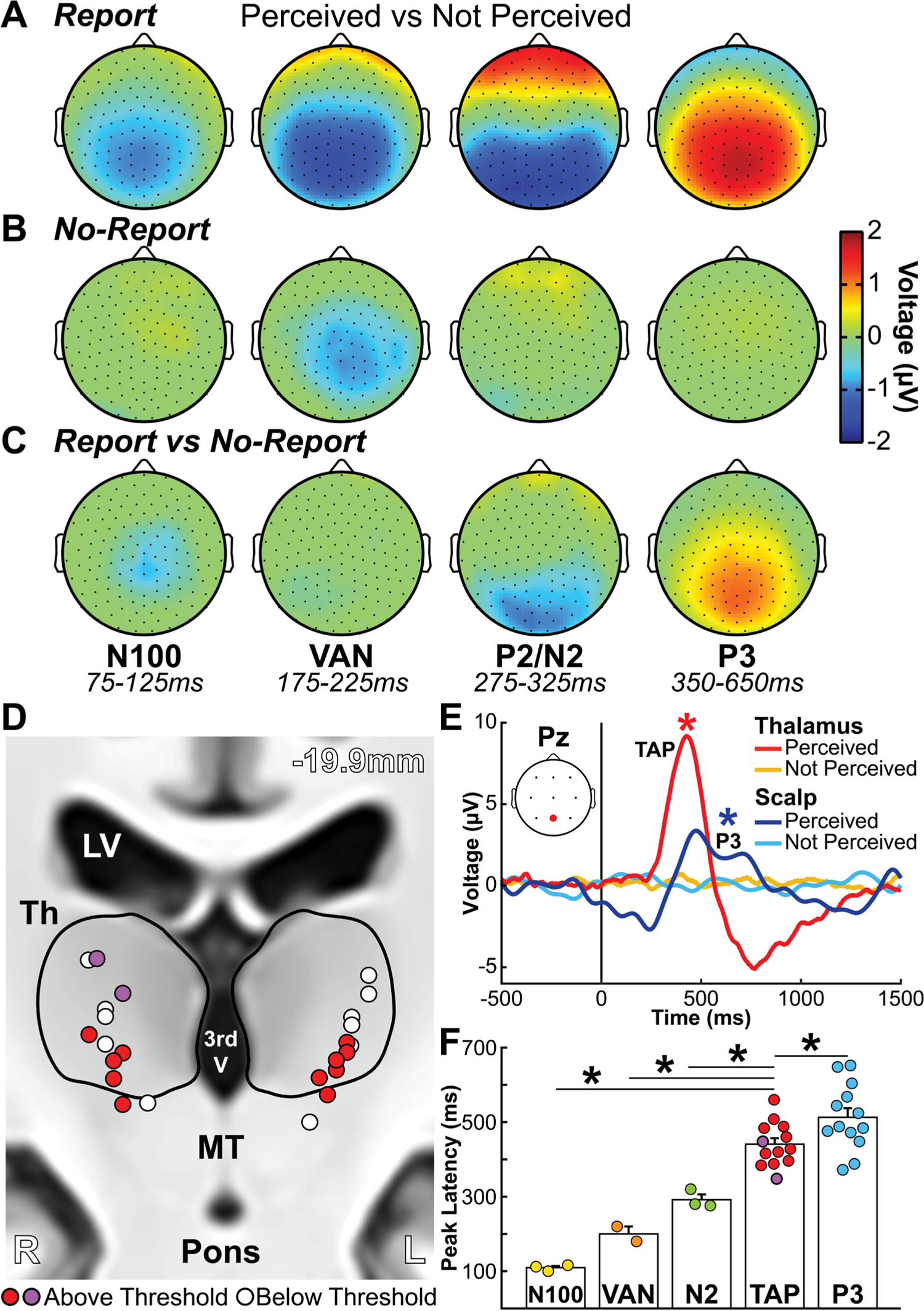
Cortical and thalamic electrophysiology signals in conscious perception. (**A-C**) Scalp topographical plots of high-density EEG showing mean voltage for all statistically significant samples by cluster-based permutation tests (*p* < 0.05) (*8*) in four time windows corresponding with the event-related potentials (ERPs) N100 (75-125ms), visual awareness negativity (VAN; 175-225ms), P2/N2 (275-325ms), and P3 (350-650ms) for perceived versus not perceived stimuli in (**A**) report, (**B**) no-report, and (**C**) report versus no-report data. The VAN is present both for report and no-report data, whereas later ERPs, especially N2 and P3 are report-dependent. (**D**) Subcortical depth channel locations collapsed onto a coronal slice (−19.9mm) in MNI brain template space (*8*). Red/purple (N = 13) and white bipolar channels (N = 11) distinguish above or below voltage threshold channels, respectively (fig. S12) (*8*). Red channels are with the RNS System (NeuroPace, Inc.) and purple channels with Natus NeuroWorks (Natus, Inc.) depth electrode recordings (Table S2). Neighboring anatomy includes lateral ventricles (LV), third ventricle (3^rd^ V), thalamus (Th), midbrain tegmentum (MT), and pons. (**E**) Thalamic awareness potential (TAP) and P3 are seen, respectively in thalamic above-threshold contacts and Pz (location on inset) scalp EEG contacts. Mean timecourses show significant differences for perceived versus not perceived stimuli in the ERP analysis for TAP and P3 by cluster-based permutation tests (* *p* < 0.05) (*8*). Stimulus onset was at time = 0. (**F**) Peak latencies from stimulus onset for scalp ERPs and TAP. Circles represent individual data (mean channel latencies) and error bars indicate standard error of the mean (SEM). Significantly different latencies from TAP found by Wilcoxon rank sum test with Holm-Bonferroni correction (* *p* < 0.05). (**A-C**) Data for report stimuli are from Report Paradigm (N = 57) and Report + No-Report Paradigm (N = 65); data for no-report stimuli are from Report + No-Report Paradigm (N = 65; 2 not perceived subject data sets were excluded because at the prediction score threshold of −0.75 there were no predicted not perceived trials; see the *Covert Prediction of Conscious Perception – Machine Learning Pipeline* section in (*8*)). (**D-F**) Data are from Report Paradigm in patient participants with thalamic depth electrodes (N = 7).

Therefore, these early signals reflect the scalp neurophysiological signatures for report-independent conscious perception (*18–20*). Furthermore, report and task-relevance introduce report-dependent changes dominated by later ERPs, including P3/LP that extend persistent neurophysiological activity for over 1 second after stimulus onset (fig. S11A; see also (*21*) and fig. S9 in (*9*)).

Having established early report-independent signals of conscious perception, and later report-dependent signals likely related to post-perceptual processing, we next sought to investigate subcortical signals selective for conscious perception and to determine their timing. A key subcortical brain structure for arousal and consciousness is the intralaminar thalamus (*22*).

Patient participants with chronically implanted deep brain recording and stimulation devices (RNS® System, NeuroPace, Inc.; Natus NeuroWorks, Inc.) for the treatment of epilepsy provided unique access to this region (*8*). We simultaneously recorded cortical electrophysiology from scalp EEG and subcortical signals from thalamic depth contacts (Fig. 2D; fig. S12, 13; Tables S2, 3) while participants completed the Report Paradigm. We found a novel, biphasic thalamic awareness potential (TAP) highly selective for perceived stimuli with an onset at ∼250ms and initial peak at ∼430ms post-stimulus presentation (Fig. 2E). TAP was localized to channels within or along the lateral border of the intralaminar thalamus (fig. S2D; fig. S12). TAP was also selectively present for perceived *auditory* stimuli in one participant (participant 1 in Table S2) who completed an analogous perceptual threshold auditory task in a separate study from our group (fig. S14) (*12*).

We next investigated the timing of TAP relative to scalp ERPs. First, we noted that the scalp ERPs recorded in the patient participants were similar to those of the healthy participants, despite different recording systems and sample sizes (fig. S13). We found that TAP preceded P3, but followed the N100, VAN, and N2 (Fig. 2F). Therefore, the timing of TAP was later than ERPs found in the prior experiment to be report-independent (VAN), and fell within or earlier than ERPs that were report-dependent (N2, P3). We did not directly test whether TAP was report-dependent because of limited recording time with the patient participants. However, because we hypothesized that TAP is one node in a broad subcortical arousal and salience network participating in attention state dynamics and consciousness (*23, 24*), we next used fMRI in a large cohort of healthy participants to investigate cortical and subcortical conscious perception-linked dynamics, with and without overt report.

We found a broad network of subcortical and cortical regions showing report-independent fMRI changes especially at earlier times after stimulus presentation. Perceived versus not perceived blood-oxygen-level-dependent (BOLD) responses were distinguished by early (<4 seconds) and late (>5 seconds) dynamics analyzed by spatiotemporal cluster-based permutation tests (*8*). At 3 seconds post-stimulus (Fig. 3A, C; fig. S15A, C; Slide S1, 2), report-independent increases were shared by both report and no-report data in signal detection regions, including primary visual cortex (V1), fusiform gyrus (FG), and the posterior middle frontal gyrus (PMFG; near to the frontal eye fields, FEF). Additional report-independent subcortical and cortical increases were found at early times in arousal and salience networks, including the midbrain tegmentum (MT), thalamus (Th), nucleus accumbens (NA), anterior cingulate/supplementary motor area (AC/SMA), and anterior insula/claustrum (AI). Finally, shared report-independent increases at early times were seen in attention and executive control networks, including the anterior inferior parietal lobule (AIPL), dorsal inferior parietal lobule (DIPL), superior parietal lobule (SPL), medial parietal cortex (MP), anterior middle frontal gyrus (AMFG), frontal pole (FP), and cerebellum (Crus I, II).

**Fig. 3.**
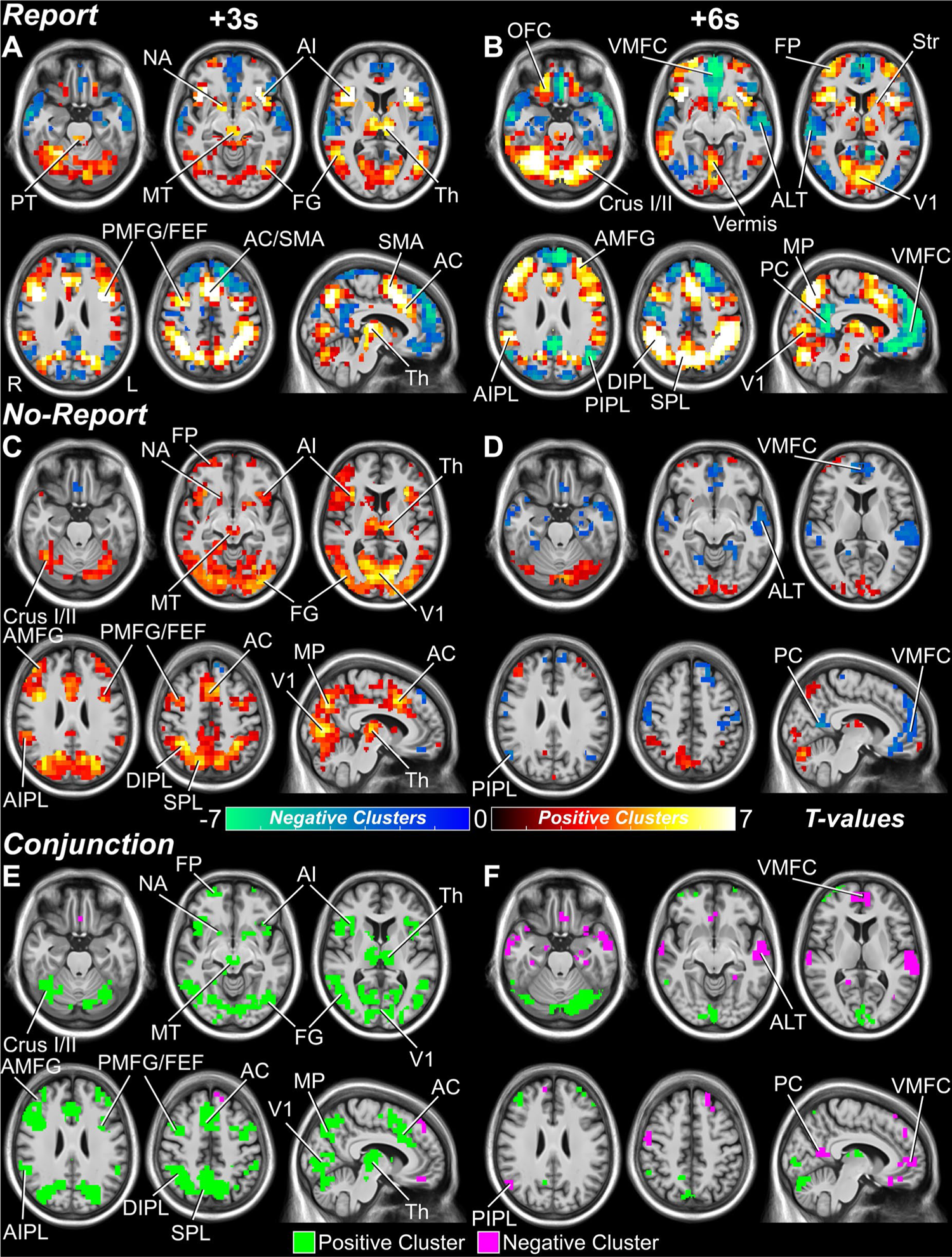
Whole brain fMRI maps for conscious perception with and without overt report. (**A, B**) Report and (**C**, **D**) no-report perceived minus not perceived statistically significant voxels from cluster-based permutation tests (*p* < 0.05) at 3 and 6 seconds post-stimulus presentation (*8*). Statistically significant positive and negative voxel cluster *t*-values are shown in warm and cool colors, respectively. Anatomical regions are labeled at their peak response between the 3 and 6-second time points. (**E**, **F**) Voxel-level conjunction analysis of report and no-report, perceived minus not perceived statistical whole brain maps with shared (report-independent) increases shown in green and shared decreases shown in purple. Pontine tegmentum (PT), midbrain tegmentum (MT), thalamus (Th), nucleus accumbens (NA), striatum (Str), anterior insula/claustrum (AI), anterior cingulate (AC), supplementary motor area (SMA), primary visual cortex (V1), fusiform gyrus (FG), anterior middle frontal gyrus (AMFG), posterior middle frontal gyrus (PMFG), frontal eye fields (FEF), frontal pole (FP), orbital frontal cortex (OFC), ventral medial prefrontal cortex (VMFC), anterior inferior parietal lobule (AIPL), dorsal inferior parietal lobule (DIPL), posterior inferior parietal lobule (PIPL), superior parietal lobule (SPL), medial parietal cortex (MP), posterior cingulate/precuneus (PC), and anterolateral temporal cortex (ALT). See Slides S1, 2, and 4 for all times 20 seconds pre and post-stimulus presentation for report, no-report, and conjunction results. Data for report stimuli are from Report Paradigm (N = 34) and Report + No-Report Paradigm (N = 65); data for no-report stimuli are from Report + No-Report Paradigm (N = 65; one perceived subject data set was excluded because at the prediction score threshold of 0.75 there were no predicted perceived trials; see the *Covert Prediction of Conscious Perception – Machine Learning Pipeline* section in (*8*)).

In contrast, at 6 seconds post-stimulus and later there were fewer shared report-independent regions seen in both report and no-report data (Fig. 3B, D; fig. S15B, D; Slide S1, 2). Most shared changes at 6 seconds involved decreases in default mode network (DMN) areas, including the ventral medial prefrontal cortex (VMFC), posterior cingulate/precuneus (PC), posterior inferior parietal lobule (PIPL), and anterolateral temporal cortex (ALT). Shared increases at 6 seconds were limited to few and relatively small regions of V1, cerebellum, fronto-parietal cortex, and Th. At the same time, the report data alone showed prominent persistent or peak increases at 6 seconds in attention and executive control networks, including AIPL, SPL, MP, AMFG, orbital frontal cortex (OFC), FP, striatum (Str), and cerebellum, as well as, more prominent decreases in DMN regions (Fig. 3B; fig. S15B; Slide S1).

Conjunction analyses emphasize the broad shared report-independent networks seen mainly at earlier times in fMRI including regions important for signal detection, arousal, salience, attention, and executive control, as well as, some involvement of DMN (Fig. 3E, F; Slide S4). Report versus no-report statistical comparisons with cluster-based permutation tests (*p* < 0.05) of perceived minus not perceived fMRI signal show greater differences at later times, including in left cortical motor regions, possibly linked to motor planning for subsequent right-hand behavioral responses (fig. S16A, B; Slide S3). Likewise, analysis of signals significant only in report (fig. S16C, D; Slide S5) highlighted the separate report-dependent regions such as motor cortex, OFC, Str, and PC, not significantly involved at any time in no-report data, as well as, attention, executive, and cerebellar regions that were shared at early times, but remained persistently activated only in the report data. Analysis of signals significant only in no-report data found early increases in sensory regions (e.g., V1 and FG) that are unique to the no-report network (fig. S16E, F; Slide S6).

To further investigate the main large-scale networks involved in conscious perception with and without report and the temporal profile of these networks, we used temporal correlation-based k-means clustering across the entire brain (*8*). Data-driven clustering of statistically significant voxels for report perceived versus not perceived fMRI signals revealed three anatomically and functionally distinct networks: (1) early positive, (2) late positive, and (3) late negative (Fig. 4A-F; fig. S17D, E, F). The early positive network has a peak at 3-4 seconds after stimulus onset and includes subcortical and cortical detection, arousal, and salience networks (DAS). They include FG, PMFG, MT, Th, NA, AC/SMA, AI, cerebellar vermis, and subregions of the Str, AIPL, SPL and MP (Fig. 4A). The late positive network peaks ∼6 seconds after stimulus onset and includes task-positive networks (TPN) such as AMFG, OFC, FP, cerebellum Crus I and II, and subregions of the Str, AIPL, SPL, and MP (Fig. 4B). The late negative network has a trough at 6-8 seconds after stimulus onset and occupies exclusively cortical DMN regions, including the VMFC, PC, PIPL, and ALT (Fig. 4C). Thus, three major and distinct brain networks for conscious perception emerge from the fMRI data entirely based on BOLD timecourse dynamics.

**Fig. 4.**
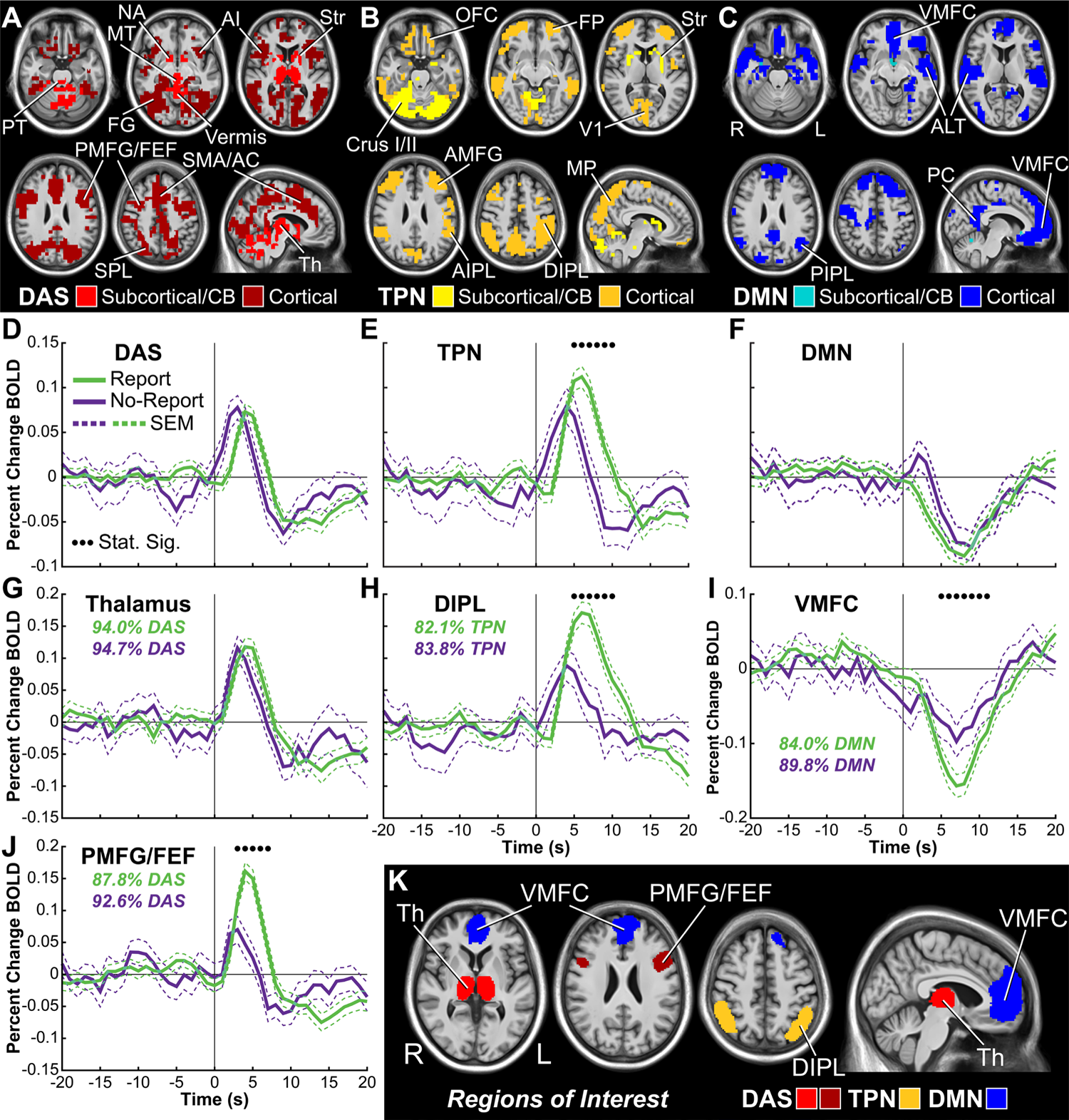
Data-driven anatomical clustering of fMRI signals for conscious perception and region of interest (ROI) timecourses. (**A-C**) Three large-scale networks found with k-means clustering based on fMRI timecourses (fig. S17). (**A**) detection, arousal, and salience networks (DAS), (**B**) task-positive networks (TPN), and (**C**) default mode network (DMN). Cortical and subcortical/cerebellar territories are shown in different shades. (**D-F**) Mean percent change blood-oxygen-level-dependent (BOLD) timecourses for report (green) and no-report (purple) perceived minus not perceived conditions for DAS, TPN, and DMN. (**G-J**) Selected subregion mean percent change BOLD timecourses (see also fig. S18-20). Percentages of voxels from a particular network (DAS, TPN, or DMN) versus all network voxels found in each ROI are shown for report (green) and no-report (purple) data (*8*). (**K**) Anatomical visualization of subregion ROIs from **G**, **H**, **I**, and **J**. Pontine tegmentum (PT), midbrain tegmentum (MT), thalamus (Th), nucleus accumbens (NA), striatum (Str), anterior insula/claustrum (AI), anterior cingulate (AC), supplementary motor area (SMA), primary visual cortex (V1), fusiform gyrus (FG), anterior middle frontal gyrus (AMFG), posterior middle frontal gyrus (PMFG), frontal eye fields (FEF), frontal pole (FP), orbital frontal cortex (OFC), ventral medial prefrontal cortex (VMFC), anterior inferior parietal lobule (AIPL), dorsal inferior parietal lobule (DIPL), posterior inferior parietal lobule (PIPL), superior parietal lobule (SPL), medial parietal cortex (MP), posterior cingulate/precuneus (PC), and anterolateral temporal cortex (ALT). Stimulus onset was at time = 0. Data for report stimuli are from Report Paradigm (N = 34) and Report + No-Report Paradigm (N= 65); data for no-report stimuli are from Report + No-Report Paradigm (N = 65; one perceived subject data set was excluded because at the prediction score threshold of 0.75 there were no predicted perceived trials; see the *Covert Prediction of Conscious Perception – Machine Learning Pipeline* section in (*8*)).

How does overt report change the signal timecourses in these networks? Analysis of mean timecourses for all voxels within each network revealed that only TPN was on the whole different between report and no-report data at later times (Fig. 4D-F). However, subregion analyses showed that all three networks contained both regions that agreed and regions that differed between report and no-report data (Fig. 4G-J; fig. S18-20). Importantly, the fMRI timecourse for Th (>94% DAS voxels) did not differ between report and no-report conditions (Fig. 4G). In contrast, the PMFG/FEF (>87% DAS voxels), DIPL (>82% TPN voxels), and VMFC (>79% DMN voxels) had greater signal amplitude and duration for report versus no-report conditions (Fig. 4H-J). Similar to the conjunction and difference analyses already discussed (Fig. 3E, F; fig. S16), additional timecourse analyses confirmed that some brain areas (e.g., left motor cortex; fig. S19) are only significantly involved in report data, whereas many more regions are shared between report and no-report data at early times, but show persistent or larger signals in report data at later times (e.g., DIPL, SPL, MP; fig. S18-20).

In summary, we found that multiple mechanisms contribute to report-dependent and report-independent conscious visual perception. Report-independent signals are early and transient, and on EEG include the N100 and VAN. Report-independent signals from fMRI with the identical behavioral paradigm and many of the same participants as EEG, showed early and transient, but widespread involvement of three major subcortical and cortical brain networks.

The importance of subcortical networks in conscious perception was further supported by our identification of the TAP through direct recordings from the human thalamus. Meanwhile, report-dependent signals were late and persistent, and on EEG include the P2/N2 and P3/LP; and late (∼6 seconds) fMRI signals in DAS, TPN, and DMN. Persistent BOLD signals for report data may be linked to prolonged neuronal activity changes needed for subsequent report, manifested electrophysiologically as late potentials (fig. S11A), or other sustained activity following perception (*9, 21*).

These findings support an approach to studying consciousness based on investigation of multiple overlapping systems in neuroscience, rather than looking for a single theoretical model. This approach has succeeded in the study of other important and complex biological functions (e.g., reproduction and digestion) where no single model is proposed, and instead multiple contributing processes are found that provide key features. Our identification of specific report-independent subcortical and cortical systems overlapping in space and time is an important step forward towards understanding the neural mechanisms of consciousness. These systems can be placed along a proposed timeline consistent with the three major networks found in our data (*1*). First, (1) *Detection/arousal/salience (DAS)*: when a visual stimulus appears, activation of V1 interacts with FG, FEF, and other regions for signal detection (*25–28*). A dynamic transient pulse in subcortical arousal (e.g., MT and Th) and emotional/motivational systems (e.g., NA) facilitates bottom-up attentional salience and top-down attentional control (*23, 29, 30*). Next, (2) *Default-mode network (DMN)*: switching off the DMN and related circuits can reduce competing signals to prevent interference with conscious perception (*9, 31–33*). Finally, (3) *Task-positive networks (TPN)*: a broad wave of hierarchical processing sweeps through task-positive cortical and subcortical areas to fully process the event before it is encoded in working and episodic memory systems. The later events along this timeline, especially for TPN, merge into report-dependent, post-perceptual processing needed for subsequent report.

Brain mechanisms of conscious visual perception are early, transient, and large-scale. The spatiotemporal signatures of conscious perception suggest that the human brain produces consciousness using synergistic and redundant systems. A large and layered neural architecture would be advantageous by making the conscious network resistant to damage or change.

Investigating these elegant and complex overlapping systems may lead to a satisfactory explanation for consciousness.

## Supporting information

Slide S1

Slide S2

Slide S3

Slide S4

Slide S5

Slide S6

## Acknowledgments

Thank you to the Yale Magnetic Resonance Research Center technicians, Yale-New Haven Hospital Neurology EEG technologist Tamara Wing and University of Pittsburgh Medical Center EEG Laboratory manager Lauren Doak R.EEG.T, CLTM. Thank you to Shweta Majumder, Zailyn Tamayo, and Lining Pan for assistance in preparing source data for public availability.

## Author contributions

Conceptualization: HB

Formal analysis: SK, MA, JD, JR, QX, ZD, JP, HK, AK, SF, DJ, KC, CH, AA, ES, KCL

Funding acquisition: SK, HB

Investigation: SK, MA, JD, SF, KW, KC, ES, JW, KCL, NH, AU, RR, MC

Methodology: SK, MA, JD, ZD, JP, KW, KC, AA, SW, OM, JW, KCL, NH, MM, AU, RC, MP, RR, MC, HB

Project administration: SK, HB

Resources: JW, NH, MM, AU, RC, RR, MC, HB

Supervision: HB

Writing – original draft: SK, HB

Writing – review & editing: SK, MA, JD, JR, QX, JD, JP, HK, AK, SF, DJ, KW, KC, CH, AA, ES, SW, OM, JW, KCL, NH, MM, AU, RC, MP, RR, MC, HB

## Data and materials availability

All data is available in the manuscript or the supplementary materials. Data are available at http://kronemer-blumenfeld-data.yale.edu <<Note: this URL will be open at the time of publication>>

## Supplementary Materials

## Materials and Methods

### Participants

#### Healthy Participants

A total of 144 healthy, adult participants were recruited from Yale University and the New Haven, Connecticut communities. Inclusion criteria were: (1) normal vision with or without correction and (2) normal hearing without an assistive hearing device. Exclusion criteria were: (1) past history or current diagnosis with a psychiatric or neurological disorder, (2) pregnant, claustrophobic, or nonremovable ferrous objects inside or on the body, if participating in the functional magnetic resonance imaging (fMRI) procedure, (3) a hair type or style limiting access to the scalp for electrode placement, if participating in the high-density scalp EEG (hdEEG) procedure, and (4) vision correction that required either hard contact lenses or glasses, if participating in the eye tracking and pupillometry procedure due to distortion of the corneal reflection. While not an exclusion criterion, no healthy participant reported a history of loss of consciousness for more than 5 minutes.

There were four primary data sets gathered from the healthy participants: (1) Report Paradigm (detailed in the *Visual Perception Paradigms* section) with simultaneous fMRI recording (N = 37; mean age = 27.22 years; age range = 18-42 years; females = 17; right-handed = 35), (2) Report Paradigm with simultaneous hdEEG and binocular eye tracking and pupillometry recordings (N = 59; mean age = 26.20 years; age range = 19-43 years; females = 37; right-handed = 53), (3) Report + No-Report Paradigm (detailed in the *Visual Perception Paradigms* section) with simultaneous fMRI and monocular eye tracking and pupillometry recordings (N = 65; mean age = 24.77 years; age range = 18-46 years; females = 39; right-handed = 64), and (4) Report + No-Report Paradigm with simultaneous hdEEG and binocular eye tracking and pupillometry recordings (N = 65; mean age = 24.58 years; age range = 18-46 years; females = 39; right-handed = 63). Summary information for the healthy participant data sets are reported in Table S1.

In the Report Paradigm, of the 37 participants with fMRI and 59 subjects with hdEEG, 16 were common between fMRI and hdEEG data sets. In the Report + No-Report Paradigm, of the 65 participants with fMRI and 65 subjects with hdEEG, 60 were common between fMRI and hdEEG data sets. There were 4 participants in common between the Report and Report + No-Report Paradigm data sets.

#### Patient Participants

Seven adult participants with chronically implanted thalamic depth electrodes for seizure monitoring and treatment were recruited from the University of Pittsburgh Medical Center Department of Neurology Epilepsy Division and the Yale-New Haven Hospital Comprehensive Epilepsy Program. Inclusion criteria were: (1) normal vision with or without correction and (2) normal hearing without an assistive hearing device. Psychiatric or neurological disorder diagnoses were not considered as exclusion criteria for the patient participant group. Two primary data sets were gathered from the patient participants: (1) Report Paradigm with simultaneous low-density scalp EEG (ldEEG) and thalamic intracranial EEG (icEEG) (N = 6; mean age = 24.17 years; age range = 20-31; females = 4; right-handed = 6) and (2) Report Paradigm with icEEG alone, without concurrent ldEEG (N = 1; age = 29; female = 1; left-handed = 1). Summary information for the patient participant data sets is reported in Table S1.

#### Participant Exclusions

Healthy participants were excluded from analyses based on behavioral performance in the Report and Report + No-Report Paradigms, or for excessive movement while participating in the fMRI procedure. The exact parameters for the behavioral and movement-based rejections are described in their respective subsections below (the *Behavioral Exclusions* and *Motion-Based Rejections* sections). In addition, two participants were excluded from the Report Paradigm with the fMRI procedure for falling asleep during the study or incorrect finger position on the task response box. Two participants were excluded from the Report Paradigm with the hdEEG procedure because of either falling asleep during the study or a corrupted data file. Finally, one participant was excluded from the Report + No-Report Paradigm with the fMRI procedure because of being unable to complete the study session due to back spasms during the fMRI sequences. No patient participant was excluded.

In summary, the current investigation comprises two main participant groups, healthy and patient volunteers, from which six primary data sets were gathered: (1) healthy, Report Paradigm with fMRI, (2) healthy, Report Paradigm with hdEEG, eye tracking, and pupillometry, (3) patient, Report Paradigm with ldEEG and icEEG, (4) patient, Report Paradigm with icEEG, (5) healthy, Report + No-Report Paradigm with fMRI, eye tracking, and pupillometry, and (6) healthy, Report + No-Report Paradigm with hdEEG, eye tracking, and pupillometry (Table S1).

All study procedures were carried out in accordance to the Declaration of Helsinki and all participants gave informed consent for participating in the study procedures approved by the Yale University and University of Pittsburgh Institutional Review Boards. For all data gathered during the COVID-19 pandemic, adapted study procedures for reducing the risk of COVID-19 transmission were approved by Yale Environmental Health and Safety, including enhanced disinfection, social distancing, and personal protective equipment protocols.

#### Visual Perception Paradigms

Our goal was to develop a valid approach to investigate brain signals during conscious perception with versus without overt report. Briefly, our overall strategy was as follows: first, we developed a Report Paradigm (*9*); next we developed a Report + No-Report Paradigm and confirmed the report data from both paradigms were similar; finally, we developed a machine learning approach to classify the no-report data in the Report + No-Report Paradigm as perceived or not perceived.

The two visual, perceptual threshold paradigms (Report and Report + No-Report Paradigms) were designed for the analysis of three primary perceptual contrasts: (1) compare report perceived versus not perceived visual stimuli (tested with the Report Paradigm and the report portion of the Report + No-Report Paradigm); (2) compare no-report perceived versus not perceived visual stimuli (tested with the no-report portion of the Report + No-Report Paradigm); and (3) compare (1) versus (2), or results from report versus no-report perceptual tasks.

#### Software and Equipment

The behavioral paradigms were written in Python (www.python.org) and run using the open-source software PsychoPy (www.psychopy.org) loaded on a 15.6 inch display (screen resolution 1280×780 pixels) laptop computer equipped with a NVIDIA GeForce graphics card. The following two laptops were used with similar specifications: (1) MSI Model MS-16H2 running Windows 10 for Report Paradigm with hdEEG, ldEEG, and icEEG, and Report + No-Report Paradigm with hdEEG and fMRI, and (2) Acer Aspire V Series Model ZRQ running Windows 8 for Report Paradigm with fMRI.

In the simultaneous ldEEG and icEEG procedure, the experimental laptop was placed on a table at eye-level and centered 85cm from the participant, measured from the center of the laptop screen to the participants’ nose bridge. In the hdEEG procedure, the participants viewed the task on a 17-inch external LCD monitor (Report Paradigm: Iiyama ProLite E1780SD; Report + No-Report Paradigm: Acer V176L) mounted on a chart-attached arm mount (EyeLink 1000 Plus System, SR Research, Inc.) that mirrored the image from the experimental laptop connected via Mini DisplayPort to VGA adapter. The external monitor was positioned at eye-level and centered 55cm from the center of the screen to the participants’ nose bridge. In the fMRI procedure, participants laid supine and viewed the task via a mirror mounted on top of the head coil that reflected the experimental laptop display image on an MRI-compatible rear-projection screen (Psychology Software Tools, Inc.) that was positioned inside or immediately behind the MRI bore (fig. S2A). The image on the projector screen was shown via a projector placed either in an adjacent room that projected the image through a waveguide into the MRI room (Report Paradigm: Christie LW401; fig. S2A, see i) or inside the magnet room and projected via a mirror onto the projector screen (Report + No-Report Paradigm: Hyperion, Psychology Software Tools, Inc.; fig. S2A, see ii).

All display systems (laptop monitors, external monitors, and projectors) were repeatedly tested for consistent display duration and onset time of target stimuli in the experimental paradigms based on photodiode measurements from the display screens. These results showed that all displays had a target stimulus duration of between 48-51 milliseconds (ms; programmed duration was 50ms) for over 95% of presentations and a variable stimulus onset time accuracy to within approximately one video frame (16.7ms). Details for implementation of the display temporal reliability tests are reported in another publication from our group (*9*). Synchronization of timing between recording systems is described below (the *Behavior and Task Event Synchronization* section).

For all procedures, behavioral responses were recorded with a 1×4 inline button response box connected to the experimental laptop via USB and sampled at 1000Hz (Current Designs, Inc.; hdEEG, ldEEG, and icEEG procedures: Model OTR-1×4-L; fMRI procedure: Model HHSC-1×4-L). Regardless of handedness, participants were instructed to make responses with the button response box using their right hand and with fingers sequentially placed along the four buttons, with the first button pressed with the index finger and the fourth button pressed with the pinky finger.

### Report Paradigm

The Report Paradigm was previously administered and published by our group (*9*). In summary, the paradigm consists of two sequential phases: (1) a perceptual threshold calibration and (2) testing phase. In both task phases, the target stimulus was a greyscale, neutral expression, human face (3.7×4.6 degrees) selected from the FACE database (*34*). The target stimulus appeared for 50ms in one of four pre-selected quadrant locations of the display screen (Fig. 1A). The background of the screen was filled with either a visual static noise or a nature documentary (BBC series Blue Planet episode “Coral Seas”). The documentary background was played with audio, aiming to mimic a naturalistic viewing environment. All participants experienced both background conditions in alternating task runs. The initial background used in a run (static noise or documentary) was counterbalanced across participants. In the documentary background condition, four rectangular static noise patches were shown in each of the quadrants locations of the screen where the target stimulus could appear to control the background image over which the target stimulus was presented. At the center of the screen was a white fixation cross (0.3×0.3 degrees).

#### Calibration Phase

The goal of the calibration phase was to set the perceptual threshold opacity of the target stimulus. Participants were instructed to fixate on the central fixation cross at all times and to immediately respond with either an index or middle finger button press (counterbalanced across participants) each time they saw the target stimulus. During calibration, the target stimulus appeared at a jittered interval of between 1-1.5 seconds. The opacity for any given stimulus was one of 25 pre-defined opacity values ranging between 0.01 to 0.25 (static noise background) or 0.08 to 0.32 (documentary background) with an increment of 0.01 between these opacity ranges, where full opacity was 1.00. Each of the 25 opacity values was selected for 8 stimuli presentations and twice within each of the quadrant locations for a total of 200 stimulus presentations. Background conditions were tested separately with two calibrations phases of approximately 4 minutes each.

The perceptual threshold opacity value (i.e., the opacity value where only 50% of the target stimulus presentations are perceived) was estimated for each participant and background condition by modeling the detection psychometric response across opacity values with a sigmoidal cumulative function. Full details on the perceptual threshold opacity value estimation are provided in our previous report (*9*).

#### Testing Phase

The goal of the Report Paradigm testing phase was to gather instances of report perceived and not perceived stimuli. For the fMRI and hdEEG procedures, the testing phase was broken into approximately 11-minute runs of 32 trials each. For 6 of 7 participants completing the Report Paradigm with the icEEG procedure, the testing phase run duration was defined by the recording duration limits of the NeuroPace RNS® (NeuroPace, Inc.) icEEG recording system (the *Intracranial EEG (icEEG)* section). When recording with this system, whenever the run duration breached either 210 or 450 seconds for the 4 and 8-minute limited recording systems, respectively, the task was instructed to end the run after completion of the current trial. These icEEG participants were also those with simultaneous ldEEG recordings. The remaining icEEG participant recorded with the Natus NeuroWorks, Inc., icEEG system did not present a recording duration limit and the testing phase duration was identical to that used in the healthy participant fMRI and hdEEG procedures.

A single trial comprised of four phases: (1) pre-stimulus, (2) stimulus, (3) post-stimulus, and (4) response phases (Fig. 1A). The fixation cross and background were continuously shown in the pre-stimulus, stimulus, and post-stimulus phases. In the response phase, the fixation cross was removed and the background was replaced with a solid grey image while perception and location questions were displayed. Details of these question for the Report paradigm are in (*9*) and were similar to those used for quadrant relevant stimuli in the Report + No-Report paradigm (fig. S1A). Participants were instructed to maintain fixation while the fixation cross was present, but could break their fixation in the response phase.

The pre-stimulus phase duration was jittered between 6-10 seconds (fMRI, hdEEG, and icEEG-only procedures) or 3-5 seconds (simultaneous ldEEG and icEEG procedure). In the subsequent stimulus presentation phase, the target stimulus appeared in one of the four screen quadrant locations. The stimulus opacity was set to the previously estimated perceptual threshold opacity value. However, in 12.5% of trials, no stimulus appeared during the stimulus presentation phase. These so-called blank trials were later used to assess participant performance and establish a behavioral-based exclusion criterion (see the *Behavioral Exclusions* section). Following stimulus presentation was the post-stimulus phase with a variable latency of either 1 and 15 seconds (fMRI, hdEEG, and icEEG-only procedures) or 3, 4, and 5 seconds (simultaneous ldEEG and icEEG procedure). In subsequent analyses of healthy participant fMRI and hdEEG recordings from the Report Paradigm, only the 15-second post-stimulus interval trials were analyzed (detailed further in the *Functional Magnetic Resonance Imaging (fMRI)* and *Low and High-Density Scalp EEG (ldEEG and hdEEG)* sections). Nevertheless, the 1-second post-stimulus trials were included to affirm that “forgetting” was not contributing substantially to the not perceived trials (*9*). The post-stimulus delay latencies were equally likely and randomly selected on a trial-by-trial basis. Shorter pre and post-stimulus latencies were programmed for the simultaneous ldEEG and icEEG recordings due to a brief study session duration and wanting to maximize the number of completed trials per participant.

Finally, the response phase posed two sequential, self-paced forced-response questions regarding the most recently completed trial. The initial query was a perception question (“Did you see a stimulus?”) with a binary selection of yes or no. The index or middle finger buttons on the response box corresponding for either response to the perception question was counterbalanced across participants. Regardless of the response to the perception question, a location question followed (“Where was the stimulus located?”) with the numbers “1”, “2”, “3”, and “4” shown in each of the quadrants of the screen (fig. S1A). Each of the numbered quadrant locations were paired to the four response box buttons and these pairings were made constant across participants. Participants were instructed that if they saw the target stimulus in the current trial they should select the correct quadrant where it appeared. However, if they did not see the stimulus in the current trial they should randomly guess one of the quadrant locations. Upon responding to the location question, either a new trial began from the pre-stimulus phase or, if the final trial in a run, a break screen was presented (“Great job! Take a break.”) and the experimenter communicated with the participant on when they were prepared to commence the subsequent run or to complete the study session.

After each run, the stimulus detection rate was calculated from the responses to the perception question to assess if perceptual thresholding of the target stimulus was achieved. If perception rates fell above or below the target 50% detection rate by more than 3.5% (i.e., a perception rate percentage above 53.5% or below 46.5%), a new opacity value was estimated to achieve perceptual threshold and this opacity value was used in the subsequent run. These run-by-run opacity adjustments were necessary to track the intrinsic fluctuations in perceptual behavior by changes in participant motivation, attention, and arousal during the study session. This procedure ensured that the mean opacity values for perceived and not perceived stimuli remained within 5-10% of each other across subjects. Details for calculating the run-by-run adjustments to the perceptual threshold stimulus opacity are previously published (*9*).

### Report + No-Report Paradigm

The Report + No-Report Paradigm was a modified version of the Report Paradigm with an embedded no-report condition. The critical adaptation for the Report + No-Report Paradigm was the addition of four central stimulus locations: above, below, left, and right of the fixation cross (Fig. 1B). Thus, the Report + No-Report Paradigm included 8 non-overlapping stimulus presentation sites between two stimulus location sets: (1) four quadrant locations and (2) four central locations. These location sets defined either report (task-relevant) or no-report (task-irrelevant) stimuli, detailed below. Moreover, the Report + No-Report Paradigm omitted the documentary background condition and only displayed the static noise background condition from the Report Paradigm because it was previously found that results did not differ between these two backgrounds (*9*). And, just as in the Report Paradigm, the target stimulus was the same greyscale face that appeared for 50ms and consisted of two sequential phases: (1) a perceptual threshold calibration and (2) testing phase.

### Calibration Phase

The aim of the calibration phase was to estimate the perceptual threshold opacity value for the target stimulus in both the quadrant and central location sets. The perceptual threshold opacity was independently defined for each location set due to differential visibility of the target stimulus nearer to the center of vision (the center locations) versus the periphery (the quadrant locations). Participants were instructed to maintain fixation to the central fixation cross at all times during the calibration phase and to immediately respond with a button press either with the index or middle finger (counterbalanced across participants) whenever they saw a target stimulus appear in any location on screen. Stimuli appeared at a jittered interval of between 1-1.5 seconds and the stimulus location for any one presentation was randomly selected among the 8 possible locations between the two location sets. The opacity values for all faces that appeared during the calibration phase were set to one of 25 pre-defined opacity values ranging between 0.01 to 0.25, incremented 0.1 over this range. Each opacity value was shown in each of the 8 locations and twice within each location. Thus, 200 stimuli per location set (quadrant and center), or 400 stimuli total were presented during the calibration phase.

Just as in the Report Paradigm, the stimulus perception responses across opacity values within each location set was modeled with a sigmoidal cumulative function from which the perceptual threshold opacity values for the centrally and peripherally located stimuli were estimated independently. The estimated perceptual threshold opacity values were used in the subsequent run of the Report + No-Report Paradigm testing phase.

### Testing Phase

The testing phase consisted of approximately 11-minute runs of 24 trials each. A trial included six sequential phases: (1) pre-stimulus, (2) first stimulus, (3) inter-stimulus, (4) second stimulus, (5) post-stimulus, and (6) response phase (Fig. 1B; fig. S1A, B). The static noise background appeared continuous from the onset of the pre-stimulus phase through the post-stimulus phase and was then replaced with a solid grey background in the response phase. The target stimulus could appear among one of three opacity conditions: (1) no stimulus or blank (12.5%), (2) perceptual threshold (75%), and (3) fully opaque (12.5%). The addition of fully opaque stimuli allowed measurement of participant false negative detection rate. In a single trial, there could be a minimum of no target stimulus presentations (first and second stimulus phases showed blank opacity target stimuli) and a maximum of two target stimulus presentations (first and second stimulus phases showed either threshold or opaque opacity target stimuli). The frequency of the target stimulus appearance in any one of the 8 possible stimulus locations was proportional and randomly selected for each presentation. However, the stimulus could only appear once per trial within each location set. Therefore, the first stimulus was equally likely to appear in either the center or quadrant location sets, while the second stimulus was required to appear in whichever location set was not selected for the first stimulus in each trial.

The center and quadrant location sets defined task-relevant and irrelevant stimulus conditions during the testing phase. In each testing session, participants were instructed that one location set was task-relevant while the other location set was task-irrelevant and testing sessions with different location instructions were conducted on different days. For example, on a day where the center stimuli were task-relevant, they were task-relevant for all trials in the session on that day. For the stimuli that appeared in the task-relevant location set, participants were asked to recall and report via button presses on the perception of these stimuli in the trial response phase (fig. S1A, B). Meanwhile, for stimuli that appeared in the task-irrelevant location set, participants were instructed that they would not be required to remember or respond to these stimuli. In other words, the trial response phase only inquired on the task-relevant stimuli. Therefore, the task-relevant stimuli represented a reported perceived and not perceived stimulus condition, identical to those of the Report Paradigm, while the task-irrelevant stimuli represented a no-report perceived and not perceived stimulus condition.

Participants were recruited to complete the Report + No-Report Paradigm with both task-relevant location set conditions in separate study sessions, conducted on different days. Both task-relevant location set conditions were completed for each participant over interleaved hdEEG and fMRI study sessions such that the hdEEG and fMRI data sets for the first task-relevant location set condition was acquired prior to gathering the hdEEG and fMRI data sets for the alternative task-relevant location set condition to minimize contamination by opposing task relevancy rules from previous study sessions. The initial task-relevant location set condition and initial study session modality (hdEEG or fMRI) was counterbalanced across participants resulting in a total of four possible study schedules with the following initial study sessions: (1) quadrant location set task-relevant and fMRI, (2) quadrant location set task-relevant and hdEEG, (3) center location set task-relevant and fMRI, (4), center location set task-relevant and hdEEG (fig. S1C). While participants were recruited to complete all 4 study sessions over an equal number of days, a total of 14 participants did not complete all study sessions because of being lost to follow-up or due to interruption in data gathering by COVID-19 restrictions.

The pre-stimulus, inter-stimulus, and post-stimulus trial phases were jittered intervals between 6-10 seconds during which participants were instructed to maintain fixation at all times (Fig. 1B). The first and second stimulus phases consisted of a 50ms stimulus presentation, unless there was a blank presentation when no stimulus appeared. Finally, the response phase was self-paced and presented two sequential questions for stimuli in the task-relevant location set. First, the perception question appeared (“Did you see a stimulus in a corner?” or “Did you see a stimulus near the center?” for the task-relevant quadrant and center location set conditions, respectively; fig. S1A, B). The participants were instructed to respond with either an index and middle finger button press (counterbalanced across participants) corresponding with either a yes or no response. Next, the location question appeared (“Where was the stimulus located?”) with the numbers “1”, “2”, “3”, and “4” shown in the four locations corresponding to the task-relevant location set (fig. S1A, B). The participant were instructed to respond with one of the four response box buttons corresponding with each of the numbered, task-relevant locations either the correct location where they had noticed a task-relevant stimulus in the current trial or to randomly guess a task-relevant stimulus location if they did not notice a stimulus in the current trial. Upon responding to the location question, the subsequent trial would begin or a post-run break screen would appear (“Great job! End of run, take a break.”). Participants did not provide overt report for task-irrelevant stimuli during the testing phase.

Run-by-run adjustments of the perceptual threshold opacity for the task-relevant stimulus was implemented exactly as detailed for the Report Paradigm and as previously published (*9*).

However, because no perceptual reports were provided by participants for the task-irrelevant stimuli, the detection rate was unknown for these stimuli that form the basis for justifying adjusting the stimuli opacity to better match perceptual threshold performance. Instead, on the premise that underlying fluctuations in motivation, attention, and arousal can alter perception rate uniformly for task-relevant and irrelevant stimuli, the run-based opacity adjustments implemented on the task-relevant stimulus were used to calculate run-based opacity adjustments for the task-irrelevant stimuli. Because task-relevant and irrelevant location sets may have different perceptual threshold opacities, the same multiplicative ratio for the run-by-run opacity adjustment was applied for both task-relevant and irrelevant stimulus using the following formula:

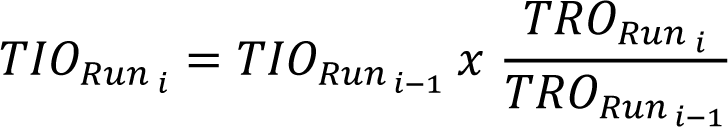

where *TIO* is the task-irrelevant perceptual threshold opacity, *TRO* is the task-relevant perceptual threshold opacity, *Runi* is the current testing phase run, and *Runi-1* is the previous testing phase run.

After completing all runs of the testing phase, participants were administered a free answer questionnaire that inquired on the general experiences during the study session, including whether if at any time during the testing phase the participants perceived stimuli in the task-irrelevant location set and how the opacity of these stimuli compared to the task-relevant stimuli. The goal of this questionnaire was to offer a coarse assessment of what participants perceived during the testing phase, particularly for the task-irrelevant stimuli, which would later be used as the basis for a behavioral exclusion criterion for the Report + No-Report Paradigm, although this criterion resulted in no participant exclusions.

### Behavioral Exclusions

Two report-based behavioral exclusion criteria were applied on the healthy participant data sets from the Report and Report + No-Report Paradigms: (1) if the location accuracy fell below 75% for stimuli that were reported as seen, or (2) if the perception rate of blank (no stimulus) trials exceed 30%. Individual participant behavioral performance was measured and exclusions implemented on the study session-level. For the Report + No-Report Paradigm, only the stimuli at perceptual threshold (Fig. 1B) were considered when calculating the location accuracy percentage for seen stimuli (100 × [seen stimuli with correct locations/all seen stimuli]). For participants who completed only one study session, as for the Report Paradigm with hdEEG and fMRI data sets, session-level behavioral rejections functioned as participant-level rejections.

Meanwhile, for participants who completed multiple study sessions, as in the Report + No-Report Paradigm with hdEEG and fMRI data sets, all remaining participant sessions after behavior-based rejections were included for subsequent analyses.

From these behavioral exclusion criteria, no participants were rejected from the Report Paradigm data set. Meanwhile, a total of four participant study sessions were rejected from the Report + No-Report Paradigm. Behavioral performance exclusions were not applied on the patient participants N = 7) due to low sample size. However, only one patient participant would be eligible for removal by these behavioral exclusions criteria by falling below the seen stimuli location accuracy percentage (71.72% location accuracy percentage of reported seen stimuli trials).

Neither location accuracy nor detection rate based rejections could be applied to the no-report (task-irrelevant) stimuli in the Report + No-Report Paradigm because responses for these events were not gathered. Instead, if participants reported not noticing any task-irrelevant stimuli in the post-testing phase questionnaire administered at the end of each study session this would be interpreted as the task-irrelevant stimuli opacity being sub-perceptual threshold throughout the entire testing phase, thwarting the goal of creating instances of both no-report perceived perceptual events. Therefore, study sessions from the Report + No-Report Paradigm were rejected from analyses if participants reported not seeing any task-irrelevant stimuli during the testing phase. However, ultimately no session rejections were made based on the recall of task-irrelevant stimuli because for all study sessions participants indicated seeing non-opaque, task-irrelevant stimuli during the task testing phase. For example, if a subject was instructed that center stimuli were relevant for a session, they reported also seeing stimuli in the corners (quadrants) that were not fully opaque (i.e., the threshold stimuli) during that session.

### Behavior and Task Event Synchronization

The Report and Report + No-Report Paradigms were administrated with several combinations of simultaneous fMRI, hdEEG, ldEEG, icEEG, eye tracking, and pupillometry. Each combination of behavioral testing and physiological recordings required a unique approach for synchronizing behavioral and task events with ongoing physiological signals. The wiring diagrams for each testing setup is provided in fig. S2 as a supplement to the written descriptions below.

#### fMRI

The experimental laptop received inputs from participant button presses while inside the magnet and the onset of each fMRI volume acquisition (i.e., repetition times) via a USB connection to an interface device (932 Interface, Model FIU-932, Current Designs, Inc.). The input settings of the interface device were adjusted so that button presses were received as the numbers “1”, “2”, “3”, and “4”, each value corresponding with one button on the 1×4 response box. The number “5” was dedicated to signify fMRI volume acquisition events. The first volume acquisition event of each fMRI sequence initiated the behavioral task. All button press and fMRI events were recorded on the experimental laptop by their numerical value in a behavioral log file documenting the occurrence and timing of all salient task events. In subsequent analyses, fMRI signals were related to task and behavioral events by matching these events to their nearest acquired volumes. See fig. S2A.

#### hdEEG

Salient task and behavioral events were related to hdEEG recordings by digital inputs from the experimental laptop recorded in the same time basis as the physiological signals. Four event-linked digital inputs types were marked: (1) trial onset, (2) stimulus presentation, (3) question presentation, and (4) participant button presses. Digital inputs were delivered by the experimental laptop signaling an Arduino Uno R3 board (www.ardunio.cc) via USB to deliver a transistor-transistor logic (TTL) pulse via a DB9 connection to the digital input port of the Net Amps 200 128-channel EEG Amplifier (Model No. C-AMP-128-2000-001; Magstim Electrical Geodesics, Inc.). Each of the four recorded event types were designated a separate paired Arduino Uno channel and pin of the DB9 digital input port. Therefore, the occurrence of a salient task or button press event would initiate the delivery of a TTL pulse from the corresponding Arduino Uno channel that would then be received by the paired pin of the digital input port of the hdEEG amplifier and be marked by the hdEEG recording software (NetStation, Magstim Electrical Geodesics, Inc.) as an event flag with a preprogrammed initial (e.g., “Trl” = trial onset) to distinguish digital event marker types in post-processing. See fig. S2B.

The temporal precision between the TTL pulse delivery and on-screen stimulus presentation was tested, as previously published (*9*). A consistent latency of approximately two to three video frames (16.7ms per frame) was found between the TTL pulse onset time and stimulus presentation onset time. The mean TTL-stimulus presentation latency of 42ms was corrected in preprocessing analyses.

#### ldEEG

TTL pulses of varied duration were used to mark salient task events on the same time basis as ldEEG recordings. Four task event types were synchronized to the ldEEG recordings: (1) trial onset, (2) stimulus presentation, (3) question presentation, and (4) participant button presses. The occurrence of one of these events initiated the delivery of a TTL pulse through an electrical safety transducer commanded by the experimental laptop via an Arduino Uno R3 board (www.arduino.cc) and fed into empty channels (CHIN1 and CHIN2) of the ldEEG breakout box (XLTEK Sleep/EEG Breakout, Natus, Inc.). The TTL pulses appeared in a channel of the ldEEG montage in the same time basis as the on-going ldEEG recordings. The TTL pulse duration coded the event type allowing for event type discrimination in post-processing (TTL pulse durations by salient task event: trial onset = 250ms, stimulus presentation = 150ms, question presentation = 100ms, and button presses = 50ms). See fig. S2C.

Again, the latency between the TTL pulse onset time and stimulus presentation were tested as previously detailed (*9*) and found to have a reliable latency of on average 42ms that was corrected in preprocessing analyses.

#### icEEG – NeuroPace

The synchronization of task events with the RNS System (NeuroPace, Inc.) icEEG recordings was achieved by inducing transient artifacts in the icEEG signals that could be reliably distinguished from background noise and relevant physiology in post-processing. The procedure for inducing event markings in the RNS recordings has been previously published (*35*). In summary, the event marking procedure requires usage of a research tool hardware (Programmer Tool and Wand Tool; fig. S2C) that interfaces with the implanted Neurostimulator device via the Wand and RNS® Tablet Programmer (Programmer). This implementation allows the experiment to be conducted with no changes to the FDA regulated product. An electrical pulse commanded by the experimental laptop and delivered via the coil antenna of the Wand Tool interrupts the telemetry between the Wand (connected to the Programmer by USB) and Neurostimulator embedded in the skull that stores the icEEG recordings, thus inducing a reproducible artifact in the real time icEEG recordings that appears as a large downward deflection of three brief nadirs of two samples each. The delivery of these synchronization artifacts was initiated by the experimental laptop that commanded the Programmer Tool via micro-USB that subsequently communicates with the Wand Tool via a WA22 4-pin connector.

Unlike other behavioral synchronization procedures that utilize a dedicated event channel to mark salient task events (e.g., a separate empty or digital input channel), the current approach applies event-time artifactual marks directly in all icEEG channels, corrupting the signal at the moment of the event-synchronization mark. Therefore, event markings were only applied prior to trial onset, so as not to overlap with in-trial events of interest. To prevent the possibility of missed synchronization signals due to dropped marks, each trial onset was designated by a sequence of 3 or 4 artifactual marks. In-trial task event times were calculated by adding the time between the initial trial onset mark to the true trial onset time and then adding the known intervals from trial onset to the in-trial event (e.g., stimulus presentation time corrected for the previously described latency between the TTL pulse onset and on-screen stimulus presentation tested with a photodiode), as recorded in the behavioral task log file. The task code was programmed with a 0.5 second delay from the initial event mark to trial onset. An additional delay of approximately 10ms was found between the receipt of TTL pulses command by the experimental laptop from an Arduino Uno R3 (as used in ldEEG event synchronization) and the Wand Tool, tested by recording the outgoing Arduino Uno R3 and Wand Tool signals (National Instruments 12-bit 10kS/s DAQ, Model USB-6008) and comparing their latency. The duration between the electrical pulse to the induction of the artifactual event marks in the icEEG recording was not tested but assumed to be practically instantaneous. Therefore, the total delay from the initial icEEG recording mark to trial onset was estimated to be 510ms and, by using the behavioral trial event-time intervals, the temporal location of in-trial task events of interest were calculated with high temporal precision.

#### icEEG – Natus NeuroWorks

Task and behavioral events were synchronized with the Natus NeuroWorks (Natus, Inc.) icEEG recordings using the same approach detailed for ldEEG and as described previously (*9*). In summary, TTL pulses of variable length coding the event type were delivered via an electrical safety transducer into an empty port of the Natus breakout box that was designated as a research event channel in the icEEG recording montage. As with the ldEEG and hdEEG recordings, the testing of latency between TTL pulse deliver and on-screen stimulus presentation found a consistent latency of on average 42ms that was corrected in post-processing.

#### Eye Tracking and Pupillometry – EyeLink

Synchronization of eye tracking and pupillometry recordings made with EyeLink (SR Research, Inc.) and behavioral events was achieved by digital messages sent by ethernet between the experimental laptop and the EyeLink Host PC (fig. S2A, B). These digital messages reported both the event times (e.g., stimulus presentation, button presses, and response presentations) and details about the trial and target stimuli (e.g., stimulus opacity, presentation location, and task-relevant condition) that could be used independently of the behavioral log file to organize behavioral events in the EyeLink recordings.

### Testing Facilities

All testing rooms were indoor, temperature-controlled facilities at Yale University, Yale-New Haven Hospital, and the University of Pittsburgh Medical Center. The ambient light of the testing rooms was kept consistent by maintaining the identical configuration of indoor light sources across each testing session. In the hdEEG procedure, the testing room lighting was periodically monitored for consistency with a digital light meter (Model LX1330B; Dr.meter, Inc.). In the ldEEG and icEEG procedures, the outdoor light from a window was blocked by a heavy window blind. There were no outdoor light sources in the testing rooms for the hdEEG or fMRI procedures.

### Functional Magnetic Resonance Imaging (fMRI)

#### Equipment and Software

fMRI measurements were acquired with 3 Tesla Siemens Magnetom scanners (Siemens, Inc.) and either a 32-channel or 64-channel head coil at the Yale Magnetic Resonance Research Center (Report Paradigm: Magnetom Trio and 32-channel head coil; Report + No-Report Paradigm: Magnetom Prisma and 64-channel head coil). A high-resolution T1-weighted, whole brain 3D structural image was acquired for each participant at each study session with a magnetization-prepared rapid gradient-epoch sequence (repetition time (TR) = 2010ms; echo time (TE) = 2.81mm; flip angle = 9 degrees; field of view (FOV) = 256×256mm; spatial resolution = 1mm^3^; number of slices = 176). The blood-oxygen-dependent-level (BOLD) fMRI volumes were acquired with a multiband echo-planar imaging sequence (TR = 1000ms; TE = 30.00mm; flip angle = 60 degrees; FOV = 220×220mm; spatial resolution = 2mm^3^; number of slices = 60). For the calibration phase of the Report and Report + No-Report Paradigms, each fMRI run lasted 270 and 600 seconds, respectively, with a corresponding total of 270 an 600 volumes of data acquired. For the testing phase of the Report and Report + No-Report Paradigms, each fMRI run lasted 720 and 700 seconds, respectively, with a corresponding total of 720 and 700 volumes of data acquired.

#### Overall Data Analysis

All fMRI analyses were completed in MATLAB (Mathworks, Inc.) using custom functions and those available through the open-source neuroimaging analysis package Statistical Parametric Mapping (SPM12; https://www.fil.ion.ucl.ac.uk/spm/software/spm12) and the SPM extension toolbox MarsBaR (http://marsbar.sourceforge.net) (*36*). Initial fMRI processing is described here and final analyses are described in the *Statistical Analyses* section.

#### Data Preprocessing and Artifact Rejection

Standard fMRI data preprocessing was applied, including motion correction, nonlinear spatial normalization to the standard Montreal Neurological Institute (MNI) brain template space, and spatial smoothing using Gaussian kernel (FWHM = 6mm). To improve functional image spatial alignment to the MNI brain template space, the acquired MPRAGE for each participant at each study session was manually reoriented to the anterior commissure, coregistered to the mean functional image of each acquired BOLD sequence, and the transformation matrix of warping the MPRAGE to MNI space was applied to the functional images. A trained experimenter reviewed the registration of the preprocessed functional images for optimal alignment to MNI space. When BOLD volumes and the MNI brain template spatial alignment were found unsatisfactory, non-brain tissues were excluded from the MPRAGE image by whole-brain white and grey matter segmentation prior to warping to the MNI brain template. In all cases, the additional step of whole-brain segmentation achieved improved BOLD volume normalization.

Preprocessed BOLD volumes were subsequently passed through a 5-staged denoising procedure previously published from our group (*37*). In sequence, (1) volumes were grey matter masked (i.e., excluding non-grey matter voxels), (2) application of 128Hz high-pass filter, (3) removal of head motion-related signals by regression utilizing a general linear model with the six rigid-body motion parameters estimated during preprocessing image realignment, (4) rejection of voxels at timepoints with a BOLD signal volume-to-volume root mean squared difference that exceeded 5 (DVARS) (*38*), and (5) rejection of voxels at timepoints that exceeded a framewise displacement (FD) threshold of 0.3 (FD was calculated as the sum of the absolute values of change in head movement among the six rigid-body motion parameters) (*38*). Rejected voxels and timepoints were replaced with the MATLAB designation of not-a-number or “NaN”. The denoised fMRI data for each run was converted to percent change relative to the mean BOLD signal of each voxel across the entire BOLD sequence. Finally, behavioral event epochs (e.g., around stimulus presentation) were cropped for analysis from the percent change BOLD signal. Only the 15-second post-stimulus interval trials from the Report Paradigm were considered because the 1-second post-stimulus interval trials were too short to isolate the hemodynamic responses from stimulus presentation versus those of the subsequent trial response period. Behavioral events and BOLD volumes were related by finding the volumes that immediately preceded stimulus presentation, designating this volume as the event-onset volume, and then cropping out event-onset volume centered epochs (total epoch duration = 41 seconds; 20 seconds pre and post the event-onset volume). In the Report + No-Report Paradigm where pupillometry and eye tracking were acquired simultaneously with fMRI, all trials with a blink event at the time of stimulus 28 presentation (50ms stimulus duration window) were rejected from subsequent analyses (see *Eye Tracking and Pupillometry – EyeLink* section for additional details).

#### Motion-Based Rejections

To limit the influence of motion induced artifacts, Report and Report + No-Report Paradigms testing phase runs and corresponding fMRI sequences were rejected from analyses if head motion exceeded 2mm (the equivalent of 1 voxel) in x, y, or z axes, or greater than one degree of rotational motion in the pitch, row, or yaw orientations. This resulted in a total of 74 rejected runs between the Report (12 runs rejected) and Report + No-Report Paradigms (62 runs rejected). The entire fMRI study session for 4 participants in the Report + No-Report Paradigm was rejected because all study session runs were removed by the head motion rejection criteria. One participant who completed the Report Paradigm with fMRI had all but one run rejected by excessive head motion and the remaining run did not have any instance of a perceived stimulus trial. Due to the necessity by subsequent statistically analyses for each participant to have instances of both perceived and not perceived trials, this participant was rejected from fMRI statistical analyses.

### Low and High-Density Scalp EEG (ldEEG, hdEEG)

#### Equipment and Software

Non-invasive hdEEG data were collected with 257 Ag/AgCl electrodes embedded into an elastic net (Hydrocel GSN 256, Magstim Electrical Geodesics, Inc.) and recorded on a desktop computer (Power Mac G5 Quad; Mac OS X v10.5.8, Apple, Inc.) running NetStation version 4.2.2 (Magstim Electrical Geodesics, Inc.). Electrode impedance values were maintained below 50kΩ with conductance gel (Signagel Electrode Gel, Parker Laboratories, Inc.). The EEG signal was digitized at 1000Hz, amplified with two 128-channel amplifiers (Magstim Electrical Geodesics, Inc.) and high and low-pass hardware filtered at 0.1 and 400Hz, respectively. Signals were acquired as Cz-referenced.

Non-invasive ldEEG were collected via 20 Ag/AgCl electrodes (Grass, Natus, Inc.) that were pasted to the participants’ scalp (Ten20 Conductive, Weaver and Company, Inc.) and arranged in the 10-20 system. Electrode leads were passed by touchproof connections into a 32-channel breakout box (XLTEK Sleep/EEG Breakout, Natus, Inc.) and recorded on clinical mobile desktop station running Natus NeuroWorks 9.2.1 Build 5186 (Natus, Inc.) Electrophysiology was sampled at 256Hz and low-pass hardware filtered at 128Hz. Recordings were acquired referenced to an electrode placed on the left hemisphere of the scalp in the middle of the Fz, Cz, C3, and F3 contacts and re-referenced during processing as described below.

#### Data Preprocessing and Artifact Rejection

hdEEG and ldEEG preprocessing was implemented for each study session independently in MATLAB with an in-house, semi-automated data processing pipeline utilizing custom functions and those available by the open-source EEG processing toolbox EEGLAB (*39*). EEG preprocessing consisted of two stages: (1) session-level and (2) epoch-level preprocessing. The specific steps and sequence of the preprocessing pipeline were selected from the most commonly used approaches for EEG preprocessing (e.g., see EEGLAB preprocessing and artifact rejection documentation: https://eeglab.org/tutorials/).

In the session-level stage, data were first 1Hz high-pass filtered (EEGLAB *clean_drifts* function with the transition bands of 0.25 and 0.75Hz). Next, line noise was rejected using the Cleanline procedure (github.com/sccn/cleanline; EEGLAB *pop_cleanline* function with the specified lines frequencies to remove of 60 and 120Hz). Noisy channels were found (EEGLAB *clean_channels* function with the parameters 0.8, 0.5, and 4) and noisy samples were found (EEGLAB *clean_windows* function with the parameters -Inf, 7, and 0.25). The original data for the identified noisy channels were rejected and restored with spherical interpolation (EEGLAB *pop_interp* function). Finally, the hdEEG data was re-referenced to the common average reference. Meanwhile, the ldEEG data was re-referenced to the average of the mastoid electrodes (A1 and A2).

In the epoch-level preprocessing stage, stimulus presentation epochs of 4001ms duration (2000ms before and 2000ms after each stimulus) were cropped for analysis from the preprocessed session data, concatenated, and passed through, first, a 10-component principal component analysis (PCA) and, second, an independent component analysis (ICA) applied on the PCA decomposed data (EEGLAB *pop_runica* function utilizing the infomax algorithm for ICA decomposition). Dimension reduction by PCA was implemented before ICA to reduce the computational time and the amount of data required for robust ICA estimations. A trained experimenter identified and removed ICA components dominated by blinks, eye movements, or heartbeat determined by the inspection of the component scalp topography. Finally, epochs were rejected from subsequent analyses according to two criteria: (1) epochs with more than 25% of samples between 200ms pre-stimulus and 500ms post-stimulus that were found bad in the preceding session-level noisy sample designation stage (i.e., via the EEGLAB *clean_windows* function), and (2) epochs with any instance of a blink between 200ms pre-stimulus and 500ms post-stimulus. For the hdEEG data set, blink occurrence during the critical epoch window were determined from the simultaneous eye tracking recordings with EyeLink. One testing phase run for one participant in the hdEEG Report + No-Report Paradigm condition was rejected from hdEEG analysis because the EyeLink data was corrupted during this run and could not be used to identify blink events for trials acquired in this run.

For the ldEEG recordings, blink occurrences during the critical epoch window were found by an in-house outlier detection algorithm tailored for detecting blinks from the anterior scalp electrodes (FP1, FP2, and FPz). First, the electrophysiology recorded from the selected anterior scalp electrodes were smoothed to emphasize low frequency dynamics (MATLAB *movmean* function; bin size = 10 samples; increments = 1 sample). Next, outlier samples from all 2000ms pre and post-stimulus presentation times were found with the MATLAB *isoutlier* function using the median method (documentation: https://www.mathworks.com/help/matlab/ref/isoutlier.html). Of all the outlier samples identified, samples that exceeded 50µV were maintained for subsequent blink identification. Finally, the remaining outlier samples were identified as blinks if they met two criteria: (1) instances of two or more consecutive outlier samples and (2) agreement of two or more consecutive outlier samples for at least two of the three queried scalp EEG channels. In the case of disagreement among the three channels (i.e., when outlier samples were identified but no two channels with at least two consecutive overlapping outlier samples), a trained experimenter inspected the trial to make a final judgment of whether these outliers exhibited the stereotyped profile of electrophysiological potentials derived from blink.

Corresponding to the hdEEG data set, all ldEEG trials that were found to have a blink during the critical window (between 200ms pre-stimulus and 500ms post-stimulus) were excluded from subsequent analysis.

Finally, like in the Report Paradigm fMRI data set, the 1-second post-stimulus interval trials were excluded due to possible signal contamination from the trial response period 1 second post-stimulus presentation, and only the 15-second post-stimulus interval trials were included.

Analogous trial rejections were not required for the Report Paradigm ldEEG and Report + No-Report Paradigm hdEEG data sets because these paradigms were adapted to have minimum post-stimulus intervals of 3 and 6s, respectively (Fig. 1A, B). Analyses including the 1-second post-stimulus interval trials (representing approximately 20% of all trials from the Report and Report + No-Report Paradigms) were conducted and found without meaningful difference from analyses excluding these trials (results not shown).

### Intracranial EEG (icEEG)

#### Equipment and Software

Two recordings systems were used to acquire icEEG signals from 7 adult, patient participants reported in the current investigation: (1) RNS System (NeuroPace, Inc.; N = 6), and (2) Natus NeuroWorks Quantum (Natus, Inc.; N = 1). See Table S2 for individual patient participant details.

The NeuroPace RNS System is FDA-approved as an adjunctive therapy of drug-resistant focal epilepsy in adults. The RNS System consists of three primary components: (1) depth or cortical strip leads placed at the foci of patient seizure sites, (2) a battery-powered Neurostimulator encasement implanted in the cranium, and (3) a Programmer (tablet or laptop) that communicates with the Neurostimulator via telemetry with a wand attachment held in close proximity to the neurostimulator (fig. S2C). Two generations of RNS Neurostimulator were recorded from in the current investigation: the RNS-300M (N = 1) and RNS-320 (N = 5). The experimentally relevant difference between these RNS Neurostimulator models was that the RNS-300M allows for 4 minutes and the RNS-320 unlimited continuous, real time icEEG recording. As discussed in the *Report Paradigm* methods section, the task run phase duration was modified to accommodate the 4-minute icEEG recording limit imposed by the RNS-300M and was limited to 8 minutes by the experimenter for the RNS-320 system. The RNS System real time icEEG recordings are stored in and obtained from the NeuroPace Patient Data Management System.

The RNS System is capable of targeting numerous cortical and subcortical foci via a maximum of two, 4-contact leads (DL-330-3.5, NeuroPace, Inc), bipolar referenced to its adjacent contact along the lead shaft providing a total of up to 4 icEEG recording channels. We recruited participants with depth contacts targeting the centromedian nucleus (CM) of the intralaminar thalamus because of its known role along with adjacent nuclei, including the central lateral nucleus (CL) in arousal and awareness via broad cortical connectivity, therefore, a possible subcortical candidate of the core consciousness network (*22, 29, 40, 41*). Accordingly, all but one recruited RNS System patient participant were implanted with bilateral depth leads targeting the CM (Table S2). One RNS System participant had one unilateral depth lead targeting the CM and a second lead placed in the white matter of neocortex. Contact/channel localization (see the *Depth Contact and Channel Localization* section) confirms that many channels were in the CM or intralaminar thalamus, or immediately lateral to these sites (fig. S12).

RNS System icEEG recordings were sampled at 250Hz and high and low-pass hardware filtered according to parameters set by the clinician and NeuroPace technician to optimize detection of epileptiform discharges and seizure (Table S2). The RNS System is constrained to collect two channels of data per four contact lead. Therefore, we used the four thalamic contacts (note that one participant had only two thalamic contacts) on each lead to collect two channels of differential icEEG voltage data per lead as follows: (1) contact 1 minus 2 and (2) contact 3 minus 4 (Table S3). By definition, contact 1 was most ventral and contact 4 was most dorsal.

One patient participant’s icEEG recordings were made with Natus NeuroWorks Quantum (8.5.1 Build 6067; Natus, Inc.) while the participant was being seen for in-house seizure monitoring by the Yale-New Haven Hospital Comprehensive Epilepsy Program. The patient was implanted with intracranial electrodes for seizure localization, including one unilaterally positioned 12-contact depth lead targeting the thalamus. Localization of these contacts (see the *Depth Contact and Channel Localization* section) revealed that the three most anterior contacts were embedded in the body of the thalamus, thus, included for analyses along with the icEEG recordings from the RNS System patient participants. These data were acquired at a sampling rate of 4096Hz and referenced to the first contact of a 1×4 frontal depth lead. Hardware low and high-pass filter frequency cutoffs were set to 0.01 and 1757Hz, respectively. To better correspond with the RNS System icEEG recordings, the three thalamic contacts were bipolar referenced to each other for analysis (contact 1 minus 2; contact 2 minus 3), resulting in two icEEG recording channels (Table S3).

#### Data Processing and Rejections

As detailed in the event synchronization section, the research tools laptop directed the RNS Tablet Programmer to interrupt the telemetry with the RNS Neurostimulator creating a reproducible artifact in the icEEG recordings that marked the onset of each task trial. These artifact-based event marks were synchronized to the trial onset. In post-processing, event marks were found automatically by an in-house outlier detection procedure programmed in MATLAB. First, one of the 4 icEEG channels was selected for identifying event time marks. While all channels were simultaneously marked, channels with less baseline signal fluctuations were selected that best emphasize the event markers. Next, the selected channel voltage was z-scored to the mean and standard deviation of all samples within each testing phase run of continuous icEEG recordings. All samples that exceed a threshold of −5µV z-scored voltage were tagged as outliers. Note that only negative z-scored voltage outliers were queried because the event markers created rapid apparent *decreases*. Subsequently, a custom outlier sample-sequence matching algorithm detected among the identified outlier samples the precise distribution and duration of outlier samples that corresponded to the reproducible profile of the induced event markers by telemetry interruption. All outlier samples that matched the known artifactual sequence were designated as event synchronization marks and were confirmed by review from a trained experimenter. After visual inspection, missing artifacts were restored or inaccurately identified event marks were excluded from consideration.

Stimulus presentation epochs were cropped for analysis using the behavioral task logged pre-stimulus intervals to calculate the duration between the icEEG trial-onset event mark and stimulus presentation. Stimulus-centered epochs were cropped to include 2 seconds pre and 2 seconds post-stimulus presentation (total epoch duration 4 seconds). All channels and epochs were visually inspected by a trained experimenter to remove from analyses any trial with samples that included artifact (e.g., dropped signal) or epileptiform activity. Only one participant had trials removed due to epileptiform activity (35.15% of perceived and 51.06% of not perceived channel-trial pairs). A total of 5.79% of perceived and 8.14% not perceived channel-trial pairs were rejected across all participants by data quality rejections by visual inspection.

The RNS icEEG recordings from one participant had a constant baseline voltage offset of 5µV in two icEEG channels and 5.5µV in the other two icEEG channels. These voltage offsets were eliminated by an additional preprocessing stage for this participant of subtracting the corresponding channel offsets from all icEEG data samples across recording sessions. This preprocessing step was not necessary for the other participant data because their recordings did not include a baseline voltage offset.

#### Channel Selection, Polarity Identification, and Visualization

The icEEG depth contact locations were heterogeneous (see the *Depth Contact and Channel Localization* section). Accordingly, some channels and their corresponding contacts may demonstrate potentials ranging from no response to robust signals for conscious visual perception based on anatomical/functional location. Therefore, channel selection was implemented to include only the icEEG channels that demonstrated responses linked to conscious visual perception for subsequent visualization and analyses. Channel selection also helped to localize the anatomy most responsive to conscious visual perception. Channels were selected based on peak voltage amplitude following perceived task stimuli, which was determined by a staged process. First, the average voltage of all perceived stimuli trials was computed for each icEEG channel across all participants. Not perceived stimuli trials were not considered in channel selection. Next, the absolute value of the mean channel voltage was computed and the maximum voltage value was found for each channel in the first 2000ms post-stimulus presentation. If the maximum voltage value was greater than 3µV the channel was labeled as “above threshold”, while channels with maximum voltage less than 3µV were labeled as “below threshold” (e.g., see Fig. 2D and fig. S12). The 3µV voltage threshold was selected based on visual inspection that revealed this voltage cut-off categorized channels with and without perception-link responses. Only above threshold channels were included in subsequent visualization and analysis.

The thalamic icEEG potentials for perceived stimuli was found to have a biphasic response with two polarity variants: (1) first peak positive and second peak negative, or (2) first peak negative and second peak positive (e.g., see fig. S14A channel 1 versus channel 2; Table S3, 1^st^ Peak Polarity). Interestingly, for all participants where both polarity types were seen (Table S3, participants 1, 3, 6, and 7) the electrode contacts with first peak positive were more dorsal than the contacts with the first peak negative, suggesting that a phase reversal occurred in the bipolar recordings along the electrode shaft within the thalamus. All above threshold channels were visually inspected to determine its polarity identity of either (1) or (2). Prior to computing the group mean across icEEG channels, for any channel with the polarity identity of (2) (i.e., a negative first peak), the voltage of all perceived and not perceived trials for that channel was flipped by multiplying the channel trial voltage by −1, converting the voltage polarity profile from (2) to (1). This sign transformation was necessary to prevent negation of signal when averaging across channels with opposite polarity.

Finally, for visualization of the group mean icEEG responses (Fig. 2E; fig. S14), prior to averaging all above threshold channel perceived and not perceived trials, bandpass filtering was applied with a custom finite impulse response filter (MATLAB *designfilt* and *filtfilt* functions) with high and low-pass bands of 0.5 and 14Hz, respectively. Bandpass filtering was implemented to eliminate low frequency drifts and high frequency noise.

#### Depth Contact and Channel Localization

The depth contact locations for the RNS System patient participants were found using Lead-DBS (https://www.lead-dbs.org) (*42*). In summary, the patient pre-operative T1-weighted anatomical MRI and post-operative CT images were coregistered and normalized to the MNI brain template space. Lead localization was guided by the Morel Atlas and using the PaCER, TRAC/CORE, or manual targeting methodologies in Lead-DBS (*43*). All localizations were visually inspected for accuracy. The depth contact locations for the single patient participant acquired with Natus NeuroWorks (Natus, Inc.) is detailed in (*9*). In summary, the pre and post-operative anatomical MRIs and post-operative CT images were coregistered and normalized to MNI space using BioImage Suite (https://bioimagesuiteweb.github.io/webapp). Anatomical localization for each contact is reported in Table S2.

Depth icEEG channel locations were determined by finding the central 3-dimensional coordinate between the contributing contacts of each bipolar referenced channel. This was achieved by computing the mean x, y, and z coordinates between the two contributing channel contacts and rounding the resulting coordinate values to the nearest integer (mm). The resulting channel locations were visualized on a high-resolution brain (0.3mm^3^ voxels) normalized to MNI space, overlaid with selected human thalamic sites voxelized from the Morel atlas (*43–45*) (Fig. 2D; fig. S12). The MNI coordinates for each channel is reported in Table S3.

### Eye Tracking and Pupillometry – EyeLink

#### Equipment and Software

Eye tracking and pupillometry data were collected with the EyeLink 1000 Plus System and software (version 5.09; SR Research, Inc.) running on a Dell PC desktop (Model D13M; Dell, Inc.). During hdEEG acquisition, head-fixed, binocular EyeLink recordings were acquired at 1000Hz with a 35mm camera lens and infrared illuminator mounted below the task LCD display. The participant head position was stabilized by a table mounted chin-rest that avoided contact with the face and forehead scalp electrodes. During fMRI acquisition, head-fixed (stabilized by the head-coil and padding), long range monocular (right eye) EyeLink recordings were acquired at 1000Hz with an MR-compatible camera and infrared illuminator mounted to a mounting bar and affixed to a stand placed inside the magnet bore behind the participant. The camera lens was oriented to view the participants’ right eye via the head-coil mounted mirror that the participants used to view the projector screen displaying the experimental task. To limit instances of the EyeLink software tracking spurious, non-eye sites (e.g., hdEEG scalp electrodes or the fMRI head-coil element), search limits were drawn on the EyeLink camera image to exclude regions beyond the area immediately surrounding the eyes.

Prior to each testing session, pupil and corneal reflection thresholds were set, used to search the camera image and identify the pupil and eye glint from the corneal reflection. If tracking was lost during the testing session, prior to continuing the next task block, the pupil and corneal reflection thresholds were updated to restore tracking. For most participants, the pupil was fitted using the EyeLink centroid procedure that implements a center of mass algorithm to estimate the pupil mass (SR Research EyeLink 1000 User Manual Section 3.6). However, for participants where the pupil was occluded (e.g., drooping eyelids and eyelashes), the ellipse procedure was selected that uses a ellipse-fitting algorithm designed to better handle partial pupil occlusions.

At the onset of each study session, the participant on-screen gaze position was estimated by a 9 or 5-point automated EyeLink gaze calibration procedure for the hdEEG and fMRI sessions, respectively. The gaze position was confirmed by EyeLink gaze position validation and the gaze calibration estimation was repeated if the validation step found poor coherence with the initial estimates for on-screen gaze position. The gaze position was monitored by the experimenter throughout the study session and if at any time the gaze position drifted from the known screen fixation point (often due to a drifting head position), a preprogrammed online gaze drift correction was applied to recenter the gaze position to the central fixation cross (SR Research EyeLink 1000 User Manual Section 3.11). If the gaze position was severely altered during the study session (e.g., by large head movements), prior to continuing the next task run, gaze calibration and validation were repeated to re-estimate the gaze position.

### Data Processing

Four data types were extracted from the EyeLink recordings: (1) artifact-interpolated pupil diameter, (2) artifact-interpolated gaze position, (3) blink occurrence, and (4) microsaccade occurrence. The extraction procedure for each of these data types first involved cropping out 12-second pupil diameter and gaze position (x and y-axes) epochs for analysis centered around stimulus presentation (i.e., 6 seconds pre and post-stimulus presentation). Next, artifacts were removed from the pupil and gaze timecourses on a trial-by-trial basis using the artifact identification procedure *Stublinks* (Siegle, 2003). In summary, Stublinks uses the pupil timecourse data to find artifactual time points, including blinks that are subsequently extracted from the pupil and gaze timecourses and interpolated to restore rejected samples. First, Stublinks down sampled the pupil timecourses data from 1000 to 60Hz. Next, artifactual time points were identified based on the following criteria: (1) pupil diameter changes of more than 0.5mm between consecutive samples, (2) pupil diameter below a threshold of 0.1mm or above 4mm from the median pupil diameter during each trial, (3) pupil diameter changes of 0.4mm or more across any four consecutive samples, (4) time points with more than 1mm difference for any sample between Stublinks smoothed (MATLAB *linspace* function) and unsmoothed pupil timecourses, and (5) time points that fell outside the Tukey’s test interquartile range (i.e., below or above the lower and upper hinges, respectively) for the smoothed pupil timecourses. All time points identified with irregularities from the above criteria that lasted between 100-400ms were labeled as blinks based on the known blink duration (*46*). Meanwhile, all other consecutive artifactual samples that persisted for less than 100ms or more than 400ms were marked as non-blink, miscellaneous artifacts. These miscellaneous artifacts were commonly prolonged eye closures or instances of eye tracking loss. Finally, all identified artifactual samples, both blink and miscellaneous events, were removed from the pupil and gaze timecourses and the rejected samples linearly interpolated (MATLAB *interpl* function) with temporally adjacent samples to restore the omitted time points.

The blink occurrence timecourses were constructed by converting the Stublinks-identified blink samples into a binary vector where for each 1000Hz sample in an epoch a value of 0 or 1 indicated a blink or the absence of a blink, respectively. Microsaccade occurrence timecourses were made by first finding instances of saccades in the x and y-axis gaze timecourses determined when the gaze position exceed a velocity threshold of 5 degrees/second of visual angle in 2-dimensional gaze-velocity space with a minimum duration of 5ms to protect against identifying high-frequency noise as saccades (*47*). From these saccades, microsaccades were isolated by extracting all saccades with movement of less than 1 degree of visual angle (*48*). Microsaccade occurrence timecourses were stored as a binary vector where each sample in an epoch was assigned either a 0 or 1 to designate a microsaccade or the absence of a microsaccade event, respectively.

When binocular EyeLink recordings were acquired (hdEEG data sets), EyeLink data processing was applied on the left and right eyes independently. Subsequent implementation and analyses of these EyeLink data only considered the right eye for consistency with the right eye, monocular EyeLink recordings acquired in the Report + No-Report Paradigm fMRI data set.

### EyeLink-Based Rejections

EyeLink blink occurrences during stimulus presentation were used as a trial-based rejection criterion, as described in the *Functional Magnetic Resonance Imaging (fMRI)* and *Low and High-Density Scalp EEG (ldEEG and hdEEG)* sections. In summary, for the Report + No-Report Paradigm fMRI and EyeLink data sets, trials were rejected when a blink occurred during the 50ms stimulus presentation period, defined as when there was at least 1 sample of blink occurrence during the stimulus presentation. For the Report and Report + No-Report Paradigms hdEEG data sets, trials were rejected when a blink occurred between 200ms pre-stimulus and 500ms post-stimulus to avoid the electrophysiological artifact blinks can induce in scalp sensors. Finally, an entire run for one participant completing the Report + No-Report Paradigm with the hdEEG procedure was excluded from analyses due to lost EyeLink data acquisition.

### Covert Prediction of Conscious Perception – Machine Learning Pipeline

A machine learning pipeline was implemented with the goal of achieving accurate trail-by-trial predictions of perceived and not perceived no-report (task-irrelevant) stimuli from the Report + No-Report Paradigm in lieu of overt report. The pipeline includes trial segmentation, feature extraction, feature normalization, feature selection, information fusion, and classification stages. As detailed below, the perception class (perceived or not perceived) of task stimuli was predicted using classification models trained on the EyeLink data (see the *Eye Tracking and Pupillometry – EyeLink* section).

### Software

The machine learning pipeline was implemented in MATLAB using custom functions and those available in the MATLAB Statistics and Machine Learning Toolbox. The python package tsfresh (Version 0.18.0) was used to extract time series features (https://tsfresh.readthedocs.io/en/latest/).

### Trial Segmentation and Feature Extraction

Eye dynamics have been shown to be a covert measure of perceptual and attentive states, including the optokinetic nystagmus (*10, 49*), pupil diameter (*10, 49–51*), fixation position (*11*), saccade latency (*52*), microsaccade frequency, (*53, 54*) and blink frequency (*54, 55*). The current investigation builds on these findings to develop a covert marker of perception by implementing machine learning and multiple eye tracking and pupillometry-based features to inform trial-based predictions of the perception of visual stimuli. All data used in machine learning (i.e., the training and testing sets) were acquired from the Report + No-Report Paradigm, where the report (task-relevant) trials formed the training set and the no-report (task-irrelevant) trials formed the testing set (fig. S3A). Only trials with perceptual threshold opacity task-relevant or irrelevant stimuli were considered in the training and testing data sets, respectively (i.e., opaque and blank stimuli trials were excluded). To maximize the size of the training and testing data sets, all trials with available processed EyeLink data (see the *Eye Tracking and Pupillometry – EyeLink* section) were considered for machine learning, including those trials that would be subsequently rejected by the trial and testing block rejections described earlier, for example, rejecting testing phase runs with excessive movement during fMRI acquisition and rejecting trials with blinks at stimulus presentation (fMRI) or during a critical window (hdEEG). However, as detailed in their respective method sections, all subsequent fMRI and hdEEG analyses applied block and trial-based rejection criteria for both the overt report and classified or predicted, no-report perceived and not perceived trials.

The extracted feature categories included: (1) pupil diameter timecourse, (2) tsfresh features from pupil diameter timecourse, (3) blink occurrence binary timecourse, (4) microsaccade occurrence binary timecourse, and (5) x and y-axis gaze position timecourses (see the *Eye Tracking and Pupillometry – EyeLink* section). The 12-second pupil, blink, microsaccade, and gaze timecourse epochs from the right eye EyeLink recordings corresponding to each trial were cropped to 3 seconds post-stimulus epochs and then resampled from 1000 to 50Hz (i.e., 3000 to 150 samples) in order to improve computational efficiency and apply feature extraction on the most salient post-stimulus temporal window. Various post-stimulus periods (1 to 6 seconds post-stimulus) and resampling values (1000 or no resampling to 10Hz) were tested and the values of 3 seconds post-stimulus and resampling to 50Hz were selected because they maximized classification performance. Each of the 150 samples representing the first 3 seconds post-stimulus were considered as features for classification yielding a total of 750 features (5 x 150) representing the timecourses of pupil diameter, blink occurrence, microsaccade occurrence, and x and y-axis gaze position. Meanwhile, the entire 12-second pupil timecourse epoch at 1000Hz was considered for tsfresh feature extraction yielding 784 tsfresh features per trial, however, only a subset of these features were used after feature selection (see the *Pupil Diameter Tsfresh Feature Selection* section; Table S4). Therefore, each trial was represented by 1534 features in total.

### Pupil Diameter Tsfresh Feature Selection

Feature selection was employed on the pupil diameter tsfresh features to select the tsfresh features that are relevant to the classification problem of interest. To select the most predictive pupil diameter tsfresh features, a two-sample *t*-test was used to find which features were statistically significant (*p* < 0.05) between report perceived and not perceived trials (i.e., task-37 relevant stimuli). A total of approximately 300 of the 784 tsfresh features were found statistically significant between perceived and not perceived task-relevant report trials and only these tsfresh features were considered for training and testing the classification models for predicting the perception class of task-irrelevant stimuli. Table S4 details the number of features used in each data set and the tsfresh feature categories (see tsfresh documentation for full details on each of the listed tsfresh features: https://tsfresh.readthedocs.io/en/latest/text/list_of_features.html).

### Information Fusion and Classification

Predictions of the perception class of no-report stimuli were made using a two-staged, stacked ensemble classifier with an input combining information from different eye tracking and pupillometry measurements. In the first stage or base level, each of the feature sets were individually fed into three base models: (1) an ensemble, boosted decision tree, (2) a linear kernel support vector machine (SVM), and (3) a gaussian kernel SVM (fig. S3B). The base models were selected after extensive testing of numerous alternative machine learning and deep learning models, which had worse performance than the three selected models. The boosted decision tree was implemented with the MATLAB *fitcensemble* function with default parameters. The SVM base models were implemented with the MATLAB *fitcsvm* function, specifying the cross-trial standardization and gaussian or linear kernel parameters. The report (task-relevant) data were used to optimize the performance of the base models in a 10-fold cross-validation scheme (see the *Model Training* section). In particular, the hyperparameters of the base models were selected such that the prediction accuracy and retention (both defined below, see *Performance Evaluation* section) of the report perceived and not perceived trials were maximized. Instead of considering the predicted labels obtained from the base models for deciding on the perception class, we used the scores corresponding to the predicted labels as input into an ensemble prediction layer. The total output from the base level was 18 unique scores per trial (note that x and y components of gaze position were applied as independent features for a total of six feature sets that were separately applied among the three base models).

In the second stage or stacked ensemble level, the base model scores per trial were considered as input features for a linear kernel SVM (fig. S3B). The trial perception class predictions were made based on the scores of the ensemble level linear SVM. The predicted trial perception class label was determined by applying a positive and negative class (i.e., perceived and not perceived) score thresholds, such that any trial with a score above or below those thresholds would be labeled in their respective class (fig. S3B). The purpose of applying score thresholds was to allow the experimenters to balance classification accuracy and the retention of trials (sample size) necessary to resolve a response in the physiological recordings. After thorough testing of various score thresholds, the thresholds of 0.75 for perceived and −0.75 for not perceived classes was selected to determine the predicted trial class label. See the *Performance Evaluation* section for details on selecting the score threshold.

### Model Training

The base and ensemble level models were trained on extracted features obtained from the report (task-relevant) trials from the Report + No-Report Paradigm. For these trials the true perception class labels were known by overt report. For hyperparameter optimization (see the *Information Fusion and Classification* section) and performance evaluation on the report trials, training data were partitioned into training and testing sets using a 10-fold cross-validation scheme (MATLAB *cvpartition* function). The same subsample folds were used for all models to prevent cross model training and testing set contamination. Each behavioral task-relevant condition by location set (center and quadrant location sets) and neuroimaging modality (fMRI and hdEEG) study session combinations were considered as independent data sets each consisting of over 4000 perceived plus not perceived trials (fig. S4; Table S5), and each were used for training their own set of base and ensemble models. Therefore, a total of 4 base and ensemble model sets were trained for each study session behavioral and modality condition combination: (1) center task-relevant fMRI (2) quadrant task-relevant fMRI, (3) center task-relevant hdEEG, and (4) quadrant task-relevant hdEEG. Subsequently, the perceptual class predictions of no-report trials were made from the trained models of its corresponding study session. Training and testing within study sessions were implemented instead of combining trials across study sessions because performance evaluation (see the *Performance Evaluation* section) determined within study session predictions improved classification performance that may be because of idiosyncrasies of the feature data among the study sessions. The total number of trials in the training and testing sets for each study session are listed in Tables S5 and 6, respectively.

### Model Testing

The no-report (task-irrelevant) trials from the Report + No-Report Paradigm were tested using the trained base and ensemble models that had been trained with the report (task-relevant) trials from the Report + No-Report Paradigm (fig. S3A). Specifically, the test data were passed through the same classification architecture, beginning with the features from the no-report trials as inputs to the base models: ensemble, boosted decision tree, linear kernel SVM, and gaussian kernel SVM. The scores from the base models for each trial formed the features for the stacked ensemble linear kernel SVM model. The pre-selected score thresholds of 0.75 and −0.75 for the perceived and not perceived classes, respectively, were applied on the scores from the ensemble model to supply the no-report trial perceptual class predictions: perceived or not perceived.

Trials that did not exceed the score thresholds remained unlabeled (fig. S5). The final predictions resulted in a total of 4019 predicted perceived and 5417 predicted not perceived no-report trials across all study session conditions (see Table S6 for a breakdown of each study session). Subsequent analyses of EyeLink, fMRI, and hdEEG data used these predicted labels of perception class for the no-report trials and are referred to as predicted perceived and predicted not perceived to distinguish them from the labeled, overt report perceived and not perceived trials.

### Performance Evaluation

The implementation of pupillometry and eye tracking as a method for predicting the trial perception class was premised on report perceived and not perceived data showing unique, trial-level post-stimulus responses in pupil, blink, and microsaccade dynamics. Training and testing of the classification models on the report data supports this assumption.

While it is common to measure the performance of a classifier by its sensitivity and specificity behavior (i.e., a receiver operating characteristics curve), the unique challenge of the current classification problem was to balance prediction accuracy and trial retention sufficient for subsequent analysis of physiology. Therefore, the classifier quality was measured by positive (PPV) and negative predictive values (NPV) for the perceived and not perceived classes, respectively, against the trial retention. PPV and NPV were used as measures of prediction accuracy, and respectively, represent the proportion of trials classified correctly among those classified as perceived or not perceived. These proportions are important because they determine how much “contamination” from incorrectly classified trials exists in the classified perceived or not perceived data used ultimately for fMRI or hdEEG analyses. Retention is important because it determines sufficient sample sizes for the planned fMRI and hdEEG analyses. PPV, NPV, and retention were calculated as:

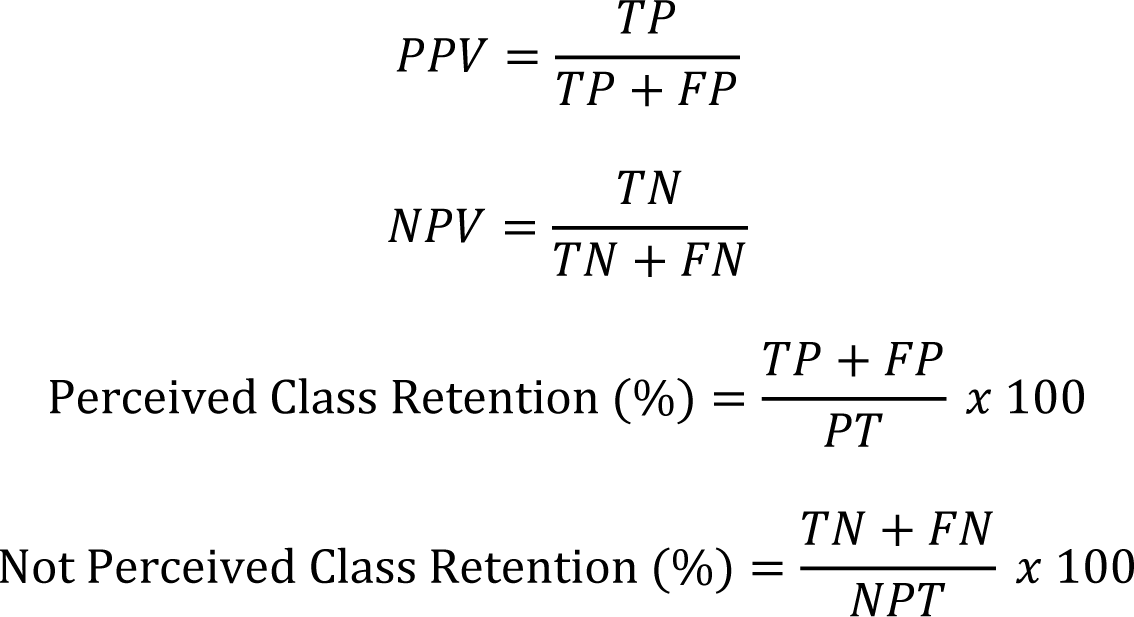

where TP are the true positive classifications (perceived trials classified as perceived), FN are the false negative classifications (perceived trials classified as not perceived), TN are the true negative classifications (not perceived trials classified as not perceived), FP are the false positive classifications (not perceived trials classified as perceived), PT is the total number of perceived trials (equal to TP + FN), and NPT is the total number of not perceived trials (equal to TN + FP). PT and NPT are known in the report condition by overt report.

The aggregated performance of models across each of the four study session task and neuroimaging condition combinations using a score threshold of 0 on the stacked ensemble SVM trial scores was 0.74 and 0.70 for PPV and NPV, with a retention of 107% and 91%, respectively (see points labeled 1 in fig. S4 and 0 score threshold in Table S5; retention can exceed 100% when the total number of predicted class trials exceed the number of true class trials). As the score thresholds become more stringent, the PPV and NPV values increases while retention decreases (fig. S4; Table S5). Retention decreases because trials with score values between the score thresholds remain unlabeled. Therefore, the numerator for calculating retention (the predicted number of perceived and not perceived trials) decreases with more stringent score thresholds, while the denominator (the true number of perceived and not perceived trials) remains constant. For example, PPV and NPV performance of >0.90 can be achieved but at the cost of a retention <30% (see points labeled 7, 8, and 9 in fig. S4 and −1.5/1.5, −1.75/1.75, and −2.0/2.0 score thresholds in Table S5). The optimal score thresholds were determined by finding the scores that balanced PPV and NPV, and retention. Therefore, the score thresholds of 0.75 and −0.75 for the perceived and not perceived classes, respectively, were selected because they offered >0.80 for both PPV and NPV with retention of approximately 60% and 50% for the perceived and not perceived class, respectively (see bolded points labeled 4 in fig. S4 and −0.75/0.75 score thresholds in Table S5). The performance from alternative score thresholds ranging between 0.25 to 2 for predicted perceived and −0.25 to −2 for predicted not perceived in increments of 0.25 are reported in fig. S4 and Table S5. In summary, PPV and NPV versus retention curves shows that the classifier performed slightly better on the perceived class (PPV curves) relative to the not perceived class (NPV curves), and there was minor variability in the classification performance among each of the four study session types.

As an additional check on the classification accuracy and performance, the fMRI and hdEEG physiology of the predicted perceived and not perceived trials on the report data (from 10-fold cross-validation) were compared to the physiology using the true perception class trial labels from overt report. The predicted and true class fMRI and hdEEG responses from a score threshold of 0.75 and −0.75 reveal similar physiological results (data not shown), suggesting this balance of PPV/NPV (>0.80) and trial retention (>50%) is sufficient to achieve the original results with the labeled report-based data.

While classification performance on the unlabeled, no-report data is challenging to assess without overt report, the classification profile suggests a similar performance to that found in the labeled, report data sets. First, the distribution of scores from the stacked ensemble SVM are similar for report and no-report data (histograms in fig. S4 versus fig. S5). Moreover, there is minimal class bias, apparent when adjusting the score threshold by equal measure above and below zero resulting in approximately similar numbers of predicted perceived and not perceived trials, although a not perceived class bias is apparent for the center location set task-irrelevant hdEEG condition (fig. S5C). In addition, the yield of predicted trials from no-report classification matches across score thresholds (yellow and green curves plotted in fig. S5). The approximate even number of trials predicted as perceived and not perceived is predicted by the performance achieved in the classified report data set for perceptual threshold opacity stimuli.

Finally, application of the classifier to no-report data yielded pupil, blink, and microsaccade mean timecourses for classified perceived and classified not perceive trials that resembled the mean timecourses for perceived and not perceived trials in the report data (Fig. 1C versus D; fig. S8A versus B).

Together, these results support that pupil, gaze, blink, and microsaccade dynamics can be used for trial-level predictions of perception, and that classification performance is sufficient with a large data set to achieve high classification accuracy while maintaining a sufficient data set size for subsequently analyses of physiology.

### Statistical Analyses

Behavioral, EyeLink, hdEEG, ldEEG, icEEG, and fMRI statistical analyses and data visualizations were executed with GraphPad Prism (version 9.1.2, GraphPad, Inc.) or custom and available MATLAB functions and toolboxes, including SPM and EEGLAB.

#### Behavioral Analysis

Behavioral performance was assessed to answer two primary questions on the report, task-relevant stimuli: (1) What proportion of stimuli are seen, when stimulus opacity is at perceptual threshold, at 0% (blanks), or at 100% (fully opaque)? And, (2) what is the location accuracy for seen stimuli and for not seen stimuli? The main analyses to resolve both questions combined the performance across trials, participants, and task conditions using report (task-relevant) data from the Report and Report + No-Report Paradigms (although only the Report + No-Report Paradigm include fully opaque stimuli). Additional behavioral performance analyses observed if stimulus perception rate was modulated by (1) stimulus location on the screen (i.e., visual fields in the Report Paradigm and center versus quadrant location sets in the Report + No-Report Paradigm), (2) the simultaneously recorded modality (i.e., hdEEG, ldEEG, icEEG, and fMRI), (3) the post-stimulus delay interval (i.e., 1 or 15 seconds and 3, 4, or 5 seconds for the Report Paradigm variants, and 6-10 seconds for the Report + No-Report Paradigm), (4) the screen background type (i.e., documentary versus static noise) in the Report Paradigm, and (5) whether the task-relevant stimulus appeared first or second in each trial for the Report + No-Report Paradigm. See fig. S6 and 7 for behavioral performance results from the Report and Report + No-Report Paradigms, respectively.

#### fMRI, hdEEG, and ldEEG Spatiotemporal Analyses

The multiple comparisons problem in statistical analyses was a concern in the current investigation because of the acquisition of high-dimensional data sets, including fMRI that is susceptible to false discoveries (*56*). In addition, analyses requiring a hemodynamic response model can miss important spatiotemporal signals that do not fit the model in some brain regions (*37, 57, 58*). Therefore, we implemented model-free cluster-based permutation tests to identify statistically significant spatiotemporal clusters from whole brain fMRI and scalp hdEEG and ldEEG. Cluster-based permutation tests have shown to be a powerful tool for limiting false positive (Type I) error rates in high-dimensional data, including EEG and fMRI.

The core principle of the cluster-based permutation analysis is to compare the experimental data to a null distribution built from a cluster-forming statistic generated after iterations of randomly permuting the experimental data (*59*). A modified version of the MATLAB Mass Univariate ERP Toolbox *clust_perm1* function was implemented to generate the null distribution (*60*). A previous fMRI study shows stability in statistically significant clusters from cluster-based permutation tests above 5000 permutation iterations, and particularly above 10000 iterations (*61*). Cluster-based permutation analysis of the BOLD dynamics were tested with both 5000 and 10000 iterations and both showed stability of the statistically significant spatiotemporal clusters (results not shown). Therefore, 5000 iterations was selected for both fMRI and EEG cluster-based permutation analyses to balance the competing demands of computational efficiency and robust statistical results.

Prior to implementing cluster-based permutation analysis, the fMRI and EEG data were preprocessed specifically for this statistical analysis (see also *Data Preprocessing and Artifact Rejection* described above). First, for whole brain fMRI cluster-based permutation analysis, the spatial dimension of these data were reduced to improve computational efficiency. This was achieved by excluding from analyses all non-grey matter voxels, implemented by applying a whole brain grey matter mask created in the SPM toolbox MarsBaR, as described previously (*37*). Next, the whole brain volume spatial resolution was reduced from 2 to 6mm^3^ voxels. This was achieved by combining spatially adjacent 2mm^3^ grey matter voxels into combined-voxels of 6mm^3^. Specifically, the central voxel for each of the combined 6mm^3^ voxels was found by selecting all the original 2mm^3^ voxels that were regularly positioned with exactly 2 intervening voxels until the next central voxel in the x, y and z directions, applied across the entire whole brain, grey matter-masked volume. For each central voxel, all adjacent voxels were found (i.e., any voxel that shared a face, edge, or vertex with the central voxel). These adjacent voxels along with the central voxel would form the combined 6mm^3^ voxels. In total, the 6mm^3^ voxels would comprise a maximum of 27 (^33^) of the original 2mm^3^ voxels. In some instances a complete 6mm^3^ voxel could not be built if the adjacent voxels were excluded because they occupied non-grey matter territories (i.e., at the edges of the brain grey matter). In these cases, the newly constructed voxel would take all available adjacent voxels and, consequently, its shape would be non-cuboidal. This spatial down sampling procedure reduced the number of voxels considered in whole brain cluster-based permutation analysis from 902,629 to 6934 voxels. Finally, the BOLD signals for the new, lower spatial resolution voxels were determined by computing the mean BOLD signal across all 2mm^3^ voxels within each of the constructed 6mm^3^ voxels at each sample/volume.

Likewise, the high temporal resolution of the EEG signals (hdEEG: 1000Hz; ldEEG: 256Hz), introduced a computational challenge for the spatiotemporal cluster-based permutation analysis. Instead of a temporal down sampling procedure, as applied to the spatial dimension of the fMRI data, statistical analyses were focused on only the samples representing 500ms pre-stimulus to 1500ms post-stimulus, a total of 2000 and 512 samples for the hdEEG and ldEEG data trials, respectively. By analyzing only the critical temporal window of these EEG data, the processing speed was more than doubled compared to using the entire original 4-second epochs of 2000ms pre and post-stimulus.

The following conditions were tested with spatiotemporal cluster-based permutation analysis within the report (from the Report Paradigm trials and the task-relevant trials of the Report + No-Report Paradigm) and no-report (from the task-irrelevant trials of the Report + No-Report Paradigm) conditions: (1) perceived versus baseline, (2) not perceived versus baseline, and (3) perceived versus not perceived. The comparative baseline values were taken from the mean percent change BOLD or voltage of the first 20 seconds and 1000ms pre-stimulus presentation for the fMRI and EEG data sets, respectively. Comparative tests between the report and no-report conditions were also tested: (1) report perceived versus no-report perceived, (2) report not perceived versus no-report not perceived, and (3) report perceived minus not perceived versus no-report perceived minus not perceived. One participant was excluded from the cluster-based permutation analyses that tested on the fMRI no-report perceived stimuli condition and two participants were excluded from cluster-based permutation analyses that tested on the hdEEG no-report not perceived stimuli condition because these participants did not have no-report perceived (fMRI subject) or not perceived trials (hdEEG subjects) at the 0.75 and −0.75 classification score thresholds, respectively (see the *Covert Prediction of Conscious Perception – Machine Learning Pipeline* section for details on classification scores for no-report trial predictions).

Cluster-based permutation analysis was processed by first generating the spatiotemporal cluster null distribution over 5000 permutation iterations. For each permutation iteration, participant pre-stimulus baseline or comparative group BOLD or voltage values for fMRI and EEG, respectively, were randomly shuffled with the participant-paired test value (i.e., the current sample being tested). Next, a paired, two-tailed *t*-test compared the permuted distributions across participants and found statistically significant voxels or electrodes for fMRI and EEG, respectively, at the tested time point (*p* < 0.05). Each epoch time point was tested independently within each permutation iteration (fMRI: 20 seconds pre and post-stimulus; EEG: 500ms pre-stimulus and 1500ms post-stimulus). After all epoch time points were tested, statistically significant spatiotemporal clusters were formed by spatial and temporal adjacencies. Negative and positive clusters were treated independently. For fMRI, spatial adjacency was defined as when statistically significant voxels (of the same sign) shared a face, edge, or vertex, while in EEG spatial adjacency was determined by statistically significant channels neighboring each other on the scalp surface. Meanwhile, temporal adjacency was determined if any voxel or electrode was found statistically significant (in the same direction) between two or more sequential time points. For each identified spatiotemporal cluster, the summed absolute value of *t*-values were computed across all cluster contributing voxels or electrodes and for all time points contributing to that cluster. The largest negative and positive cluster determined separately by summed absolute value of *t*-values was selected from each permutation iteration and added to the respective negative and positive spatiotemporal cluster null distribution. This procedure was repeated for 5000 iterations.

After constructing the spatiotemporal cluster null distribution, a final iteration of the spatiotemporal cluster forming analysis detailed above was completed, but, critically, without permutation. Thus, negative and positive spatiotemporal clusters were found on the original data set, their size measured by the metric of summed absolute value of *t*-values, and any cluster that exceeded the outer top 5% of the two-tailed null distribution was considered to be statistically significant.

To implement the above approach while shortening computation time, we constructed the permutation distribution in only one direction, assuming a symmetrical distribution with random shuffling and sufficient number of permutation iterations, and applied the single resulting threshold in both directions.

Visualization of the fMRI results from spatiotemporal cluster-based permutation analysis were displayed on the standard MNI brain template with either the individual voxel *t*-values (Fig. 3A, B, C, and D) or the perceived minus not perceived percent change BOLD signal value (fig. S15) shown for all voxels and time points that belonged to a statistically significant spatiotemporal cluster. For the display of the EEG results, the channel voltage timecourses were updated for all electrodes so that all non-statistically significant samples were replaced with a voltage of 0.

Next, the mean voltage for each channel was computed within the queried event-related potential (ERP) time windows (N100: 75-125ms; VAN: 175-225ms; P2/N2: 275-325ms; P3: 350-650ms).

These ERP time windows were selected to match what has been reported in the literature (*16*). The resulting values were plotted for all channels on the scalp surface (MATLAB EEGLAB *pop_topoplot* function; Fig. 2A, B, and C; fig. S10).

#### fMRI, hdEEG, ldEEG, icEEG, and EyeLink Temporal Analyses

Statistical analysis of all timecourse data (i.e., 2-dimensional data of the structure signal by time), including fMRI k-means cluster (see the *fMRI Anatomical Clustering* section) and region of interest (ROI) timecourses, hdEEG, ldEEG, and icEEG channel timecourses, and EyeLink pupil diameter, blink rate, and microsaccade rate timecourses were implemented with an adapted version of the spatiotemporal cluster-based permutation test (see the *fMRI, hdEEG, and ldEEG Spatiotemporal Analyses* section). The critical difference for the cluster-based permutation analysis of the timecourse data was that the cluster-forming procedure considered only temporal adjacency, whereas in the previous implementation both spatial and temporal adjacencies were used to form spatiotemporal clusters. Also, due to the low participant sample size in the icEEG and ldEEG datasets, cluster-based permutation tests were performed on the trial-level for these data, as compared to the participant-level analysis implemented for the fMRI and hdEEG data sets. Otherwise, the cluster-based permutation analyses were implemented as reported for the spatiotemporal analysis.

In summary, a temporal cluster null distribution was formed over 5000 iterations of permuting between participant baseline (the mean of all samples within a pre-selected baseline period; fMRI baseline: 20 seconds pre-stimulus; EEG and EyeLink baseline: 1000ms pre-stimulus) or comparative condition values and test values (i.e., each pre and post-stimulus epoch sample of the test condition). Next, the permuted distribution was tested with a paired one-sample *t*-test and positive and negative clusters were found independently by statistically significant (*p* < 0.05) temporal adjacency. The size of each negative and positive temporal cluster was determined by summed absolute value of epoch sample *t*-values contributing to each cluster and the largest negative and positive cluster values were added to their respective null distributions. Finally, temporal clusters were found on the non-permuted data using the same procedure as for the permuted data and tested for statistical significance by comparison to the null distribution.

As in the spatiotemporal analyses, to implement the above temporal cluster-based permutation analysis approach while shortening computation time, we constructed the permutation distribution in only one direction, assuming a symmetrical distribution with random shuffling and sufficient number of permutation iterations, and applied the single resulting threshold in both directions.

For the EyeLink timecourses, cluster-based permutation tests were implemented comparing report (task-relevant from the Report + No-Report Paradigm) perceived versus not perceived (Fig. 1C; fig. S8A) and no-report (task-irrelevant from the Report + No-Report Paradigm) perceived versus not perceived (Fig. 1D; fig. S8B). For the fMRI k-means (see the *fMRI Anatomical Clustering* section; Fig. 4D, E, and F) and ROI timecourses (Fig. 4G-J; fig. S18-20), and EEG channel timecourses (fig. S11), the following conditions were tested within the report and no-report conditions: (1) perceived versus baseline, (2) not perceived versus baseline, and (3) perceived versus not-perceived. Comparative tests between the report and no-report conditions were also tested: (1) report perceived versus no-report perceived, (2) report not perceived versus no-report not perceived, and (3) report perceived minus not perceived versus no-report perceived minus not perceived. For the icEEG and ldEEG timecourses with only the Report Paradigm (i.e., patient participant data set), perceived versus not perceived statistical comparisons were made within each recording stream (Fig. 2E).

Just as in the spatiotemporal clustering procedure, one participant was excluded from the cluster-based permutation analyses that tested on the fMRI no-report perceived stimuli condition and two participants were excluded from cluster-based permutation analyses that tested on the hdEEG no-report not perceived stimuli condition because these participants did not have no-report perceived (fMRI subject) or not perceived trials (hdEEG subjects) at the 0.75 and −0.75 classification score thresholds, respectively (see the *Covert Prediction of Conscious Perception – Machine Learning Pipeline* section).

The fMRI ROI timecourses considered in cluster-based permutation analysis were taken from anatomical regions in the SPM toolbox MarsBaR (*62*) and custom designed in MarsBaR based on anatomical coordinates (midbrain and nucleus accumbens), and the Harvard-Oxford cortical and subcortical atlases (https://identifiers.org/neurovault.collection:262). Functional ROIs were selected from a 90-ROI, 14-network functional atlas developed at Stanford University (*63*).

When necessary, the ROIs were converted to MNI brain template space using SPM and MarsBaR. The voxels comprising the selected ROIs were grouped according to spatial overlap with the k-means clusters (k = 3) formed from the report perceived minus not perceived percent change BOLD dynamics (see the *fMRI Anatomical Clustering* section; Fig. 4A, B, and C), and the ROI was designated to a k-means cluster if a cluster owned more than 60% of voxels in the ROI relative to all other k-means cluster voxels overlapping with the current ROI (Fig. 4G-J; fig. S18-20). When plotting the ROI timecourses and running cluster-based permutation analyses, only those voxels that were shared between the ROI and the k-means cluster voxels from the report and no-report condition were considered.

#### fMRI Spatiotemporal Conjunction and Exclusive Disjunction Analyses

The goal of voxel-level conjunction and exclusive disjunction analyses was to emphasize the report-independent and report-dependent networks by comparing shared and unshared BOLD signals. The conjunction and exclusive disjunction voxels were queried among all whole brain grey matter voxels and over all epoch samples (20 seconds pre and post-stimulus presentation). A voxel at a particular time point could be statistically significant or not among the following cluster-based permutation tests (see the *fMRI, hdEEG, and ldEEG Spatiotemporal Analyses* section for full analyses details):

1. Report perceived minus not perceived
2. No-report perceived minus not perceived
3. (1) versus (2) (i.e., report perceived minus not perceived versus no-report perceived minus not perceived).

A voxel at a particular time was included in the conjunction data set if (1) and (2) were both statistically significant in the same direction (i.e., both positive or negative), and (3) was not statistically significant. Voxels with shared positive or negative changes (increases or decreases) were plotted separately.

For exclusive disjunction analysis, two variants were considered. The first variant showed greater changes-for-report and the second greater changes-for-no-report trials. In the greater changes-for-report exclusive disjunction analysis, a voxel at a particular time was included if the following held:

Greater increases for report:

1. showed a statistically significant increase,
2. did not show a statistically significant increase, and
3. showed that (1) was significantly greater than (2).

OR

Greater decreases for report:

1. showed a statistically significant decrease,
2. did not show a statistically significant decrease, and
3. showed that (1) was significantly less than (2).

In the greater changes-for-no-report exclusive disjunction analysis, a voxel at a particular time was included if the following held:

Greater increases for no-report: (2) showed a statistically significant increase,

(1) did not show a statistically significant increase, and

(3) showed that (2) was significantly greater than (1).

OR

Greater decreases for no-report:

(2) showed a statistically significant decrease

(1) did not show a statistically significant decrease, and

(3) showed that (2) was significantly less than (1).

#### fMRI Anatomical Clustering

Data-driven, voxel-level clustering of whole brain grey matter voxels was implemented with the clustering algorithm k-means (MATLAB *kmeans* function) on the percent change BOLD responses for report perceived minus not perceived stimuli, combined from the Report and Report + No-Report Paradigms. Voxels for anatomical clustering were selected by finding all voxels that were statistically significant for at least one sample during the first 10-second post-stimulus period, as determined from cluster-based permutation analysis of the report perceived minus not perceived stimuli percent change BOLD signal (Fig. 3A, B; Slide S1). For the included voxels, k-means clustering was applied to the perceived minus not perceived percent change BOLD signal in the first 10 seconds post-stimulus averaged across participants. The k-means clustering was implemented with a *Pearson* correlation distance metric between voxel time courses. Multiple k or cluster number values were tested (2-10 clusters). For each k-value k-means clustering iteration, the voxel-based silhouette values (a metric of cluster coherence) was estimated (MATLAB *silhouette* function) from the first 10 seconds post-stimulus, the mean timecourse of the percent change BOLD signal for perceived minus not perceived stimuli of all voxels within each cluster was computed and plotted for the entire 41-second BOLD epoch (20 seconds pre and post-stimulus presentation), and a 3-dimensional volume in MNI brain template space of all contributing cluster voxels was made in SPM with the MarsBaR toolbox to offer spatial visualization of the clusters. Together, the silhouette values, cluster mean percent change BOLD timecourses, and the cluster brain volumes were used to assess the parsimony and anatomical and functional identity of each k-means cluster. Together these metrics supported the voxel-wise clustering into 3 clusters (k = 3) was most coherent of all the clustering values tested in the report condition (Fig. 4A-C; fig. S17).

The k-means clusters found in this way for the report data were next applied to the no-report (task-irrelevant) perceived and not perceived stimuli percent change BOLD signals. To accomplish this for the no-report data, voxels were only included if found statistically significant for at least one sample in the 10-second post-stimulus period based on permutation analysis of the no-report perceived minus not perceived BOLD signals. Thus, anatomical clusters from the report data (Fig. 4A-C) were used to generate BOLD fMRI timecourses for the significant voxels in both the report and no-report data sets. Mean values within each anatomical region at each time point were used for the timecourses. This was done to visualize differences in the timecourses of both report-dependent and report-independent signals in different regions of the brain (Fig. 4G-J; fig. S18-20). Although k-means clustering applied directly on the no-report percent change BOLD signals was also implemented (results not shown), the report k-means clusters were selected because they included report-dependent clusters that were absent in the no-report condition.

To enhance the spatial visualization of the k-means clusters, cluster voxels were shown as belonging to either (1) a subcortical or cerebellar site (abbreviated here together as subcortical) or (2) the cortex (Fig. 4A-C). Subcortical and cortical visualization was achieved by determining the inclusion of k-means cluster voxels among selected anatomical ROIs. A voxel was designated as subcortical if it was located within selected subcortical MarsBaR (Amygdala, Caudate, all Cerebellum ROIs, Pallidum, Putamen, Thalamus, and all Vermis ROIs) or custom designed in MarsBaR (midbrain and nucleus accumbens), and Harvard-Oxford cortical and subcortical structural atlas ROIs (brainstem ROI, including the midbrain and pons taken from HarvardOxford-sub-maxprob-thr50-2mm.nii; https://identifiers.org/neurovault.collection:262). A voxel was found to belong to a cortical region if it was among selected cortical MarsBaR ROIs (Angular, Calcarine, Cingulum_Ant, Cingulum_Mid, Cingulum_Post, Cuneus, Frontal_Inf_Oper, Frontal_Inf_Orb, Frontal_Inf_Tri, Frontal_Med_Orb, Frontal_Mid, Frontal_Mid_Orb, Frontal_Sup, Frontal_Sup_Medial, Frontal_Sup_Orb, Fusiform, Heschl, Hippocampus, Insula, Lingual, Occipital_Inf, Occipital_Mid, Occipital_Sup, Olfactory, Paracentral_Lobule, ParaHippocampal, Parietal_Inf, Parietal_Sup, Postcentral, Precentral, Precuneus, Rectus, Rolandic_Oper, Supp_Motor_Area, SupraMarginal, Temporal_Inf, Temporal_Mid, Temporal_Pole_Mid, Temporal_Pole_Sup, and Temporal_Sup).

#### icEEG and ldEEG Latency Analysis

The thalamic icEEG and scalp ldEEG ERP peak times were tested for statistically significant latencies using a peak detection procedure. Only perceived stimuli trials and above threshold channels (see the *Intracranial EEG (icEEG)* section) were considered in the latency analyses. One participant with thalamic depth recordings did not have simultaneous ldEEG recordings (patient participant 1 in Table S2), therefore, this participant did not contribute to the ldEEG peak latency data set. First, as described earlier (see the *Intracranial EEG (icEEG)* section), the trials for channels with a negative first peak polarity were flipped (multiplied by −1) and bandpass filtered (high-pass: 0.5; low-pass: 14Hz). Next, the absolute value of the mean of all trials within each channel was computed. Third, the absolute value voltage local maximums within the first 2000ms post-stimulus were found with the MATLAB *findpeaks* function.

Knowing that the above threshold channel responses for conscious perception was a biphasic potential, the two largest peaks were selected from among all the identified post-stimulus local maxima. The two largest peaks were visually inspected against the channel mean of perceived trials voltage timecourses to confirm they represented true neural electrophysiologic peak events (i.e., not peak events of artifactual signals). Finally, the two selected peaks were sorted by latency relative to the stimulus onset and the channel peak latency was calculated as the time from stimulus onset to the first peak.

For the scalp ldEEG recordings, the Oz, Pz, and Cz channels were selected for peak latency analyses because in the hdEEG data set these channels displayed the strongest early and late scalp ERPs of interest, including the N100, VAN, and P3. As with the icEEG recordings, only perceived stimuli trials were considered for peak latency analyses. First, the mean scalp voltage across all perceived stimuli trials was computed within participants and the Oz, Pz, and Cz contacts. Next, group-level ldEEG data was obtained by averaging the participant-level mean scalp channel voltage within the Oz, Pz, and Cz channels. Peak latency analyses were focused on the following scalp ERPs found in the hdEEG data set: N100 (75-125ms), VAN (175-225ms), N2 (275-325ms), and P3 (350-650ms). While the N2 and P2 temporal windows were identical in the hdEEG analyses, because a frontal scalp ldEEG channel was not included in the latency analysis (due to contamination with blink and movement artifacts making peak detection unreliable), the P2 ERP was not considered. For each scalp ERP temporal window, all local maximums were obtained (MATLAB *findpeaks* function) from the absolute value of each channel. The local maximums were inspected in the original voltage space to find only those maxima that had the expected voltage polarity corresponding with the ERP: positive for P3 and negative for N100, VAN, and N2. The local maximums that were coherent with the polarity of the corresponding ERP were ranked by absolute amplitude and the sample with the largest absolute amplitude within a scalp ERP temporal window was identified as the peak for the current scalp ERP. In instances where no local maximums were found at a certain temporal window (e.g., the local maximums found did not match the polarity of the ERP), that scalp ERP peak was considered absent. Therefore, the total number of participant-scalp channel pairs for each scalp ERP was heterogeneous (Fig. 2F). Finally, for each participant, peak latencies from stimulus onset of each scalp ERP are computed on the participant-level voltage of the Oz, Pz, and Cz channels.

Latencies between the first peak of the thalamic ERP and each scalp ERP (N100, VAN, N2, and P3) were statistically compared with Wilcoxon rank sum tests (*p* < 0.05) across participants, Holm-Bonferroni-corrected for multiple comparisons.

## Data and Materials Availability

All experimental data and code used for the main text and supplementary materials are available at http://kronemer-blumenfeld-data.yale.edu <<Note: this URL will be open at the time of publication>>

## Supplementary Text

### Behavioral Performance

Behavioral performance was assessed on the Report Paradigm and the report (task-relevant) condition of the Report + No-Report Paradigm. The perception rate (percentage of seen stimuli versus the total number of presented stimuli) showed that both paradigms maintained perceptual threshold stimuli at a perception rate of approximately 50% across all study session conditions, including task screen backgrounds (movie versus noise; Report Paradigm), post-stimulus interval, study session neuroimaging condition, whether the task-relevant stimuli appeared first or second in a trial (Report + No-Report Paradigm), and the on-screen stimulus location (fig. S6, 7). Meanwhile, on average <5% of trials without a stimulus (blank trials) were reported as perceived in both the Report and Report + No-Report Paradigms (fig. S6D, 7D). Fully opaque stimuli in the Report + No-Report Paradigm were reported as perceived in on average ∼94% of trials (fig. S7D). These results suggest that overall participants were engaged with the task and understood the rules and demands for reporting their perceptions of task-relevant stimuli.

Moreover, similar behavioral performance between the Report Paradigm and task-relevant stimuli in the Report + No-Report Paradigm supports the conclusion that the addition of task-irrelevant stimuli did not alter how participants reported on the task-relevant stimuli.

The consistency of the behavioral performance across all study sessions and task conditions justified combining trials for subsequent analysis of concurrently recorded physiology from (1) within each paradigm condition (e.g., post-stimulus interval and stimulus location), (2) across each study session condition (i.e., hdEEG and fMRI), and (3) between the Report Paradigm and report condition of the Report + No-Report Paradigm.

#### Pupillometry and Eye Tracking

The mean temporal profiles of pupil diameter, blink, and microsaccade occurrence following stimulus presentation were similar when analyzed separately for hdEEG and fMRI data, except that microsaccade baseline rate was lower during fMRI because microsaccade detection was compromised by vibrations in the MRI bore (results not shown).

The mean pupil diameter, blink occurrence, and microsaccade occurrence for predicted perceived no-report trials reveal that as the classification score thresholds become more stringent (from 0 to 2 for labeling perceived trials; from 0 to −2 for labeling not perceived trials) the dynamics becomes more exaggerated. Specifically, pupil dilation and blink occurrence increases, and microsaccade occurrence decreases at more conservative score thresholds (fig. S9, warm color traces; blink and microsaccade dynamics not shown). A similar dynamic exists for the mean response from predicted not perceived trials, however, with more conservative score thresholds the opposite dynamic profile compared to the perceived condition emerges (fig. S9, cool color traces; blink and microsaccade dynamics not shown. These results suggest that the most confidently predicted perceived and not perceived trials are those with the most robust pupil, blink, and microsaccade dynamics, and, likewise, those most opposing to the competing perceptual class.

---

**Fig. S1.**
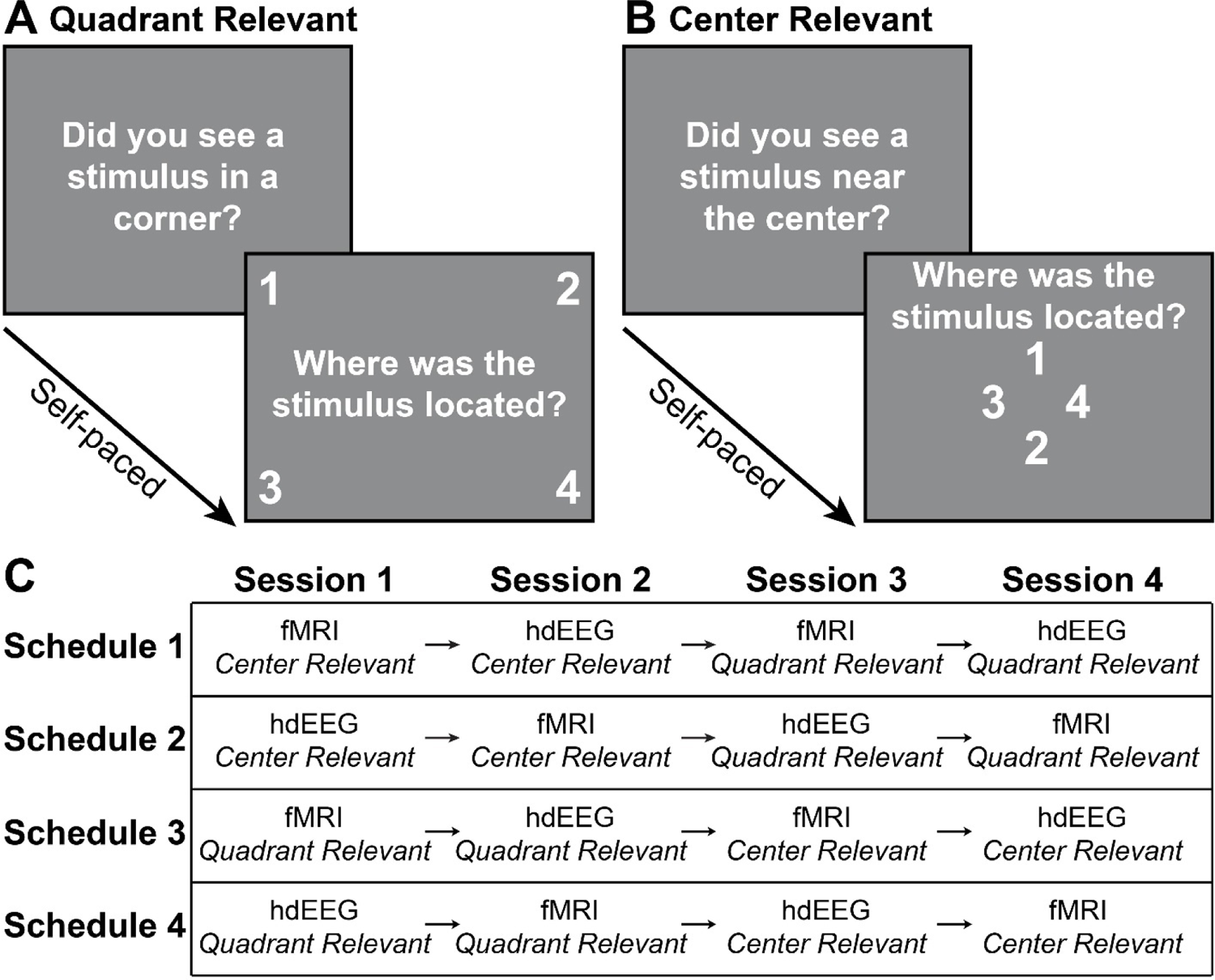
Behavioral paradigm response period task screens and study session sequence schedules. (**A**) Report and Report + No-Report Paradigms quadrant location set task-relevant perception and location questions response screens. In the Report Paradigm (Fig. 1A), the perception question read “Did you see a stimulus?” without the location specification (“in a corner”) because all stimuli appeared in a quadrant location. (**B**) Report + No-Report Paradigm center location set task-relevant perception and location questions response screens. (**C**) The Report + No-Report Paradigm four study session schedules dictating the sequence of the task-relevant and neuroimaging recording conditions. Each study session was completed on separate days of experimentation. The schedule type was counterbalanced across participants. See the *Visual Perception Paradigm* section for full details on the behavioral paradigms.

**Fig. S2.**
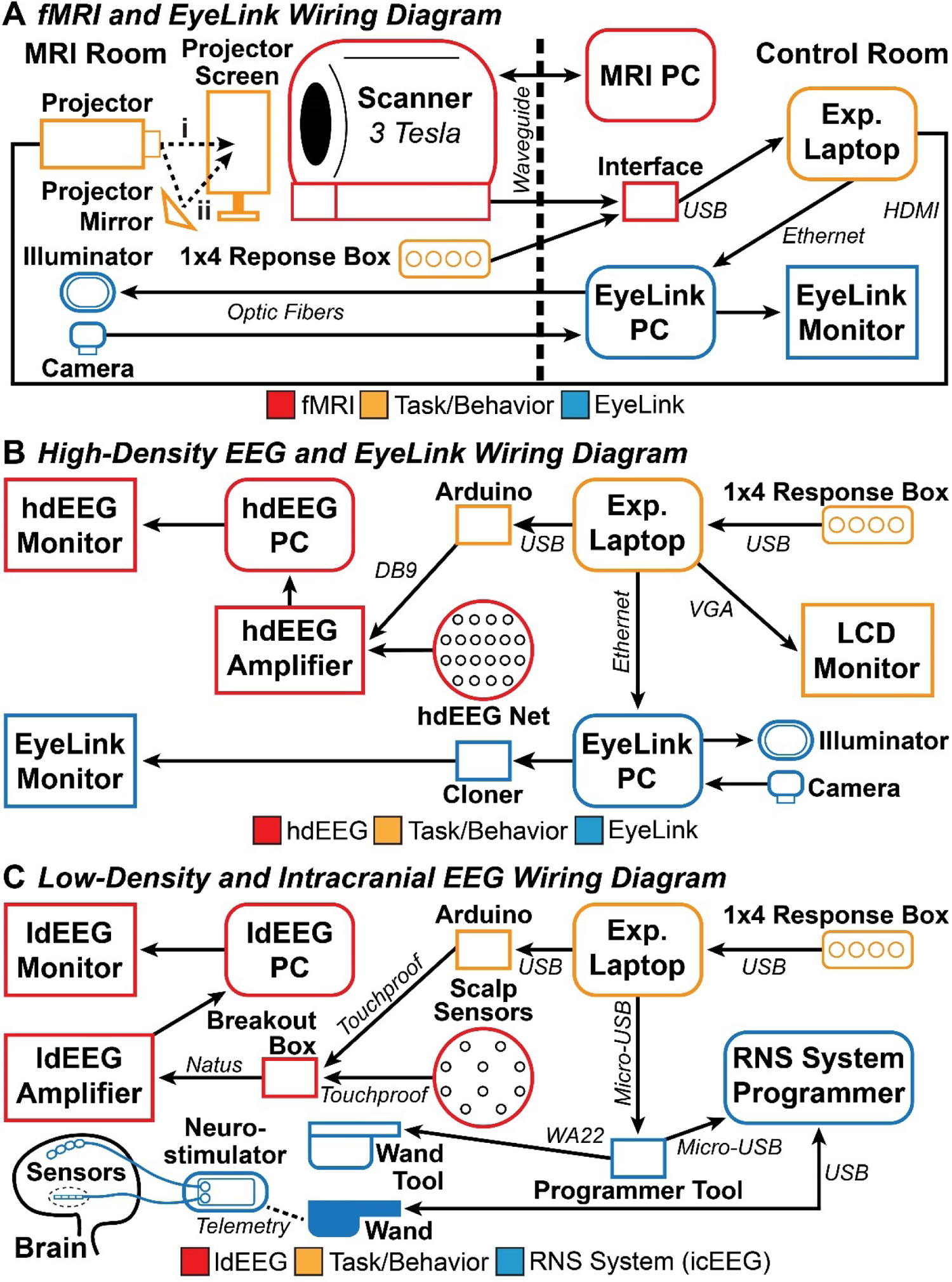
fMRI, hdEEG, ldEEG, icEEG, and EyeLink testing setup wiring diagrams. (**A**) Concurrent fMRI and EyeLink recording setup. Two behavioral task display projection approaches were used: (i) projecting onto the projector screen directly (used during acquisition of fMRI with the Report Paradigm), or (ii) projecting onto the projector screen via a mirror placed outside the MRI bore (used during acquisition of fMRI with the Report + No-Report Paradigm). (**B**) Concurrent hdEEG and EyeLink recording setup. (**C**) Concurrent ldEEG and icEEG recording with the RNS System (NeuroPace, Inc.). The icEEG recordings with the Natus NeuroWorks system (Natus, Inc.; participant 1 from Table S2) is not shown, but used a similar setup as for ldEEG. Experimental laptop (Exp. Laptop); High-density EEG (hdEEG); low-density EEG (ldEEG); intracranial EEG (icEEG). The *Behavior and Task Event Synchronization* section provides a full discrimination of setup and event synchronization approaches across each concurrent recording system.

**Fig. S3.**
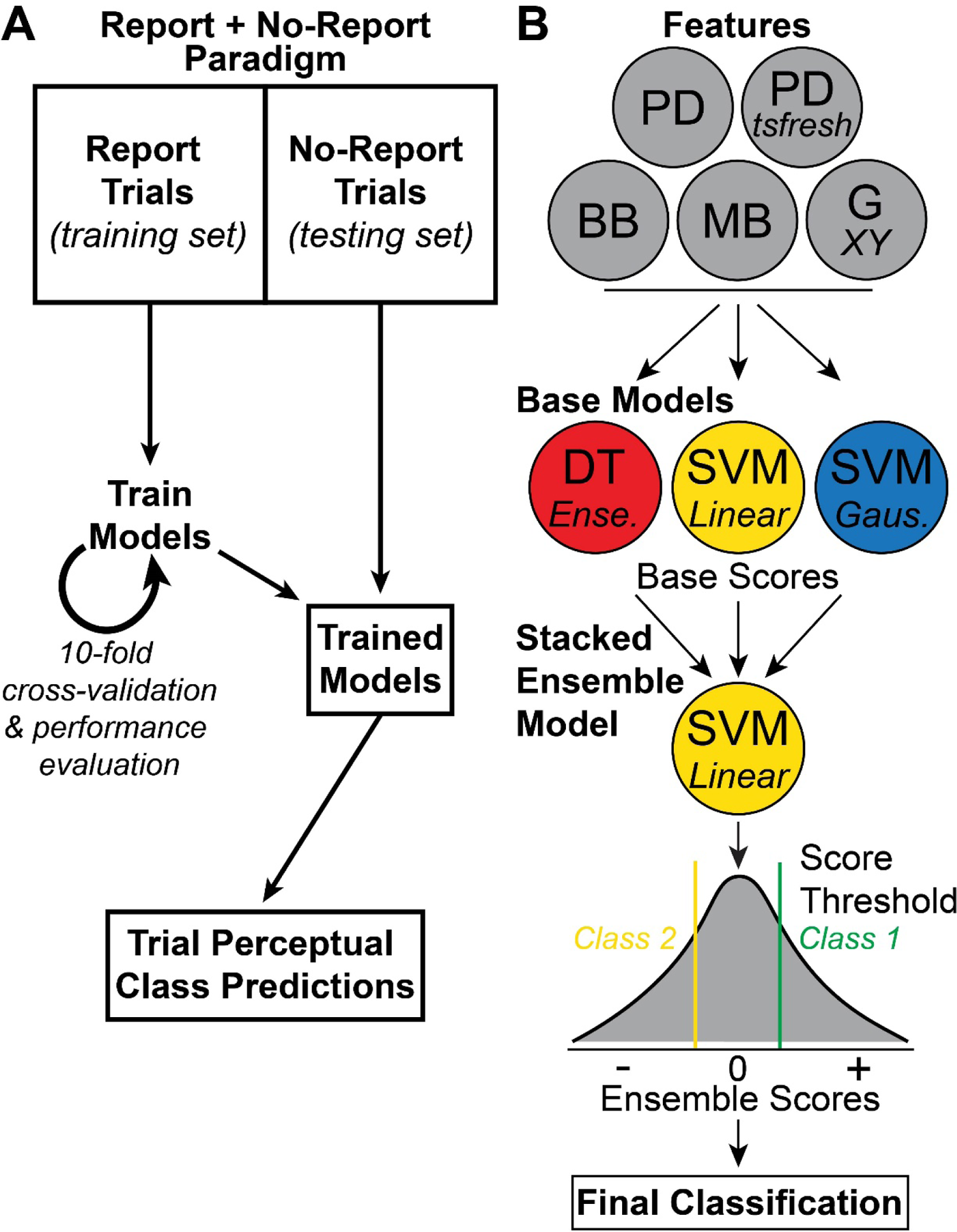
Covert measure of consciousness classification pipeline and model. (**A**) The training set is acquired from the Report (task-relevant) trials from the Report + No-Report Paradigm. The training data trained the base and stacked ensemble models with 10-fold cross-validation, and performance evaluation on predictions from these data were used to optimize classification performance. The trained models were applied to the testing set acquired from the no-report (task-irrelevant) trials of the Report + No-Report Paradigm. Trial perceptual class predictions are found for the no-report trials and implemented for subsequent analyses of physiological recordings. Training and testing sets were taken from the same task relevance and neuroimaging study session condition: (1) fMRI and quadrant location set task-relevant, (2) fMRI and center location set task-relevant, (3) hdEEG and quadrant location set task-relevant, and (4) hdEEG and center location set task-relevant. (**B**) Features from EyeLink data acquired in the Report + No-Report Paradigm and are trained/tested on three base models: (1) ensemble, boosted decision tree (DT), (2) linear kernel support vector machine (SVM), and (3) gaussian kernel SVM. The scores from the base models were used as features for a stacked ensemble model: linear kernel SVM. Scores from the ensemble model were used to determine the final trial classification of either perceived, not perceived, or unlabeled (due to classification uncertainty) by adjusting a score threshold to balance prediction accuracy and trial retention. See the *Covert Prediction of Conscious Perception – Machine Learning Pipeline* section for the full discrimination of features and classification pipeline. Pupil diameter (PD), blink binary (BB), microsaccade binary (MB), and gaze position (G) x and y components.

**Fig. S4.**
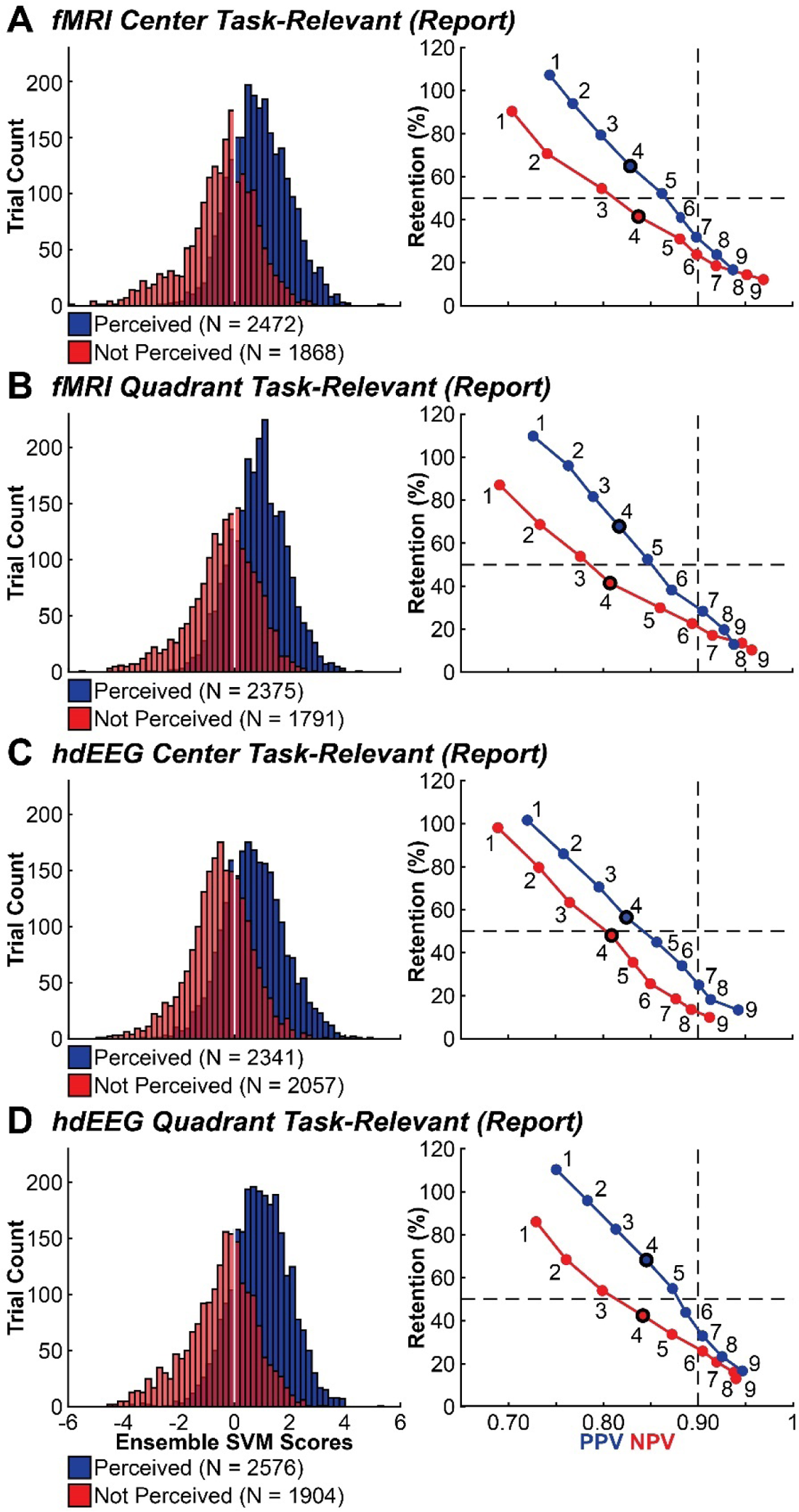
Report data classification performance from the Report + No-Report Paradigm. (**A**-**D**) Histograms show the distribution of stacked ensemble SVM scores (Fig. S3B) for all report trials within each study session condition, color coded by the true class label determined by overt report: (1) perceived (blue) and (2) not perceived (red). Line plots show positive predictive value (PPV; blue) and negative predictive value (NPV; red) for the perceived and not perceived classes, respectively, compared against trial retention (see the *Covert Prediction of Conscious Perception – Machine Learning Pipeline* section for PPV, NPV, and retention definitions) when manipulating the score threshold (Table S5). Each numbered point corresponds with the following scores for designating a perceived or not perceived trial: 1 = 0, 2 = -/+0.25, 3 = -/+0.5, 4 = -/+0.75, 5 = -/+1.0, 6 = -/+1.25, 7 = -/+1.5, 8 = -/+1.75, and 9 = -/+2.0. The ensemble SVM scores of −0.75 and 0.75 (points labeled “4” and highlighted with bold outlines) were selected for making the final determination of predicted perceived and not perceived, respectively. Justification for selecting these score thresholds for predicted perceived and not perceived trials is provided in the *Covert Prediction of Conscious Perception – Machine Learning Pipeline* section.

**Fig. S5.**
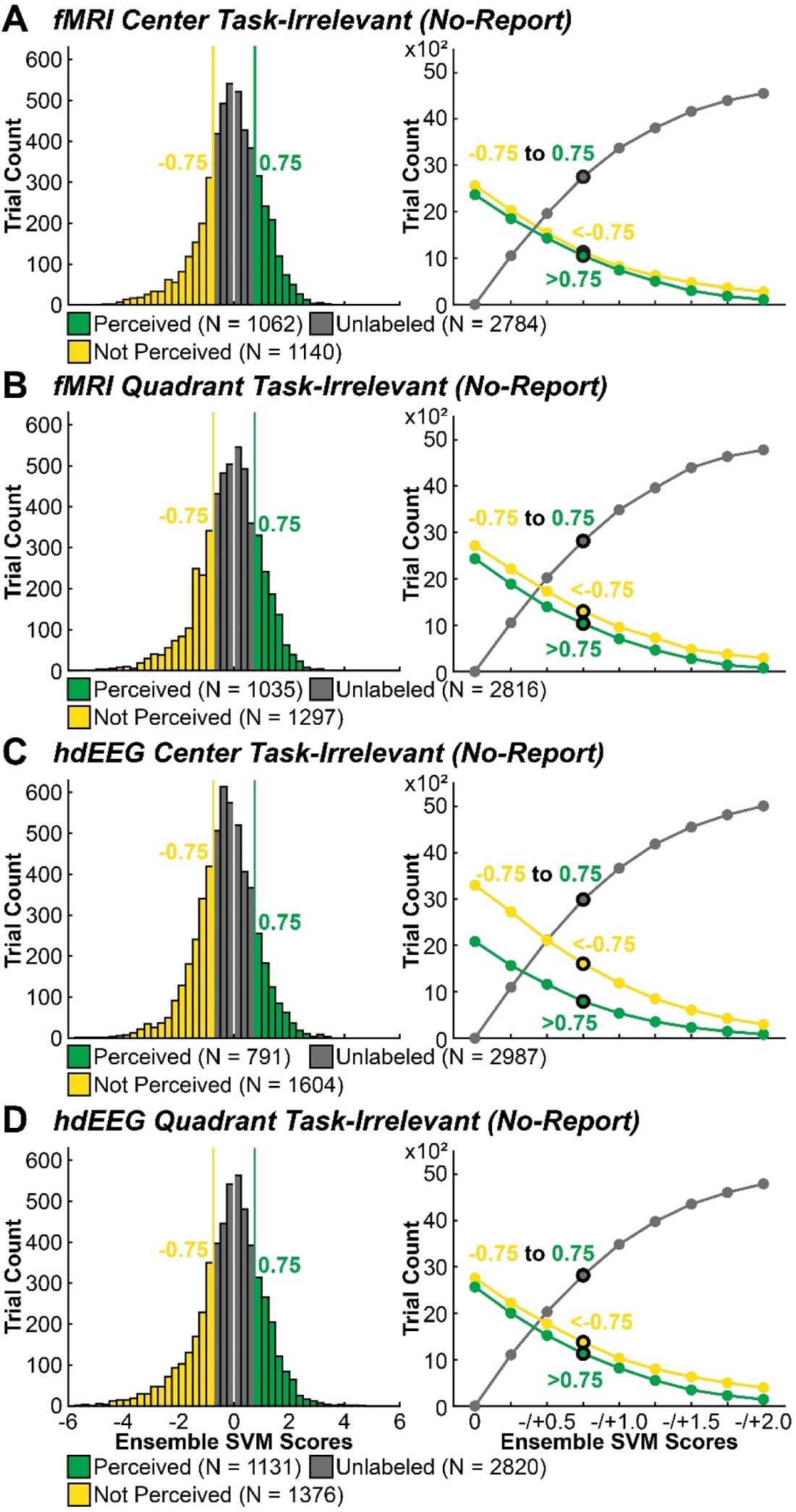
No-report (task-irrelevant) classification performance from the Report + No-Report Paradigm. (**A-D**) Histograms show the ensemble SVM score distributions for each study session condition. Portions of the score distribution are highlighted that exceed the perceived (green) and not perceived (yellow) score thresholds, 0.75 and −0.75, respectively. All trials that are found between the -/+0.75 thresholds remain unlabeled (grey). The line plots show the trial retention performance across ensemble SVM score thresholds of between 0 to -/+2.0 in increments of 0.25 for each study session condition and for the perceived (green), not perceived (yellow), and unlabeled (grey) conditions (Table S6). As score thresholds become more stringent the number of trials predicted perceived and not perceived decreases and unlabeled trials increase. The points corresponding with a score of -/+0.75 are highlighted with a bolded outline.

**Fig. S6.**
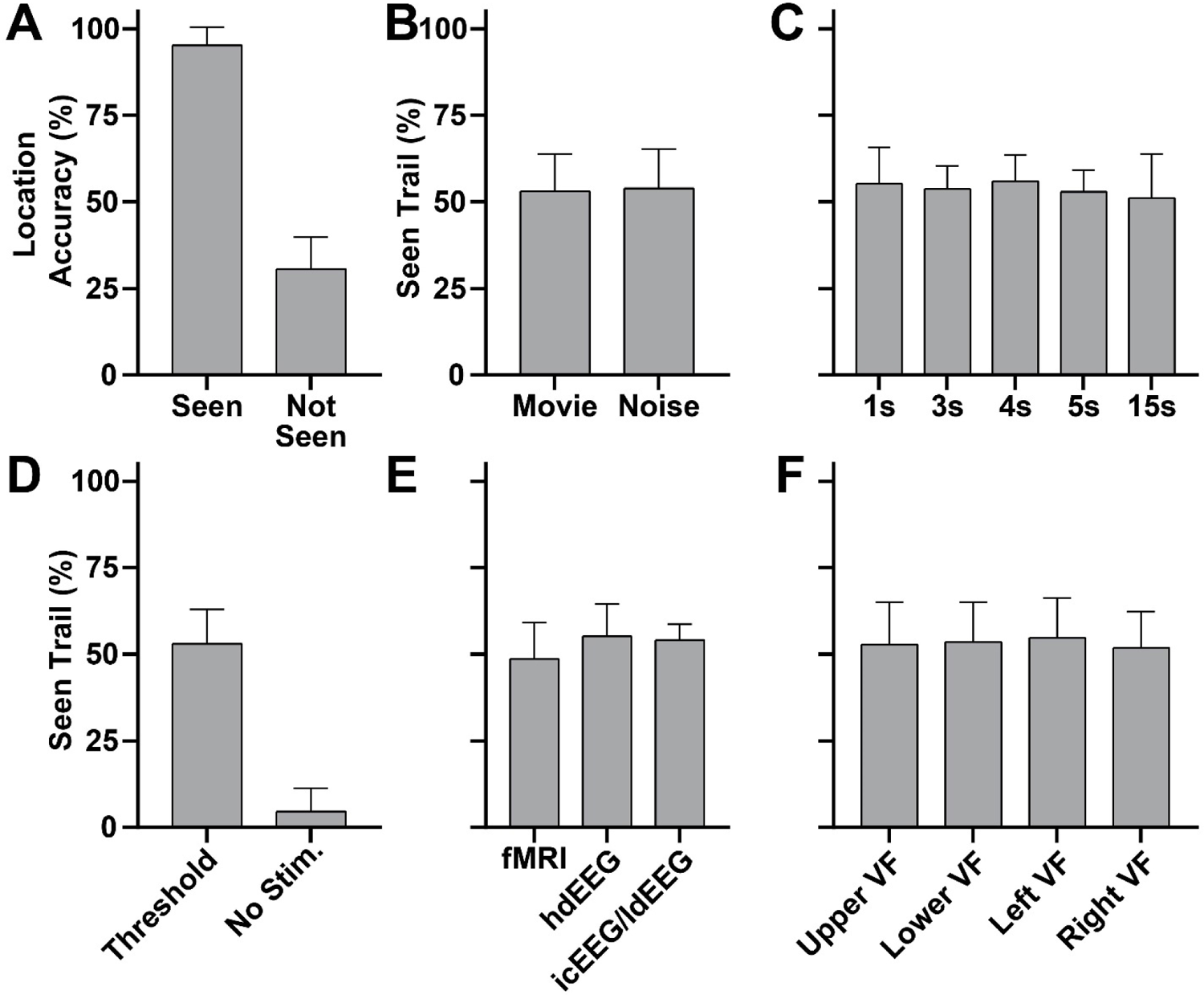
Report Paradigm behavioral performance. (**A**) Perceptual threshold stimuli location accuracy percentage for seen and not seen stimuli. (**B**) Percentage of seen perceptual threshold stimuli for movie and noise screen background conditions (see the *Visual Perception Paradigms* section). (**C**) Percentage of seen perceptual threshold stimuli for post-stimulus intervals of 1, 3, 4, 5, and 15 seconds. (**D**) Percentage of seen perceptual threshold stimuli and no stimuli (blank) trials. (**E**) Percentage of seen perceptual threshold stimuli for fMRI, high-density EEG (hdEEG), and intracranial EEG (icEEG)/low-density EEG (ldEEG) study sessions. (**F**) Percentage of seen perceptual threshold stimuli by location on screen: (1) upper visual field (VF), (2) lower VF, (3) left VF, and (4) right VF. Error bars show standard deviation. fMRI participants N = 35. hdEEG participants N = 59. icEEG/ldEEG participants N = 7 (one participant had icEEG recordings without ldEEG; participant 1 in Table S2).

**Fig. S7.**
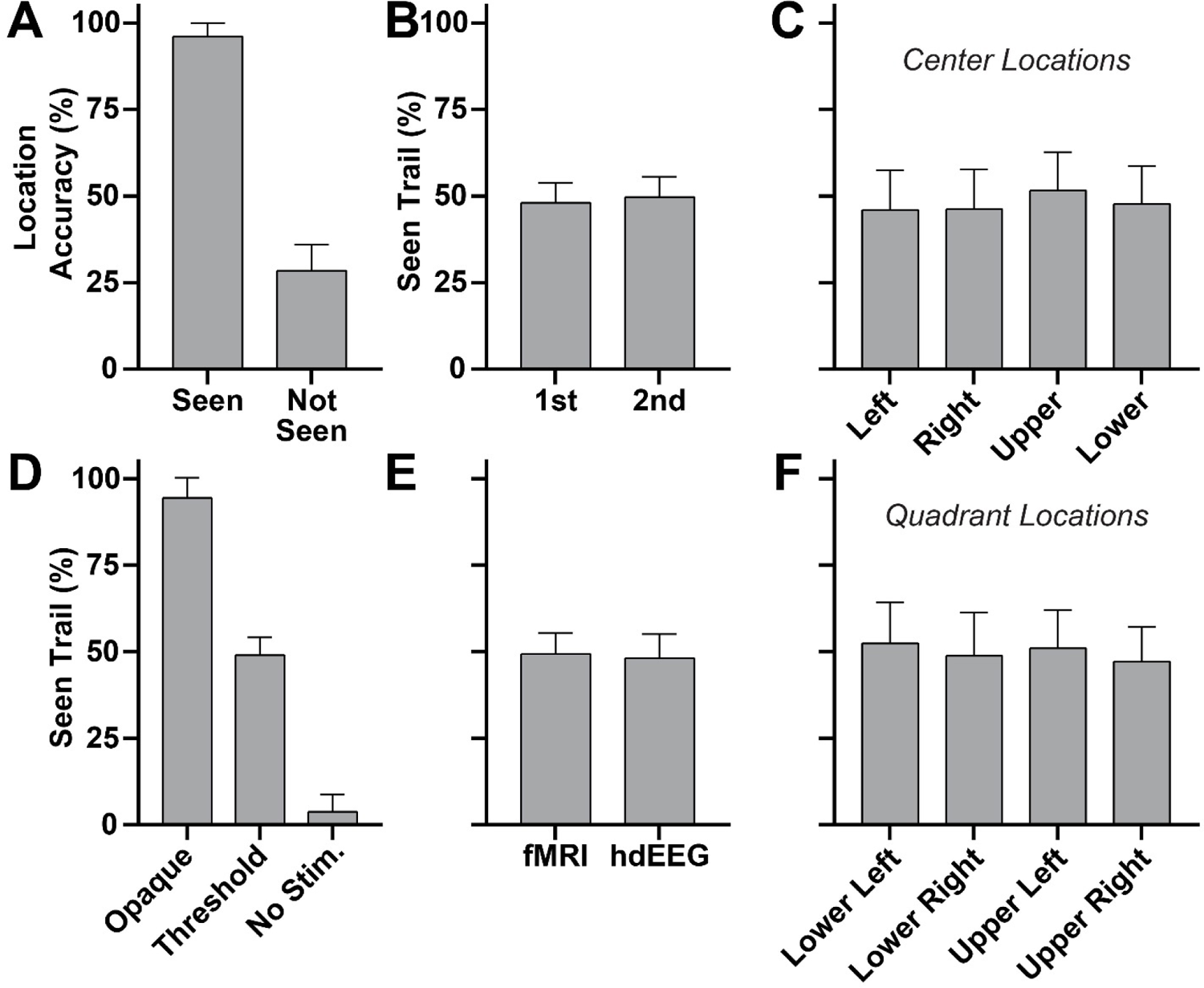
Report + No-Report Paradigm behavioral performance from the report (task-relevant) condition. (**A**) Perceptual threshold stimuli location accuracy percentage for seen and not seen stimuli. (**B**) Percentage of seen perceptual threshold stimuli for trials when the task-relevant stimulus appeared first or second (see the *Visual Perception Paradigms* section). (**C**) Percentage of seen perceptual threshold stimuli by center locations: (1) left, (2) right, (3) upper, and (4) lower positions relative to the central fixation cross. (**D**) Percentage of seen fully opaque, perceptual threshold, and no stimuli (blank) trials. (**E**) Percentage of seen perceptual threshold stimuli for fMRI and high-density EEG (hdEEG) study sessions. (**F**) Percentage of seen perceptual threshold stimuli by quadrant locations: (1) lower left, (2) lower right, (3) upper left, and (4) upper right. Error bars show standard deviation. fMRI participants N = 65. hdEEG participants N = 65.

**Fig. S8.**
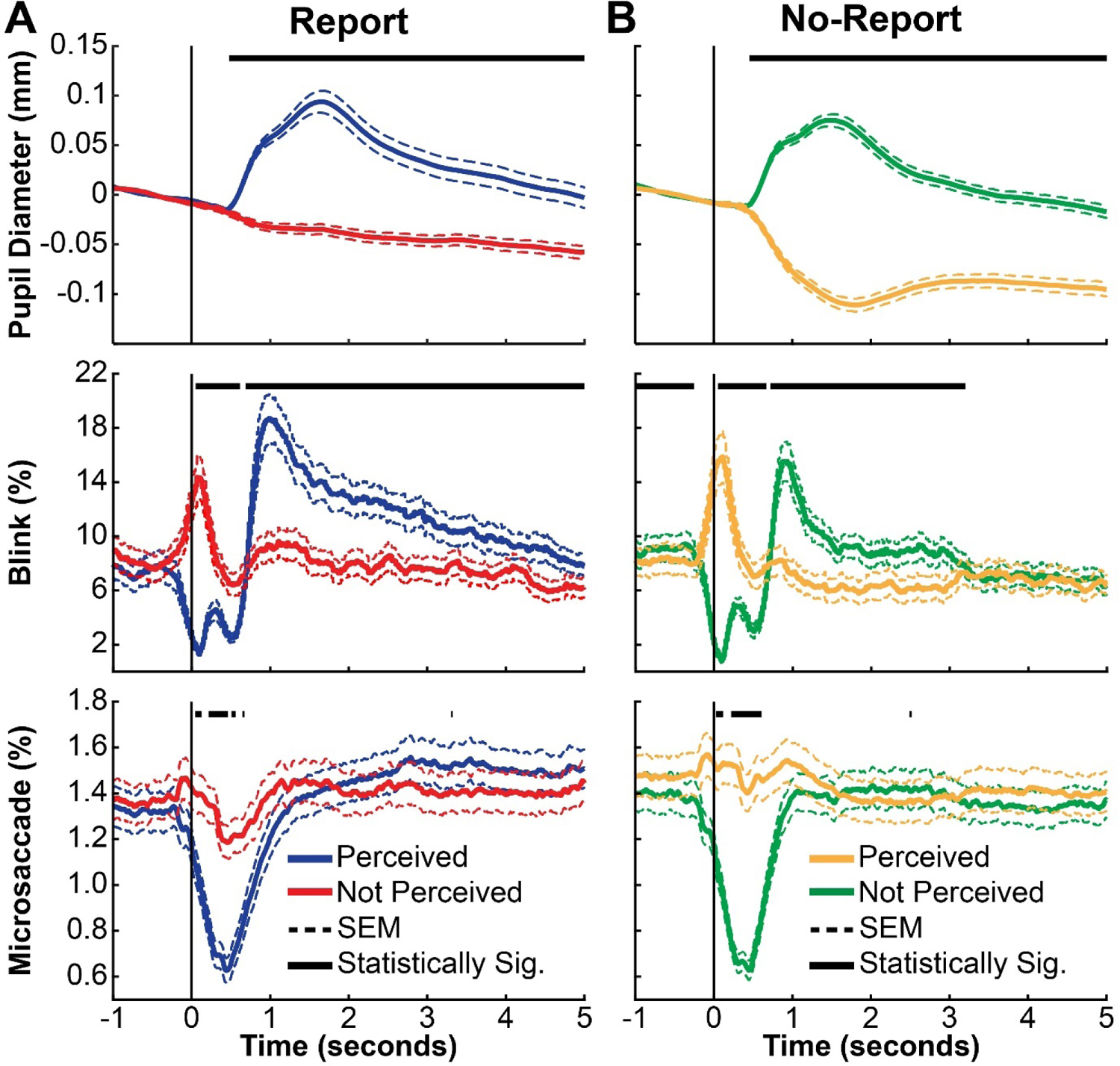
Pupillometry and eye tracking dynamics for perceived and not perceived stimuli, without removing trials that contain eye blinks during the stimulus. (**A**) Report and (**B**) no-report perceived and not perceived stimuli from the Report + No-Report Paradigm (see the *Visual Perception Paradigms* section). The pupil diameter (mm), blink occurrence (%), and microsaccade occurrence (%) are identical to those presented in Fig. 1C and D except in the current figure trials with a blink event during stimuli presentation period (50ms) are not excluded. Stimulus onset was at time = 0. Classification score threshold for (**B**) was 0. Statistically significant (Statistically Sig.) samples by temporal cluster-based permutation tests (*p* < 0.05) of perceived versus not perceived conditions are shown. Trials are combined across study session and task conditions. Standard error of the mean (SEM). Participants N = 68.

**Fig. S9.**
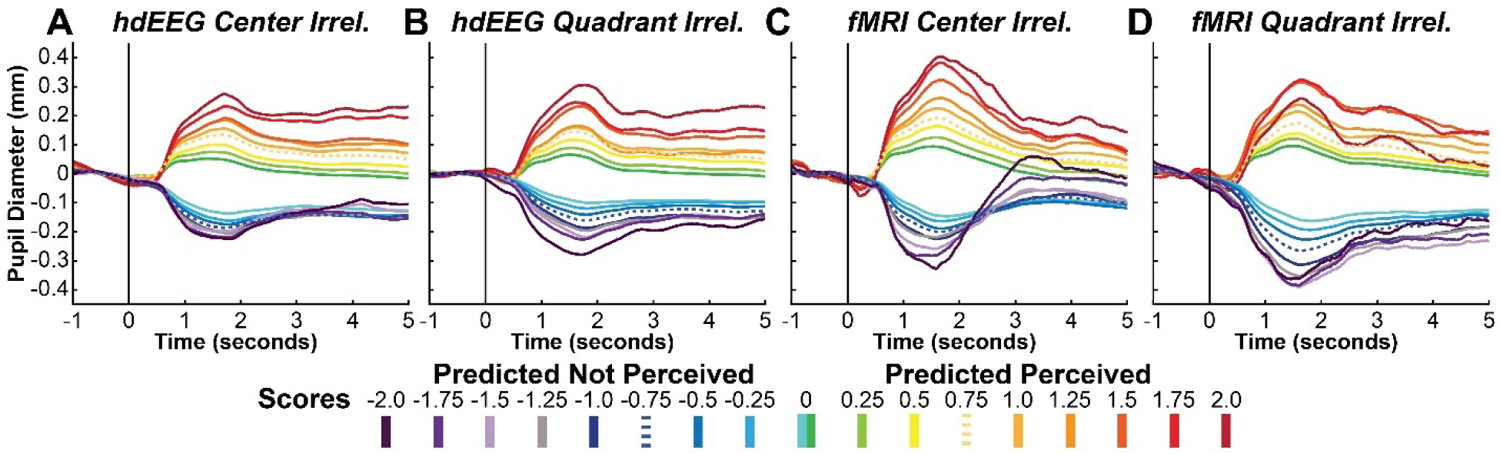
No-report predicted perceived and not perceived pupil diameter (mm) dynamics across classification score thresholds. All task-irrelevant study session conditions (no-report trials) of the Report + No-Report Paradigm are shown independently: (**A**) high-density EEG (hdEEG) center location task-irrelevant (Irrel.), (**B**) hdEEG quadrant location task-irrelevant, (**C**) fMRI center location task-irrelevant, and (**D**) fMRI quadrant location task-irrelevant. Each color trace corresponds with the mean of trials predicted as perceived (warm colors) or not perceived (cool colors) according to a specified score threshold (see the *Covert Prediction of Conscious Perception – Machine Learning Pipeline* section for the full discrimination of classification scores). The dotted trace shows the timecourses corresponding to a score threshold of 0.75 and −0.75 for predicted perceived and not perceived trials, respectively, the same thresholds used for the main results found in Fig. 2, 3, and 4. The number of participants contributing to the mean timecourses ranged between 60 to 28 participants, from the least to most stringent score thresholds. Data not shown for blink and microsaccade timecourses.

**Fig. S10.**
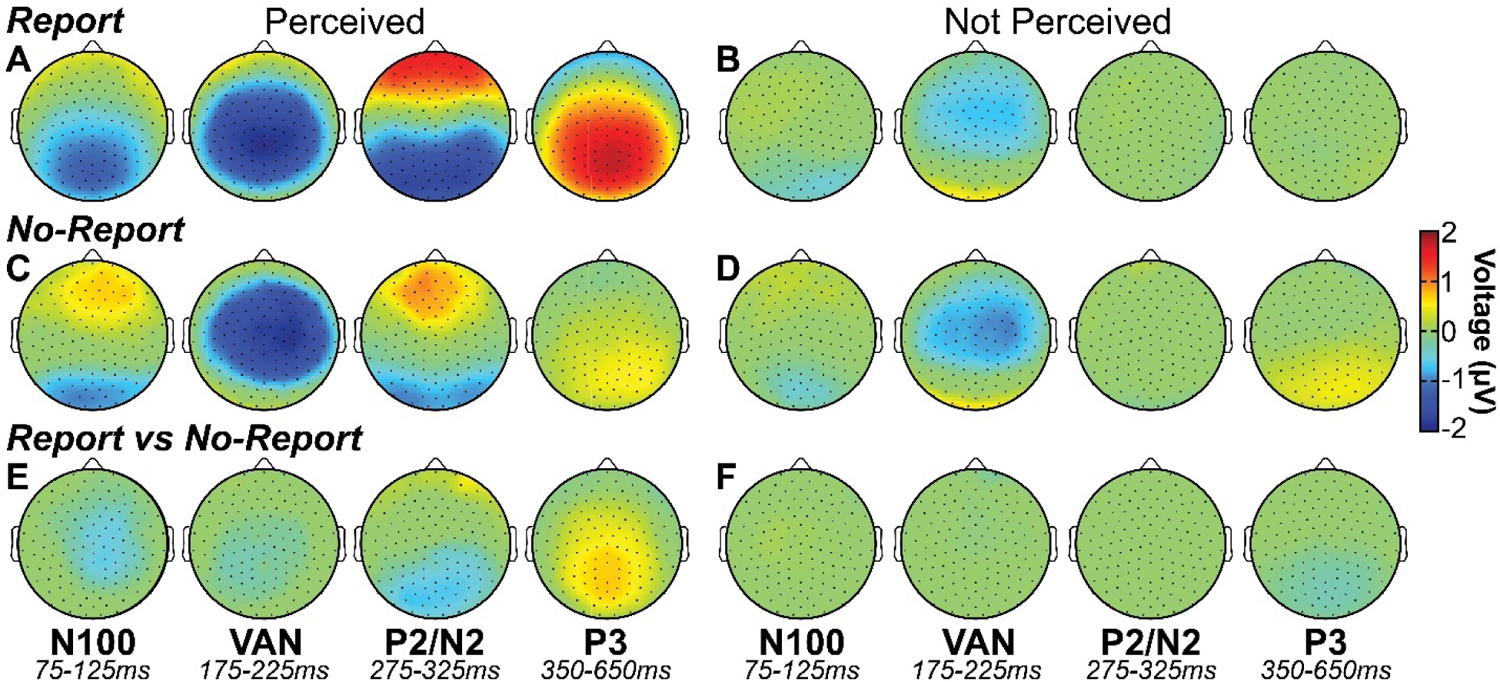
Report and no-report data perceived and not perceived high-density EEG. The mean voltage is shown for all statistically significant channels and samples found by cluster-based permutation tests (*p* < 0.05) in four time windows corresponding with the event-related potentials (ERPs) N100 (75-125ms), visual awareness negativity (VAN; 175-225ms), P2/N2 (275-325ms), and P3 (350-650ms). Data are for (**A**) report perceived, (**B**) report not perceived, (**C**) no-report perceived, (**D**) no-report not perceived, (**E**) perceived report versus no-report, and (**F**) not perceived report versus no-report. Report healthy participants N = 122. No-report healthy participants N = 65 (2 not perceived subject data sets were excluded because at the prediction score threshold of −0.75 there were no predicted not perceived trials; see the *Covert Prediction of Conscious Perception – Machine Learning Pipeline* section).

**Fig. S11.**
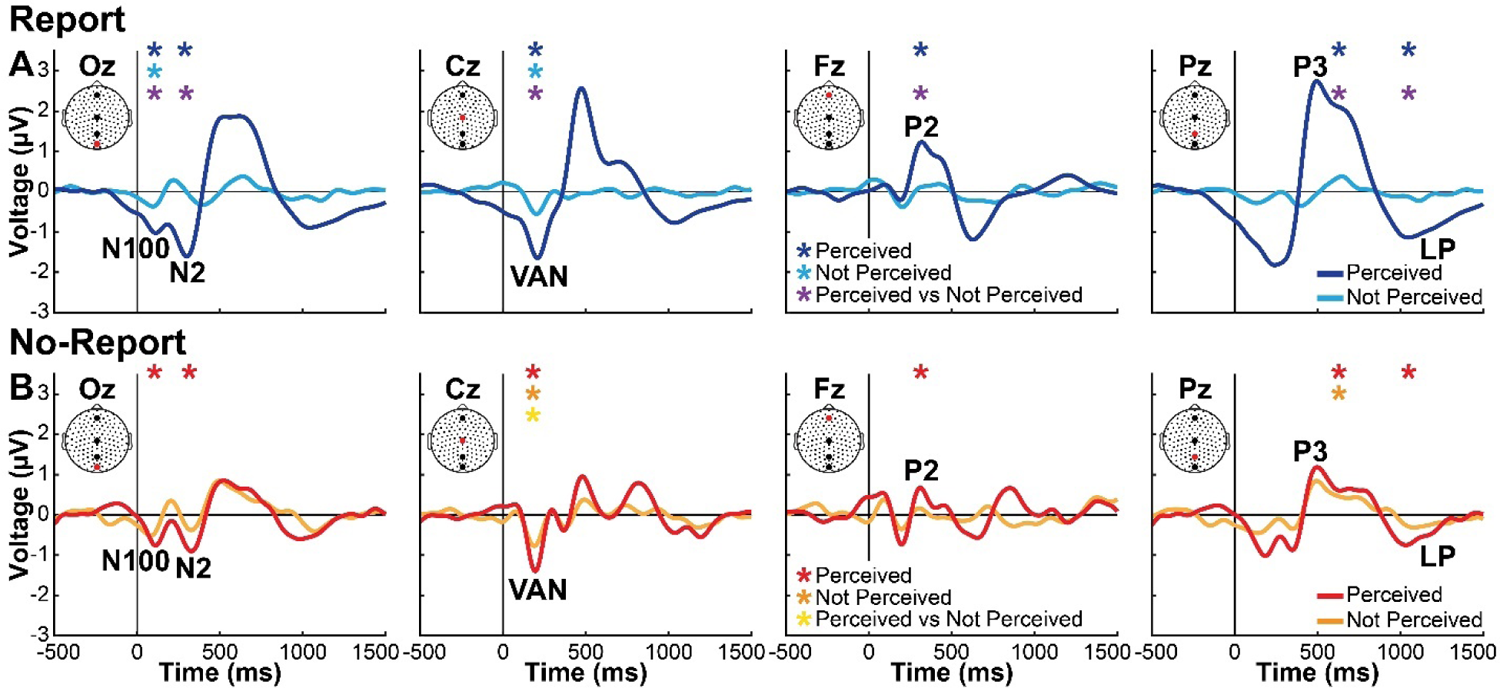
Voltage timecourses from scalp contacts Oz, Cz, Fz, and Pz for report and no-report data. (**A**) Report data perceived (dark blue) and not perceived (light blue), and (**B**) no-report data perceived (red) and not-perceived (orange). Asterisks indicate statistically significant perceived event-related potentials by cluster-based permutation tests (*p* < 0.05) of the perceived (dark blue/red asterisks), not perceived (light blue/orange asterisks), and perceived versus not perceived (purple/yellow asterisks) statistical comparisons. Location of Oz, Cz, Fz, and Pz electrodes are shown by red dots on insets. Labels show peaks of event-related potentials, including N100, N2, P2, P3, visual awareness negativity (VAN), and late potentials (LP). Report healthy participants N = 122. No-report healthy participants N = 65 (2 not perceived subject data sets were excluded because at the prediction score threshold of −0.75 there were no predicted not perceived trials; see the *Covert Prediction of Conscious Perception – Machine Learning Pipeline* section).

**Fig. S12.**
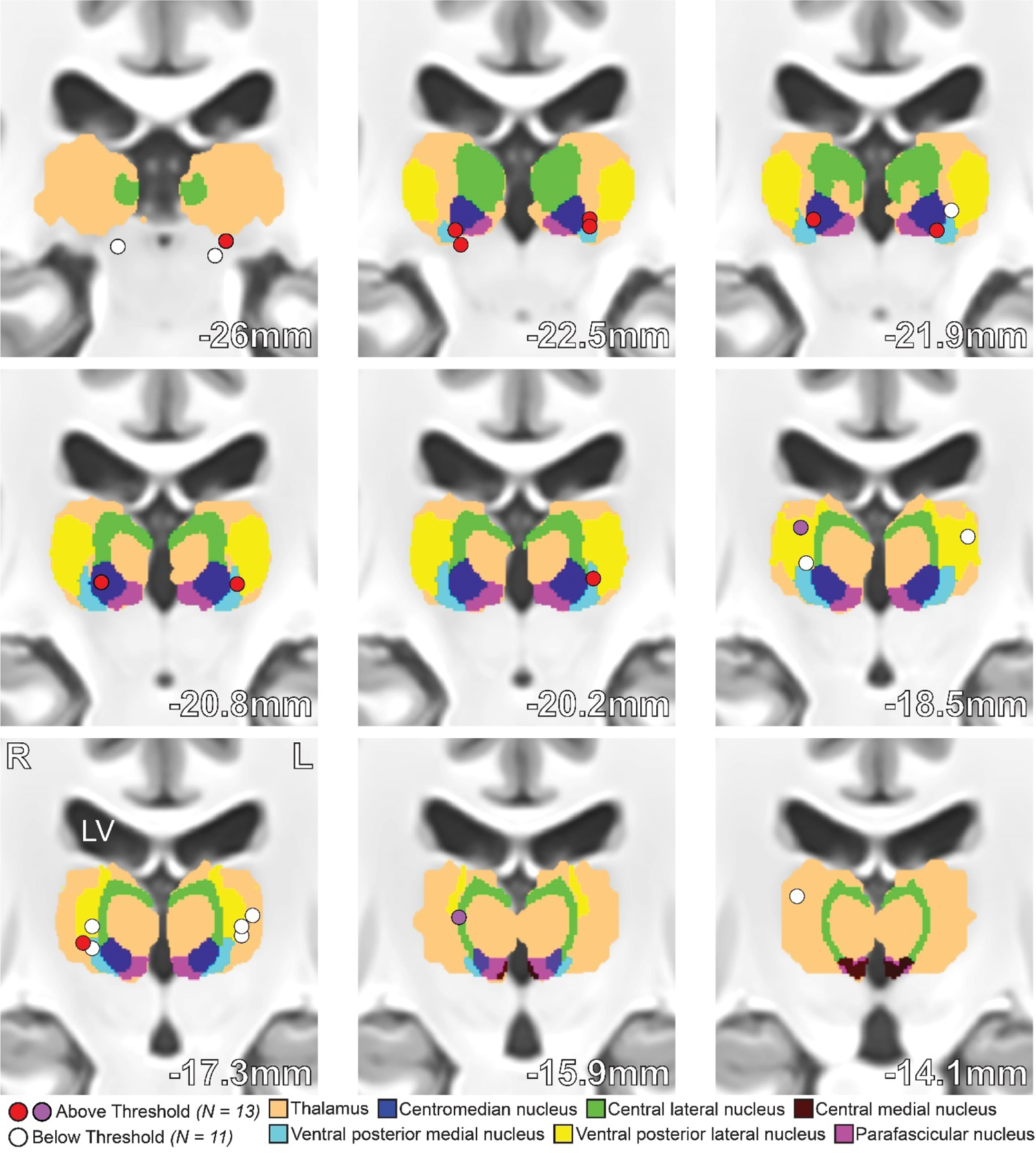
Location of intracranial EEG bipolar channels. Channels are colored coded as above (red and purple) or below threshold (white). See the *Intracranial EEG (icEEG)* section for details on finding bipolar channel locations from depth contacts and the channel selection procedure that designates above and below threshold channels. The channels drawn in purple identify the participant with icEEG recordings made with the Natus NeuroWorks system and without concurrent low-density EEG recordings (participant 1 in Table S2). Overlay of the thalamus and thalamic subregions are taken from a voxelized version of Morel atlas (*43, 44*). Two-dimensional channel locations are shown in Fig. 2D. Exact channel locations in MNI coordinate space is listed in Table S3. Patient participants N = 7.

**Fig. S13.**
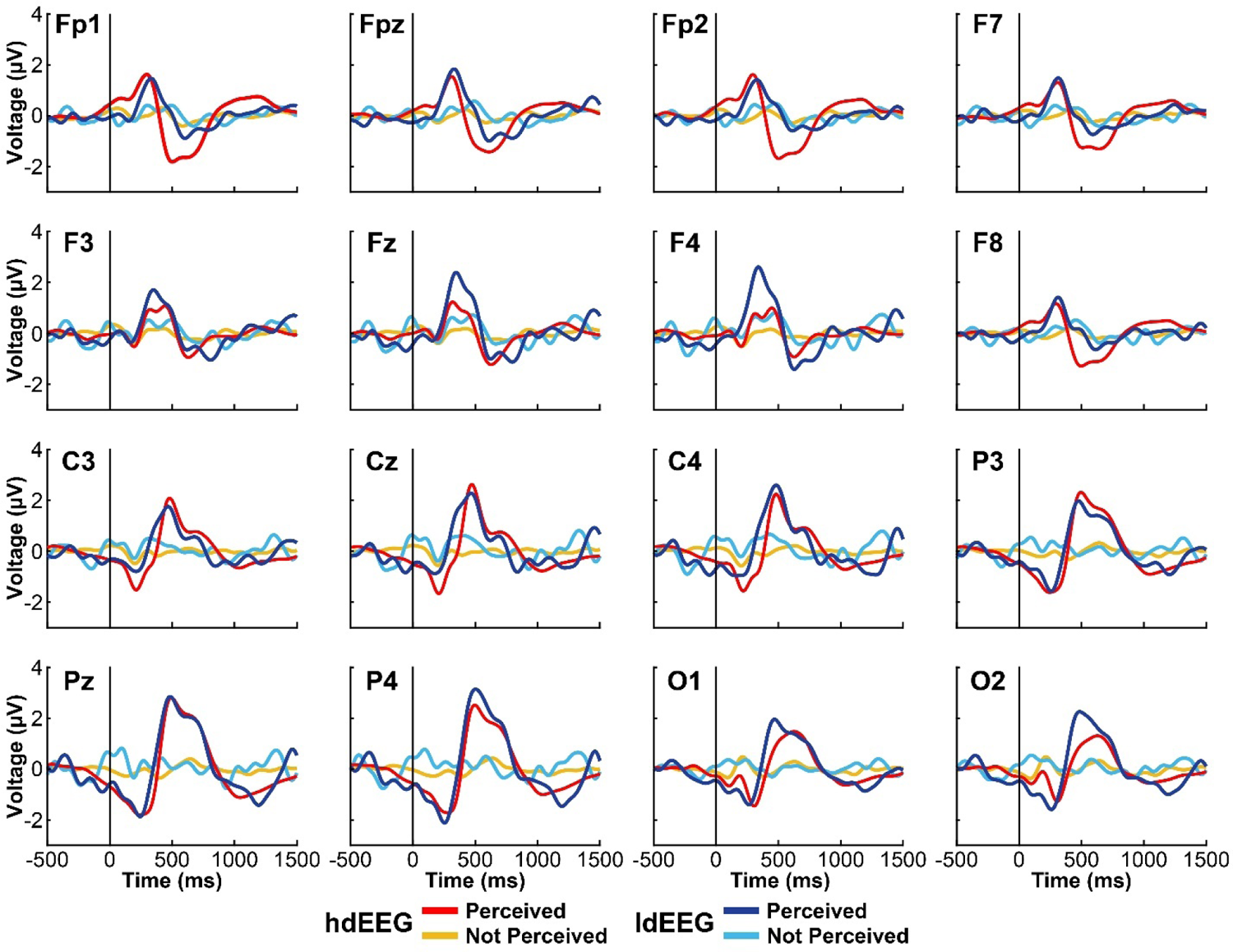
Comparison of voltage timecourses from low and high-density EEG recordings across shared scalp contacts. Perceived (red/dark blue) and not perceived (orange/light blue). Stimulus onset was at time = 0. Standard international 10-20 electrode locations are listed at top left of each panel. High-density EEG (hdEEG); low-density EEG (ldEEG). hdEEG participants N = 122. ldEEG participants N = 6.

**Fig. S14.**
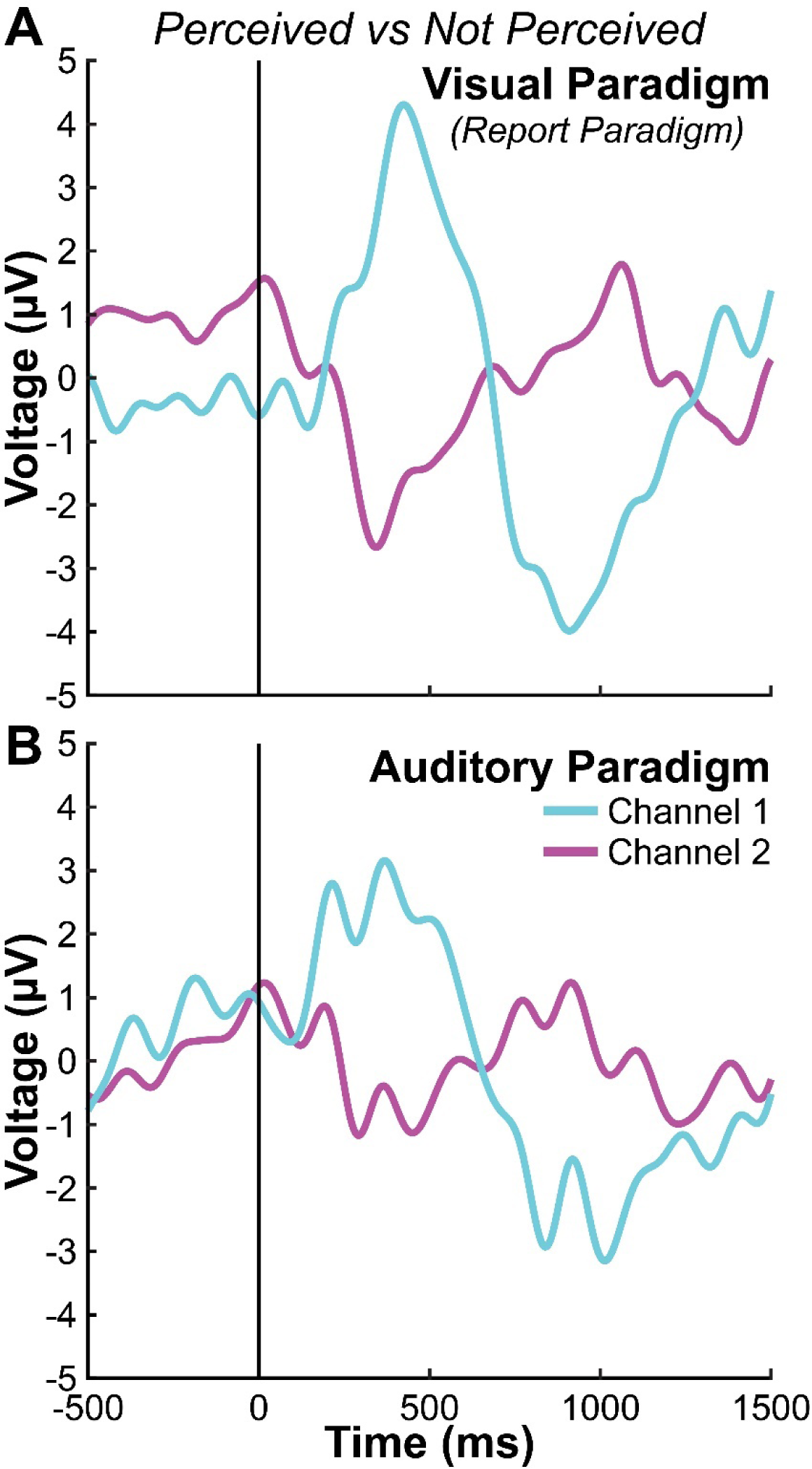
Thalamic awareness potential for conscious visual versus auditory paradigms. Perceived minus not perceived trials mean voltage timecourses for the patient participant (participant 1 in Table S2) who completed both the (**A**) Report Paradigm (“Visual Paradigm”) and (**B**) a similar auditory perceptual threshold paradigm (“Auditory Paradigm”) (*12*). Two identical thalamic bipolar channels (blue and purple) are compared between the visual and auditory paradigms. The bipolar channels drawn in purple in Fig. 2D and Fig. S12 show the channel locations for the current figure. Visual Paradigm: perceived = 353 trials; not perceived = 188 trials. Auditory Paradigm: perceived = 362 trials; not perceived = 245 trials.

**Fig. S15.**
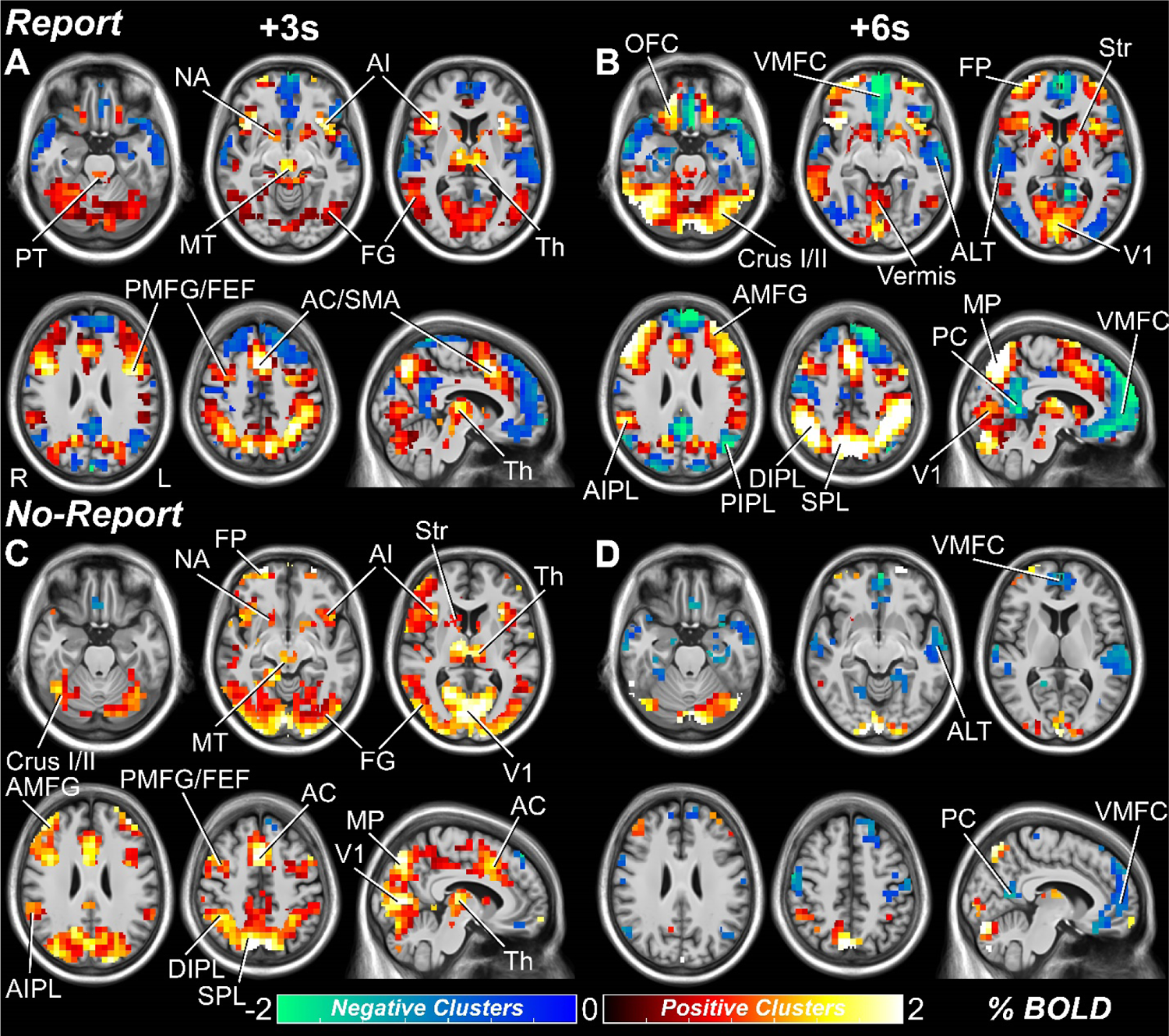
Whole brain percent change BOLD maps for perceived versus not perceived stimuli with and without overt report. (**A**, **B**) Report and (**C**, **D**) no-report statistically significant voxels from cluster-based permutation tests (*p* < 0.05) at 3 and 6 seconds post-stimulus presentation. Statistically significant positive and negative voxel percent change BOLD values are shown in warm and cool colors, respectively. Anatomical regions are labeled at their peak response between the 3 and 6-second time points. Pons tegmentum (PT), midbrain tegmentum (MT), thalamus (Th), nucleus accumbens (NA), striatum (Str), anterior insula/claustrum (AI), anterior cingulate (AC), supplementary motor area (SMA), primary visual cortex (V1), fusiform gyrus (FG), anterior middle frontal gyrus (AMFG), posterior middle frontal gyrus (PMFG), frontal eye fields (FEF), frontal pole (FP), orbital frontal cortex (OFC), ventral medial prefrontal cortex (VMFC), anterior inferior parietal lobule (AIPL), posterior inferior parietal lobule (PIPL), dorsal inferior parietal lobule (DIPL), superior parietal lobule (SPL), medial parietal cortex (MP), posterior cingulate/precuneus (PC), and anterolateral temporal cortex (ALT). Report healthy participants N = 99. No-report healthy participants N = 65 (one perceived subject data set was excluded because at the prediction score threshold of 0.75 there were no predicted perceived trials; see the *Covert Prediction of Conscious Perception – Machine Learning Pipeline* section).

**Fig. S16.**
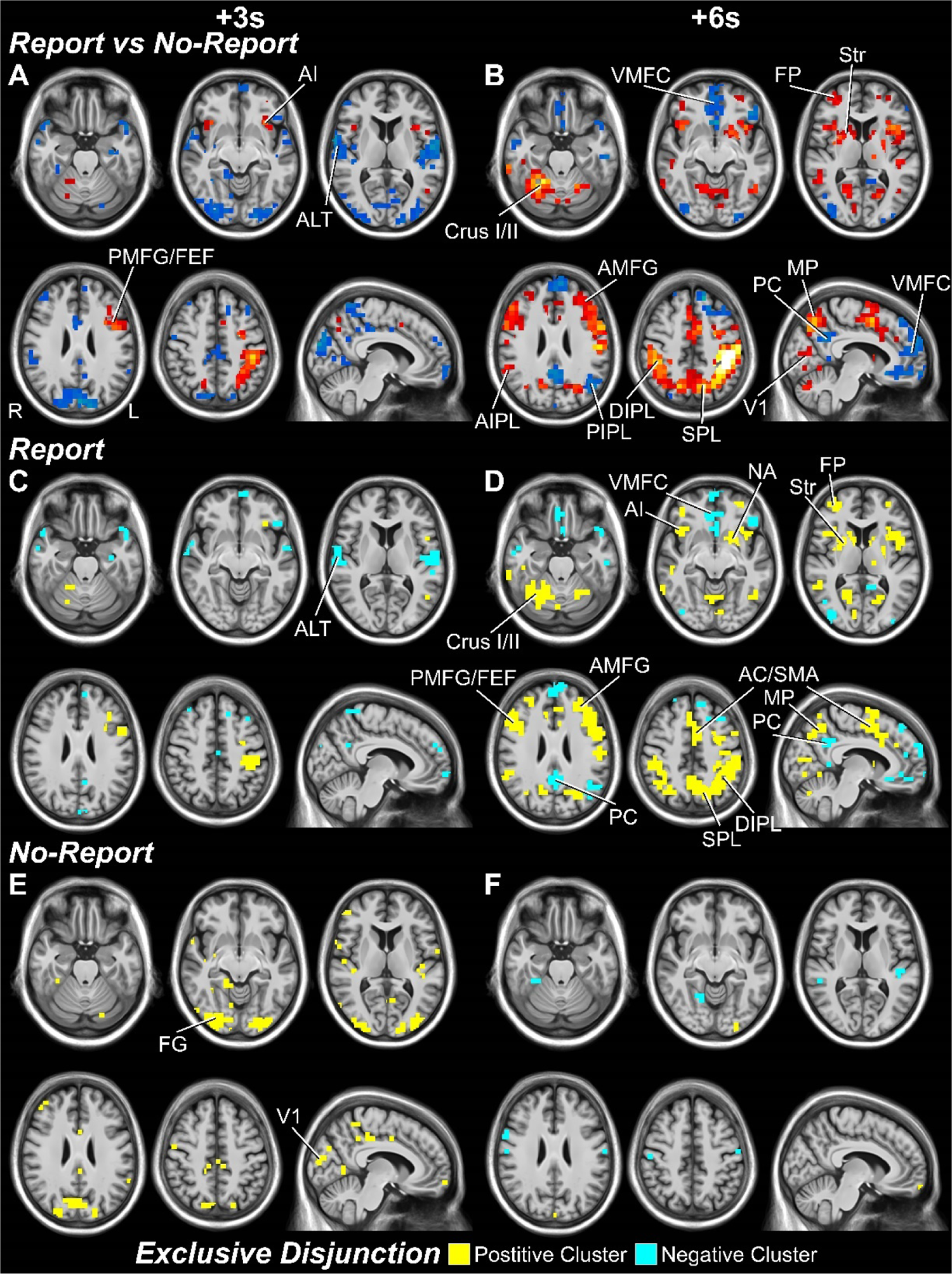
Report and no-report difference analyses. Analyses are for perceived minus not perceived stimuli, comparing report versus no-report data. (**A**) Report versus no-report statistically significant voxels from cluster-based permutation tests (*p* < 0.05) at 3 and 6 seconds post-stimulus presentation. Statistically significant positive and negative voxel *t*-values are shown in warm and cool colors, respectively. Anatomical regions are labeled at their peak response between the 3 and 6-second time points. All 20 seconds pre and post-stimulus time points for the report versus no-report comparison is shown in Slide S3. (**C**, **D**) regions significant only for report data. (**E**, **F**) regions significant only for no-report data (see *fMRI Spatiotemporal Conjunction and Exclusive Disjunction Analyses* section). Exclusive disjunction voxels are color coded by membership in either regions of significant fMRI increases (yellow) or decreases (blue) determined by cluster-based permutation tests (*p* < 0.05). All 20 seconds pre and post-stimulus time points for exclusive disjunction analyses are shown in Slides S5 and S6. Anatomical regions are labeled at their peak spatial extend between the 3 and 6-second time points. Nucleus accumbens (NA), striatum (Str), anterior insula/claustrum (AI), anterior cingulate (AC), supplementary motor area (SMA), primary visual cortex (V1), fusiform gyrus (FG), posterior middle frontal gyrus (PMFG), frontal eye fields (FEF), frontal pole (FP), anterior middle frontal gyrus (AMFG), ventral medial prefrontal cortex (VMFC), dorsal inferior parietal lobule (DIPL), posterior inferior parietal lobule (PIPL), superior parietal lobule (SPL), medial parietal cortex (MP), and posterior cingulate/precuneus (PC), anterolateral temporal cortex (ALT). Report healthy participants N = 99. No-report healthy participants N = 65 (one perceived subject data set was excluded because at the prediction score threshold of 0.75 there were no predicted perceived trials; see the *Covert Prediction of Conscious Perception – Machine Learning Pipeline* section).

**Fig. S17.**
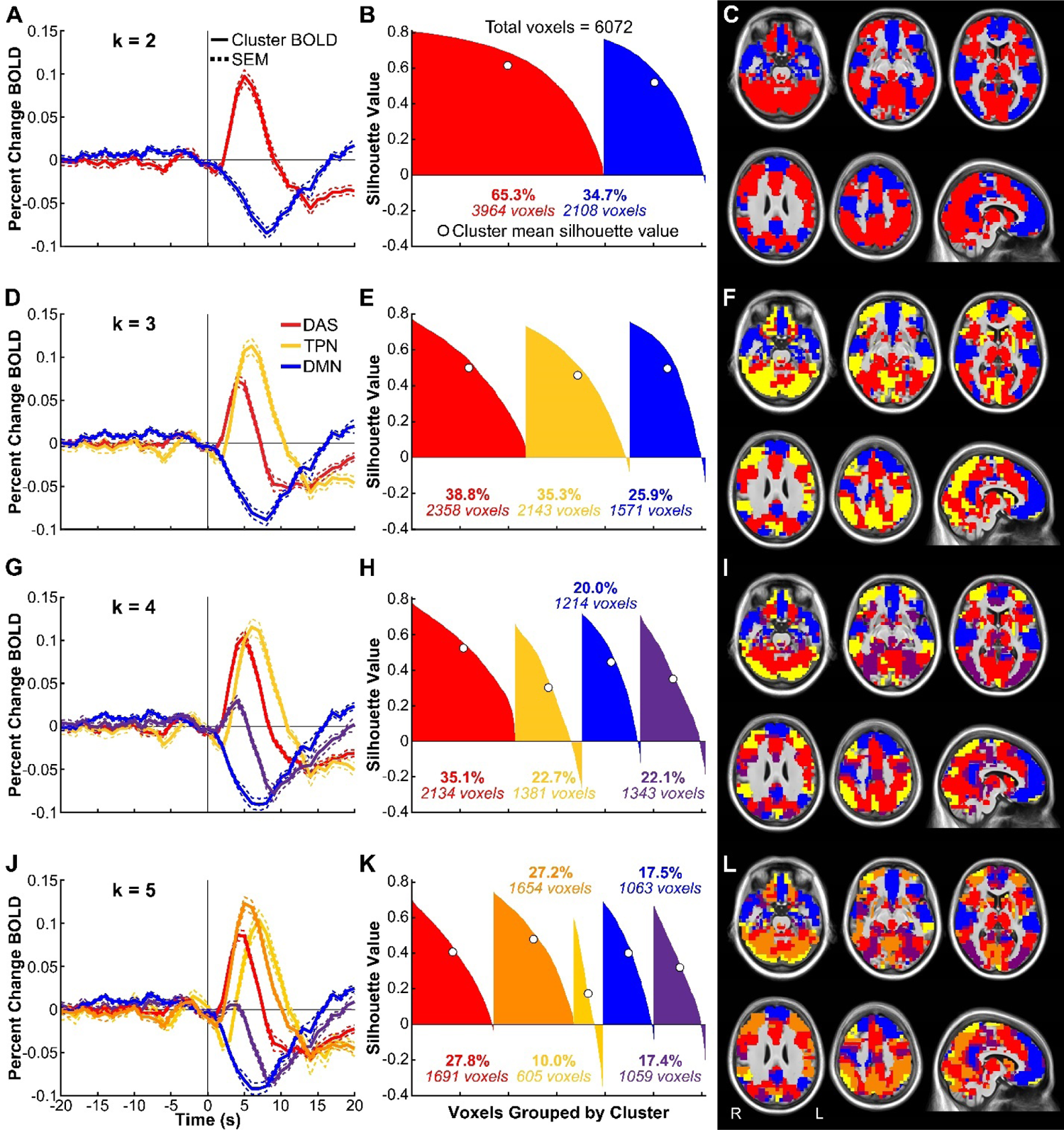
Data-driven whole brain k-means clustering of percent change BOLD signals for report perceived versus not perceived timecourses. Only voxels that were found statistically significant by cluster-based permutation analysis (*p* < 0.05) within the first 10 seconds post-stimulus were included in k-means clustering (see the *fMRI Anatomical Clustering* section for full details on the clustering method). (**A**, **D**, **G**, **J**) Mean percent change blood-oxygen-dependent-signal (BOLD) signal time courses are shown for all voxels within each k-means cluster for (**A**) k = 2, (**D**) k = 3, (**G**) k = 4, and (**J**) k = 5. Dotted lines show standard error of the mean (SEM). K = 3 was used to find the clusters reported in Fig. 4A, B, and C named by their main functional-anatomical localizations: the detection, arousal, and salience network (DAS), task-positive network (TPN), and default mode network (DMN). (**B**, **E**, **H**, **K**) Silhouette values (a measure of within-cluster agreement) are shown for each cluster for (**B**) k = 2, (**E**) k = 3, (**H**) k = 4, and (**K**) k = 5. Mean voxel silhouette value for each cluster is marked by an open circle. Number of significant voxels in each cluster, and percentage of total number of significant voxels are also indicated. (**C**, **F**, **I**, **L**) Anatomical localization of all voxels within each cluster for (**C**) k = 2, (**F**) k = 3, (**I**) k = 4, and (**L**) k = 5. Report healthy participants N = 99.

**Fig. S18.**
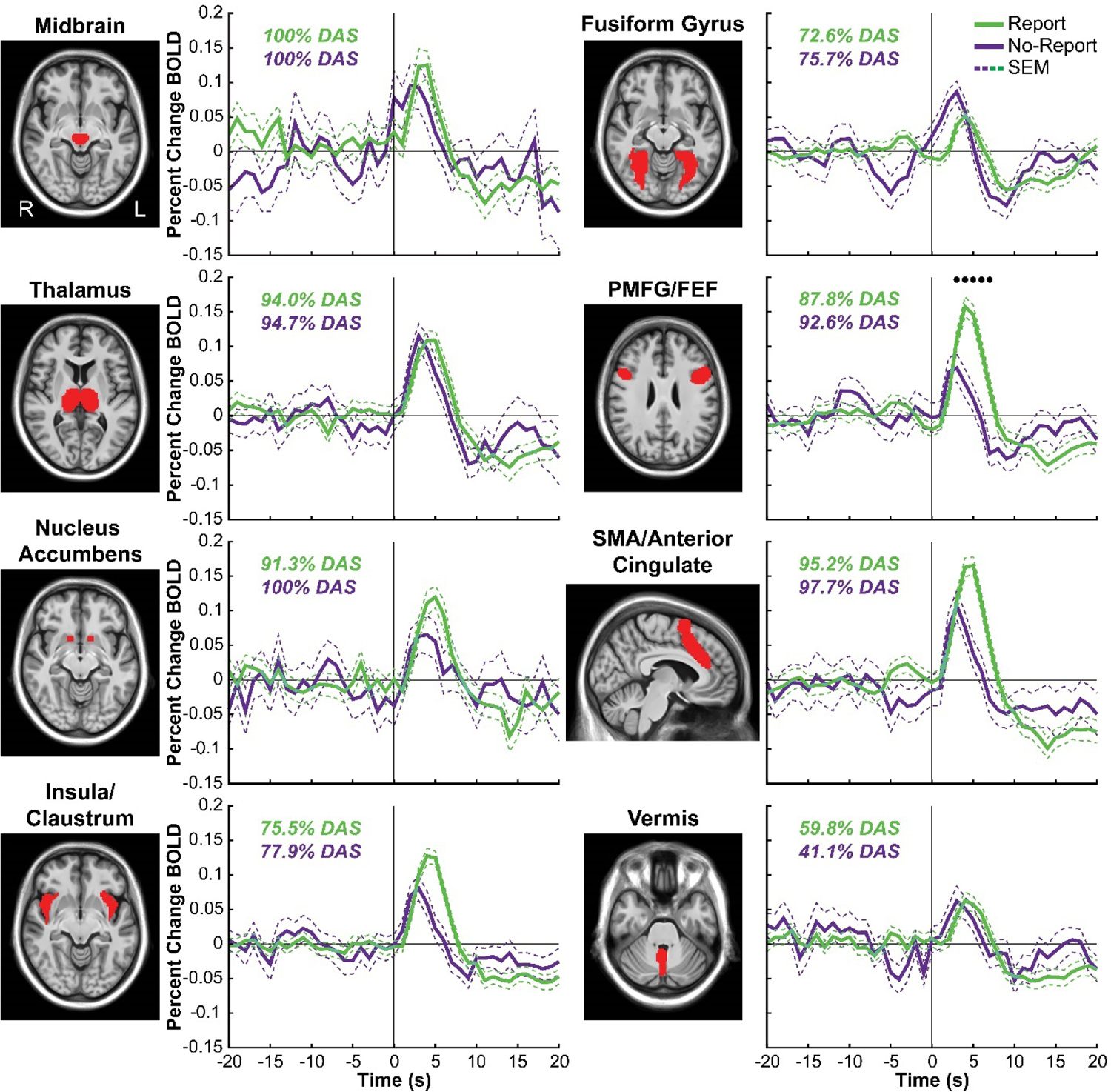
Report and no-report perceived minus not perceived percent change BOLD timecourses for DAS (detection, arousal, and salience) regions of interest. Report (green) and no-report (purple) mean percent change BOLD timecourses with standard error of the mean (SEM). The percentage of each region of interest (ROI) significant voxels overlapping with DAS voxels (see the *fMRI Anatomical Clustering* section) is shown for the report and no-report conditions. The selected ROIs were designated as belonging to DAS because more than 60% of overlapping ROI voxels from the report condition belonged to DAS. Stimulus onset was at time = 0. Times with significant differences between report and no-report data by permutation based testing (*p* < 0.05) are indicated by black dots. Posterior middle frontal gyrus (PMFG), frontal eye fields (FEF), and supplementary motor area (SMA). Data for report stimuli are from Report Paradigm (N = 34) and Report + No-Report Paradigm (N= 65); data for no-report stimuli are from Report + No-Report Paradigm (N = 65; one perceived subject data set was excluded because at the prediction score threshold of 0.75 there were no predicted perceived trials; see the *Covert Prediction of Conscious Perception – Machine Learning Pipeline* section).

**Fig. S19.**
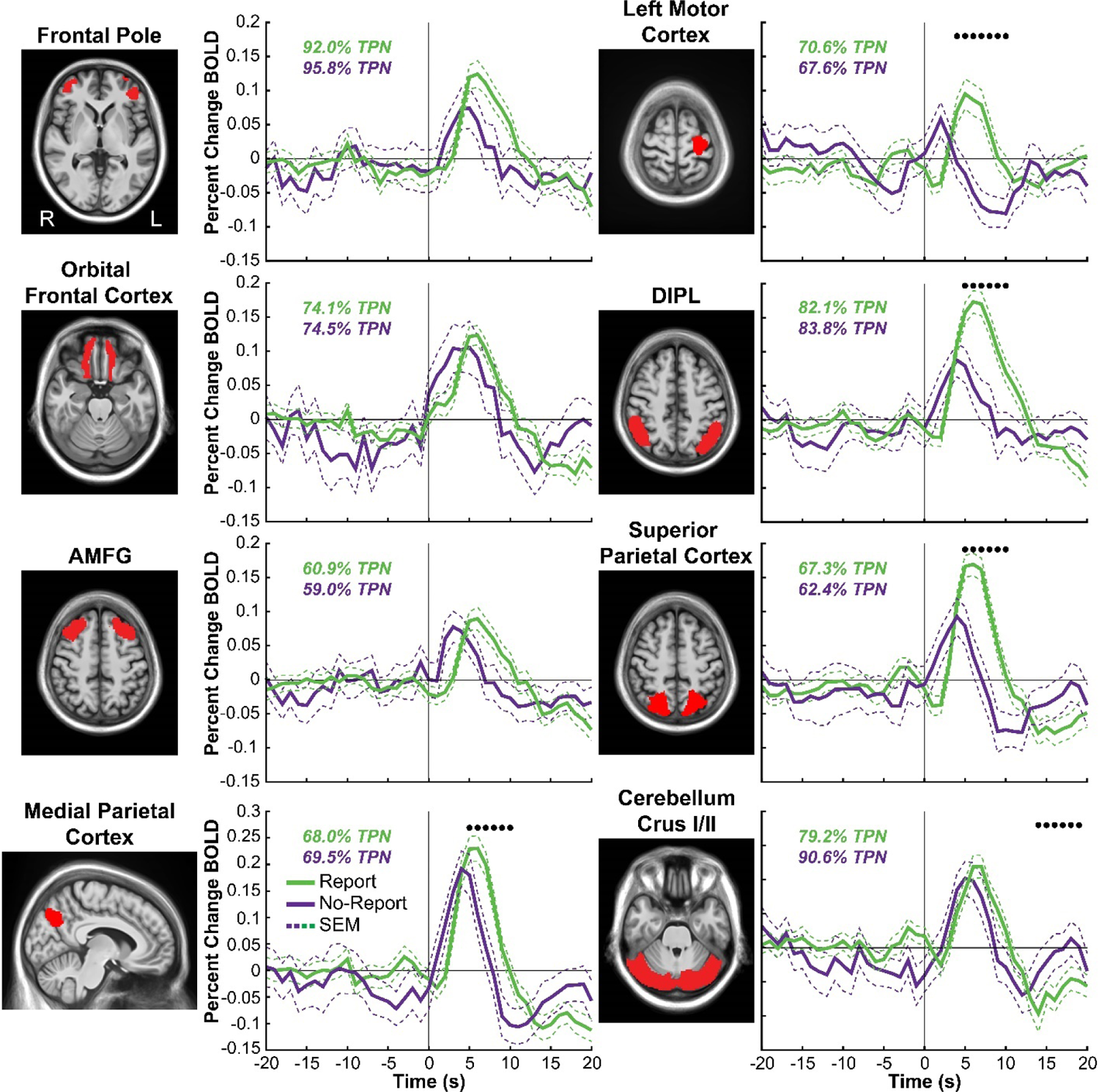
Report and no-report perceived minus not perceived percent change BOLD timecourses for TPN (task-positive network) regions of interest. Report (green) and no-report (purple) mean percent change BOLD timecourses with standard error of the mean (SEM). The percentage of each region of interest (ROI) significant voxels overlapping with TPN voxels (see the *fMRI Anatomical Clustering* section) is shown for the report and no-report conditions. The selected ROIs were designated as belonging to TPN because more than 60% of overlapping ROI voxels from the report condition belonged to TPN. Stimulus onset was at time = 0. Times with significant differences between report and no-report data by permutation based testing (*p* < 0.05) are indicated by black dots. Anterior middle frontal gyrus (AMFG), and dorsal inferior parietal lobule (DIPL). Data for report stimuli are from Report Paradigm (N = 34) and Report + No-Report Paradigm (N= 65); data for no-report stimuli are from Report + No-Report Paradigm (N = 65; one perceived subject data set was excluded because at the prediction score threshold of 0.75 there were no predicted perceived trials; see the *Covert Prediction of Conscious Perception – Machine Learning Pipeline* section).

**Fig. S20.**
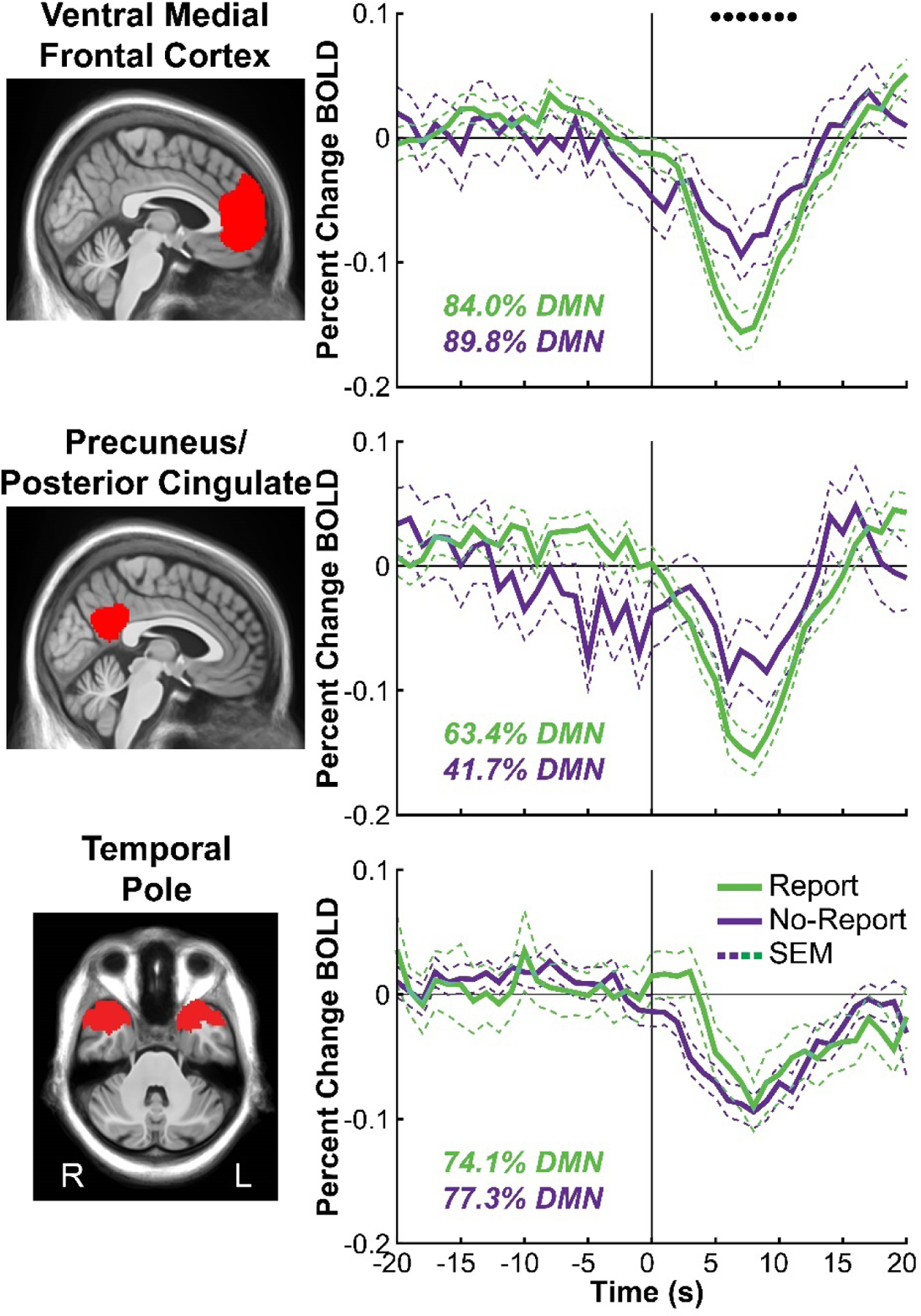
Report and no-report perceived minus not perceived percent change BOLD timecourses for DMN (default-mode network) regions of interest. Report (green) and no-report (purple) mean percent change BOLD timecourses with standard error of the mean (SEM). The percentage of each region of interest (ROI) significant voxels overlapping with DMN voxels (see the *fMRI Anatomical Clustering* section) is shown for the report and no-report conditions. The selected ROIs were designated as belonging to DMN because more than 60% of overlapping ROI voxels from the report condition belonged to DMN. Stimulus onset was at time = 0. Times with significant differences between report and no-report data by permutation based testing (*p* < 0.05) are indicated by black dots. Data for report stimuli are from Report Paradigm (N = 34) and Report + No-Report Paradigm (N= 65); data for no-report stimuli are from Report + No-Report Paradigm (N = 65; one perceived subject data set was excluded because at the prediction score threshold of 0.75 there were no predicted perceived trials; see the *Covert Prediction of Conscious Perception – Machine Learning Pipeline* section).

**Table S1.** All acquired healthy and patient participant data sets between the Report and Report + No-Report Paradigms and among the physiological recording types: fMRI, ldEEG, hdEEG, icEEG, and EyeLink. High-density EEG (hdEEG); low-density EEG (ldEEG); intracranial EEG (icEEG). *One icEEG patient participant did not have scalp ldEEG recordings.

| ***Behavioral Paradigm*** | ***Physiological Recordings*** | ***Participant #***<br>***(females)*** | ***Participant Age***<br>***Mean (range)*** | ***Right-Handed***<br>***N*** |
| --- | --- | --- | --- | --- |
| Report | hdEEG, EyeLink | 59 (37) | 26.20 (19-43) | 53 |
| Report + No-Report | hdEEG, EyeLink | 65 (39) | 24.58 (18-46) | 63 |
| Report | ldEEG, icEEG | 7* (5) | 24.86 (20-31) | 6 |
| Report | fMRI | 37 (17) | 27.22 (18-42) | 35 |
| Report + No-Report | fMRI, EyeLink | 65 (39) | 24.77 (18-46) | 64 |

**Table S2.** Patient participant icEEG recording system, recording parameters, channel information, and contact anatomical locations. icEEG recordings for participant 1 were made with Natus NeuroWorks Quantum (Natus, Inc.) and for participants 2-7 with the RNS System (Models RNS-300M and RNS-320; NeuroPace, Inc.). The depth contact location for participant 1 was determined using BioImage Suite and for participants 2-7 using Lead-DBS (see the *Depth Contact and Channel Localization* section). Left and right brain hemisphere locations are designated as “Left” and “Right.” Centromedian nucleus (CM), ventral posterior medial nucleus (VPM), ventral posterior lateral nucleus anterior part (VPLa), ventral posterior lateral nucleus posterior part (VPLp), ventral lateral posterior nucleus dorsal part (VLpd), ventral lateral posterior nucleus ventral part (VLpv), limitans nucleus (Li), posterior nucleus (Po), anterior pulvinar (PuA), neocortex (NC), and outside of thalamus (OT). *Contact 1 was defined as the most ventral and contact 4 as more dorsal. Participant 1 had only one depth lead with three contacts and two bipolar channels in the thalamus (channel 1 = contact 1-2; channel 2 = contact 2-3). For all other participants, there were four contacts and two bipolar channels per lead (channel 1 = contact 1-2; channel 2 = contact 3-4).

| **Participant** | **Recording System** | **Hardware Filters**<br>(high/low-pass) | **Lead Locations** | **Contact Anatomical Locations**<br>Contacts 1, 2, 3, 4* | **Number of Bipolar Channels** |
| --- | --- | --- | --- | --- | --- |
| 1 | Natus Quantum | 0.01/1757Hz | Right thalamus | *Right:* CL, VPLp, VPLp | *Right:* 2 channels |
| 2 | RNS-300M | 4/90Hz | Bilateral thalamus | *Left:* CM, PuA, VPLp, VPLp<br>*Right:* CM, VPLp, VLpd, VLpd | *Left:* 2 channels<br>*Right:* 2 channels |
| 3 | RNS-320 | 1/90Hz | Bilateral thalamus | *Left:* CM, VPM, VPLp, VLpv<br>*Right:* OT, CM, VPM, VLpv | *Left:* 2 channels<br>*Right:* 2 channels |
| 4 | RNS-320 | 5/70Hz | Bilateral thalamus | *Left:* Po, CM, VPM, VPLa<br>*Right:* CM, CM, VPM, VLpv | *Left:* 2 channels<br>*Right:* 2 channels |
| 5 | RNS-320 | 5/70Hz | Bilateral thalamus | *Left:* OT, OT, Po, CM<br>*Right:* OT, OT, CM, CM | *Left:* 2 channels<br>*Right:* 2 channels |
| 6 | RNS-320 | 1/90Hz | Thalamus/<br>neocortex | *Left:* OT, Li, CM, VPM<br>*Right:* NC, NC, NC, NC | *Left:* 2 channels<br>*Right:* 2 channels |
| 7 | RNS-320 | 1/90Hz | Bilateral thalamus | *Left:* OT, CM, VPM, VPLp<br>*Right:* OT, VPM, CM, VLpv | *Left:* 2 channels<br>*Right:* 2 channels |

**Table S3.** Patient participant thalamic bipolar channel information. Channels are grouped by participant and ordered within participant along the ventral to dorsal, axial (y-axis) dimension. Channels that are above or below the 3µV voltage amplitude threshold are indicated (see the *Intracranial EEG (icEEG)* section). The first peak polarity for each above threshold bipolar channel is shown. Below threshold channel first peak polarity is not applicable (NA). The position of each contact can be visualized in Fig. 2D and Fig. S12. *MNI coordinates are given for the Euclidian midpoint of the 2 electrode contacts for each recording channel.

| **Participant**<br>**(hemisphere)** | **Electrode**<br>**Contacts** | **MNI**<br>**Coordinate**<br>***x y z*\*** | **Above or**<br>**Below 3μV**<br>**Threshold** | **1<sup>st</sup> Peak**<br>**Polarity** |
| --- | --- | --- | --- | --- |
| 1 (*Right*) | Contacts 1 - 2 | 12 -16 8 | above | Negative |
| 1 (*Right*) | Contacts 2 - 3 | 15 -19 12 | above | Positive |
| 2 (*Left*) | Contacts 1 - 2 | -14 -22 2 | below | NA |
| 2 (*Right*) | Contacts 1 - 2 | 14 -19 5 | below | NA |
| 2 (*Left*) | Contacts 3 - 4 | -17 -19 10 | below | NA |
| 2 (*Right*) | Contacts 3 - 4 | 16 -14 12 | below | NA |
| 3 (*Right*) | Contacts 1 - 2 | 13 -23 -2 | above | Negative |
| 3 (*Left*) | Contacts 1 - 2 | -14 -21 1 | above | Positive |
| 3 (*Right*) | Contacts 3 - 4 | 16 -17 3 | above | Positive |
| 3 (*Left*) | Contacts 3 - 4 | -17 -17 8 | below | NA |
| 4(*Right*) | Contacts 1 - 2 | 13 -22 0 | above | Negative |
| 4 (*Left*) | Contacts 1 - 2 | -13 -23 1 | above | Negative |
| 4 (*Left*) | Contacts 3 - 4 | -15 -18 5 | below | NA |
| 4 (*Right*) | Contacts 3 - 4 | 14 -17 6 | below | NA |
| 5 (*Left*) | Contacts 1 - 2 | -10 -27 -7 | below | NA |
| 5 (*Right*) | Contacts 1 - 2 | 9 -26 -5 | below | NA |
| 5 (*Left*) | Contacts 3 - 4 | -13 -23 0 | above | Negative |
| 5 (*Right*) | Contacts 3 - 4 | 12 -21 1 | above | Negative |
| 6 (*Left*) | Contacts 1 - 2 | -12 -26 -4 | above | Negative |
| 6 (*Left*) | Contacts 3 - 4 | -14 -20 2 | above | Positive |
| 7(*Right*) | Contacts 1 - 2 | 12 -23 -5 | above | Negative |
| 7 (*Left*) | Contacts 1 - 2 | -11 -22 -2 | above | Positive |
| 7 (*Right*) | Contacts 3 - 4 | 14 -17 2 | below | NA |
| 7 (*Left*) | Contacts 3 - 4 | -15 -18 4 | below | NA |

**Table S4.** Selected tsfresh features from measurements of pupillary diameter for the covert prediction of conscious perception. Symbol represents each study session condition and is marked next to the tsfresh feature category that was selected for training the classification model (see *Covert Prediction of Conscious Perception – Machine Learning Pipeline* section). Each feature category can include multiple features. There are a total of 78 feature categories and 784 individual features across all categories. Documentation on each of the listed tsfresh features is available at https://tsfresh.readthedocs.io/en/latest/text/list_of_features.html

|  | *fMRI Center*<sup>^</sup> | *fMRI Quadrant*<sup>*</sup> | *hdEEG Center*<sup>o</sup> | *hdEEG Quadrant*<sup>#</sup> |
| --- | --- | --- | --- | --- |
| *Selected Tsfresh Features N* | 335 | 305 | 348 | 326 |
| *Feature categories* | *abs_energy*<sup>^*o#</sup>; *agg_autocorrelation*<sup>*</sup>; *absolute_sum_of_changes*<sup>^o#</sup>; *agg_linear_trend*<sup>^*o#</sup>; *approximate_entropy*<sup>^#</sup>; *ar_coefficient*<sup>^*#</sup>; *augmented_dickey_fuller*<sup>*o</sup>; *autocorrelation*<sup>*</sup>; *binned_entropy*<sup>^*</sup>; *c3*<sup>^*o#</sup>; *change_quantiles*<sup>^*o#</sup>; *count_above_mean*<sup>*</sup>; *cid_ce*<sup>^o#</sup>; *cwt_coefficients*<sup>^*o#</sup>; *energy_ratio_by_chunks*<sup>^*o#</sup>; *fft_aggregated*<sup>^*#</sup>; *fft_coefficient*<sup>^*o#</sup>; *first_location_maximum*<sup>^*o#</sup>; *first_location_minimum*<sup>o</sup>; *has_duplicate*<sup>^*</sup>; *has_duplicate_max*<sup>o</sup>; *index_mass_quantile*<sup>^*o</sup>; *large_standard_deviation*<sup>^*o#</sup>; *last_location_of_maximum*<sup>^*o#</sup>; *last_location_of_minimum*<sup>o</sup>; *length*<sup>*</sup>; *linear_trend*<sup>^*o#</sup>; *longest_strike_above_mean*<sup>^*o#</sup>; *longest_strike_below_mean*<sup>^#</sup>; *maximum*<sup>*o</sup>; *mean*<sup>^o#</sup>; *mean_abs_change*<sup>^o#</sup>; *mean_change*<sup>o</sup>; *mean_second_derivative_central*<sup>o</sup>; *median*<sup>*o#</sup>; *minimum*<sup>^*o#</sup>; *number_crossing*<sup>^</sup>; *number_cwt_peaks*<sup>^*#</sup>; *number_peaks*<sup>^*o#</sup>; *partial_autocorrelation*<sup>*</sup>; *percentage_of_reoccurring_datapoints_to_all_datapoints*<sup>^*o</sup>; *percentage_of_reoccurring_values_to_all_values*<sup>^*o</sup>; *quantile*<sup>^*o#</sup>; *range_count_max*<sup>*</sup>; *ratio_beyond_r_sigma*<sup>^o#</sup>; *ratio_value_number_to_time_series_length*<sup>^</sup>; *sample_entropy*<sup>^*#</sup>; *spkt_welch_density*<sup>^*</sup>; *standard_deviation*<sup>^*o#</sup>; *sum_of_reoccurring_data_points*<sup>^*o#</sup>; *sum_of_reoccurring_values*<sup>^*o#</sup>; *sum_values*<sup>^*o#</sup>; *symmetry_looking*<sup>^*o#</sup>; *time_reversal_asymmetry_statistic*<sup>o</sup>; *variance*<sup>^*o#</sup>; *variance_larger_than_standard_deviation*<sup>*</sup> |  |  |  |

**Table S5.**
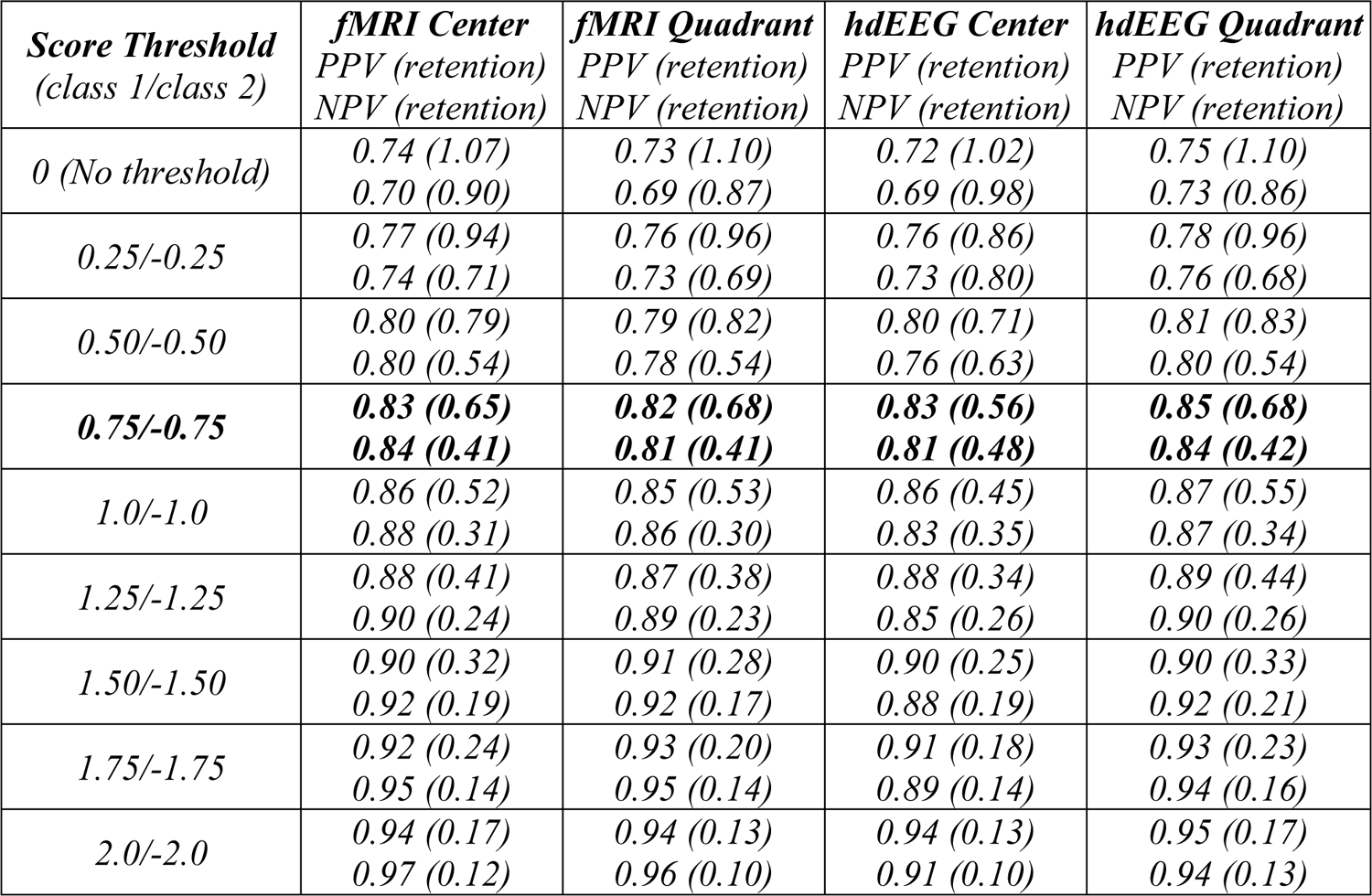
Classification performance on the report (task-relevant) condition from the Report + No-Report Paradigm. Classification performance measured by the positive predictive values (PPV), negative predictive values (NPV), and trial retention relative to number of trials for report perceived and not perceived data, respectively, across score thresholds (0 to -/+2.0 in increments of 0.25) and the four behavioral task-relevant and neuroimaging study session conditions (fMRI Center, fMRI Quadrant, hdEEG Center, and hdEEG Quadrant location set task-relevant). The number of report perceived and not perceived trials among the study session conditions: fMRI center (2472 perceived; 1868 not perceived), fMRI quadrant (2375 perceived; 1791 not perceived), hdEEG center (2341 perceived; 2057 not perceived), and hdEEG quadrant (2576 perceived; 1904 not perceived). **Bold** values represent the score thresholds implemented for main figures (Fig. 2-4). These thresholds were selected because they compromised between high PPV and NPV and maintaining high retention. Note that to maximize the number of trials for classifier adjustment, the trials used and totals reported are prior to trial, task block, and participant-level rejections implemented later for analysis and visualization of EyeLink, fMRI, and hdEEG data. The values shown in this table are the same as those found in Fig. S4. High-density EEG (hdEEG).

**Table S6.** Classification performance on the no-report (task-irrelevant) condition from the Report + No-Report Paradigm. Classification performance measured by the number of no-report predicted perceived (PP) and predicted not perceived (PNP) trials across score thresholds (0 to -/+2.0 in increments of 0.25) and the four behavioral task-irrelevant and neuroimaging study session conditions (fMRI Center, fMRI Quadrant, hdEEG Center, and hdEEG Quadrant location set task-irrelevant). The total number of task-irrelevant trials (N) are shown only for the perceptual threshold opacity stimuli, as opaque and blank opacity stimuli were not included in the classification testing set. The retention (ret.) for each class is computed as the predicted perceived and not perceived trials divided by the total number of task-irrelevant trials within each study session. As in Table S5, these values are reported prior to trial, task block, and participant-level rejections implemented later for analysis and visualization of the EyeLink, fMRI, and hdEEG data. **Bold** values represent the score thresholds implemented for main figures (Fig. 2-4). The values shown in this table are the same as those found in Fig. S5. High-density EEG (hdEEG).

| **Score Threshold**<br>(class 1/class 2) | **fMRI Center**<br>PP trials, N (ret.)<br>PNP trials, N (ret.) | **fMRI Quadrant**<br>PP trial, N (ret.)<br>PNP trial, N (ret.) | **hdEEG Center**<br>PP trial, N (ret.)<br>PNP trial, N (ret.) | **hdEEG Quadrant**<br>PP trial, N (ret.)<br>PNP trial, N (ret.) |
| --- | --- | --- | --- | --- |
| *0 (No threshold)* | 2394 (0.48)<br>2592 (0.52) | 2433 (0.47)<br>2715 (0.53) | 2083 (0.39)<br>3299 (0.61) | 2567 (0.48)<br>2760 (0.52) |
| *0.25/-0.25* | 1873 (0.38)<br>2051 (0.41) | 1887 (0.37)<br>2211 (0.43) | 1564 (0.29)<br>2724 (0.51) | 2004 (0.38)<br>2219 (0.42) |
| *0.50/-0.50* | 1445 (0.29)<br>1559 (0.31) | 1395 (0.27)<br>1729 (0.34) | 1158 (0.22)<br>2110 (0.39) | 1523 (0.29)<br>1773 (0.33) |
| **0.75/-0.75** | **1062 (0.21)**<br>**1140 (0.23)** | **1035 (0.20)**<br>**1297 (0.25)** | **791 (0.15)**<br>**1604 (0.30)** | **1131 (0.21)**<br>**1376 (0.26)** |
| *1.0/-1.0* | 747 (0.15)<br>829 (0.17) | 705 (0.14)<br>956 (0.19) | 535 (0.10)<br>1185 (0.22) | 817 (0.15)<br>1026 (0.19) |
| *1.25/-1.25* | 506 (0.10)<br>630 (0.13) | 463 (0.09)<br>724 (0.14) | 352 (0.07)<br>845 (0.16) | 551 (0.10)<br>798 (0.15) |
| *1.50/-1.50* | 298 (0.06)<br>477 (0.10) | 276 (0.05)<br>475 (0.09) | 227 (0.04)<br>604 (0.11) | 346 (0.07)<br>628 (0.12) |
| *1.75/-1.75* | 178 (0.04)<br>359 (0.07) | 139 (0.03)<br>372 (0.07) | 143 (0.03)<br>424 (0.08) | 225 (0.04)<br>498 (0.09) |
| *2.0/-2.0* | 104 (0.02)<br>277 (0.06) | 77 (0.02)<br>289 (0.06) | 81 (0.02)<br>295 (0.05) | 141 (0.03)<br>395 (0.07) |
| **Total Trials (N)** | 4986 | 5148 | 5382 | 5327 |

**Slide S1. Whole brain fMRI maps for perceived minus not perceived stimuli with overt report.** Statistically significant voxels are shown from cluster-based permutation tests (*p* < 0.05) for 20 seconds pre and post-stimulus presentation. Statistically significant positive and negative voxel cluster *t*-values are shown in warm and cool colors, respectively. These images arise from the same data and analyses displayed in Fig. 3A and B where only the 3 and 6 seconds post-stimulus presentation timepoints are shown. Data are from the Report Paradigm (N = 34) and Report + No-Report Paradigm (N = 65).

**Slide S2. Whole brain fMRI maps for perceived minus not perceived stimuli without overt report.** Statistically significant voxels are shown from cluster-based permutation tests (*p* < 0.05) for 20 seconds pre and post-stimulus presentation. Statistically significant positive and negative voxel cluster *t*-values are shown in warm and cool colors, respectively. These images arise from the same data and analyses displayed in Fig. 3C and D where only the 3 and 6 seconds post-stimulus presentation timepoints are shown. Data are from the Report + No-Report Paradigm (N = 65; one perceived subject data set was excluded because at the prediction score threshold of 0.75 there were no predicted perceived trials; see the *Covert Prediction of Conscious Perception – Machine Learning Pipeline* section).

**Slide S3. Whole brain fMRI maps for report versus no-report perceived minus not perceived.** Statistically significant voxels are shown from cluster-based permutation tests (*p* < 0.05) for 20 seconds pre and post-stimulus presentation. Statistically significant positive and negative voxel cluster *t*-values are shown in warm and cool colors, respectively. These images arise from the same data and analyses displayed in fig. S16A and B where only the 3 and 6 seconds post-stimulus presentation timepoints are shown. Data for report stimuli are from the Report Paradigm (N = 34) and Report + No-Report Paradigm (N = 65); data for no-report stimuli are from the Report + No-Report Paradigm (N = 65; one perceived subject data set was excluded because at the prediction score threshold of 0.75 there were no predicted perceived trials; see the *Covert Prediction of Conscious Perception – Machine Learning Pipeline* section).

**Slide S4. Whole brain conjunction maps for report versus no-report perceived minus not perceived.** Voxel-level conjunction analysis of report and no-report, perceived minus not perceived statistical whole brain maps with shared (report-independent) increases shown in green and shared decreases shown in purple for 20 seconds pre and post-stimulus presentation. These images arise from the same data and analyses displayed in Fig. 3E and F where only the 3 and 6 seconds post-stimulus presentation timepoints are shown. Data for report stimuli are from the Report Paradigm (N = 34) and Report + No-Report Paradigm (N = 65); data for no-report stimuli are from the Report + No-Report Paradigm (N = 65; one perceived subject data set was excluded because at the prediction score threshold of 0.75 there were no predicted perceived trials; see the *Covert Prediction of Conscious Perception – Machine Learning Pipeline* section.

**Slide S5. Whole brain exclusive disjunction maps showing perceived minus not perceived signals significant only for report.** Analysis of signals significant only in report across 20 seconds pre and post-stimulus presentation (see the *fMRI Spatiotemporal Conjunction and Exclusive Disjunction Analyses* section for full details on finding regions not shared between report and no-report conditions). Exclusive disjunction voxels are color coded by membership to either positive (yellow) or negative (blue) statistically significant clusters determined by cluster-based permutation tests (*p* < 0.05). These images arise from the same data and analyses displayed in fig. S16C and D where only the 3 and 6 seconds post-stimulus presentation timepoints are shown. Data for report stimuli are from the Report Paradigm (N = 34) and Report + No-Report Paradigm (N = 65); data for no-report stimuli are from the Report + No-Report Paradigm (N = 65; one perceived subject data set was excluded because at the prediction score threshold of 0.75 there were no predicted perceived trials; see the *Covert Prediction of Conscious Perception – Machine Learning Pipeline* section).

**Slide S6. Whole brain exclusive disjunction maps showing perceived minus not perceived signals significant only for no-report.** Analysis of signals significant only in no-report across 20 seconds pre and post-stimulus presentation (see the *fMRI Spatiotemporal Conjunction and Exclusive Disjunction Analyses* section for full details on finding regions not shared between report and no-report conditions). Exclusive disjunction voxels are color coded by membership to either positive (yellow) or negative (blue) statistically significant clusters determined by cluster-based permutation tests (*p* < 0.05). These images arise from the same data and analyses displayed in fig. S16E and F where only the 3 and 6 seconds post-stimulus presentation timepoints are shown. Data for report stimuli are from the Report Paradigm (N = 34) and Report + No-Report Paradigm (N = 65); data for no-report stimuli are from the Report + No-Report Paradigm (N = 65; one perceived subject data set was excluded because at the prediction score threshold of 0.75 there were no predicted perceived trials; see the *Covert Prediction of Conscious Perception – Machine Learning Pipeline* section).

