## Supplementary material for "Brain networks in human conscious visual perception": Slide S1

### Slide 1:

#### T-Value Maps of Statistically Significant Voxels from Cluster-Based Permutation Analysis

Report - Perceived minus Not Perceived

*20 seconds pre-stimulus and post-stimulus*

*Authors:*

*Kronemer et al., 2021*

### 20s pre-stimulus

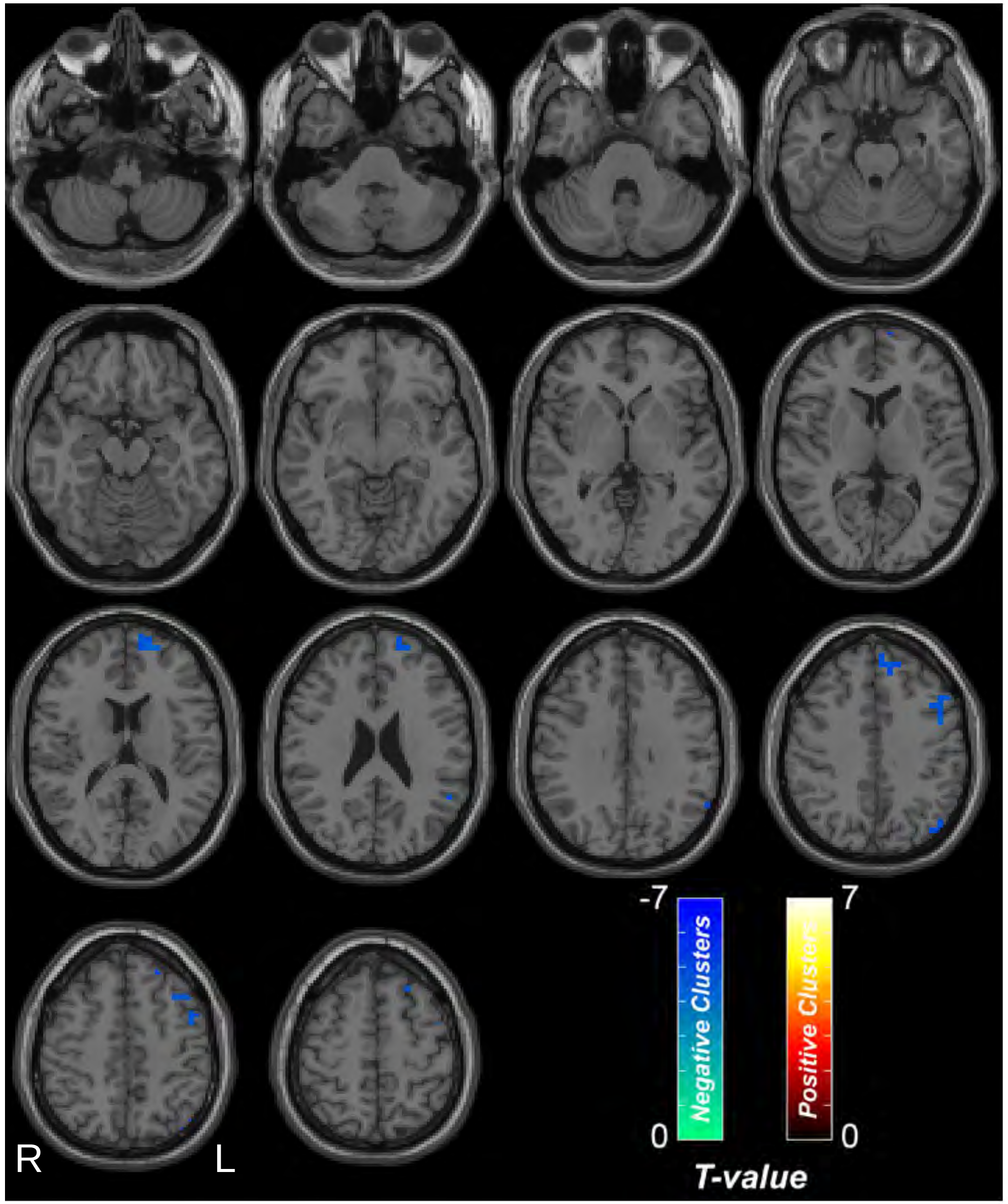


Kronemer et al., 2021

### 19s pre-stimulus

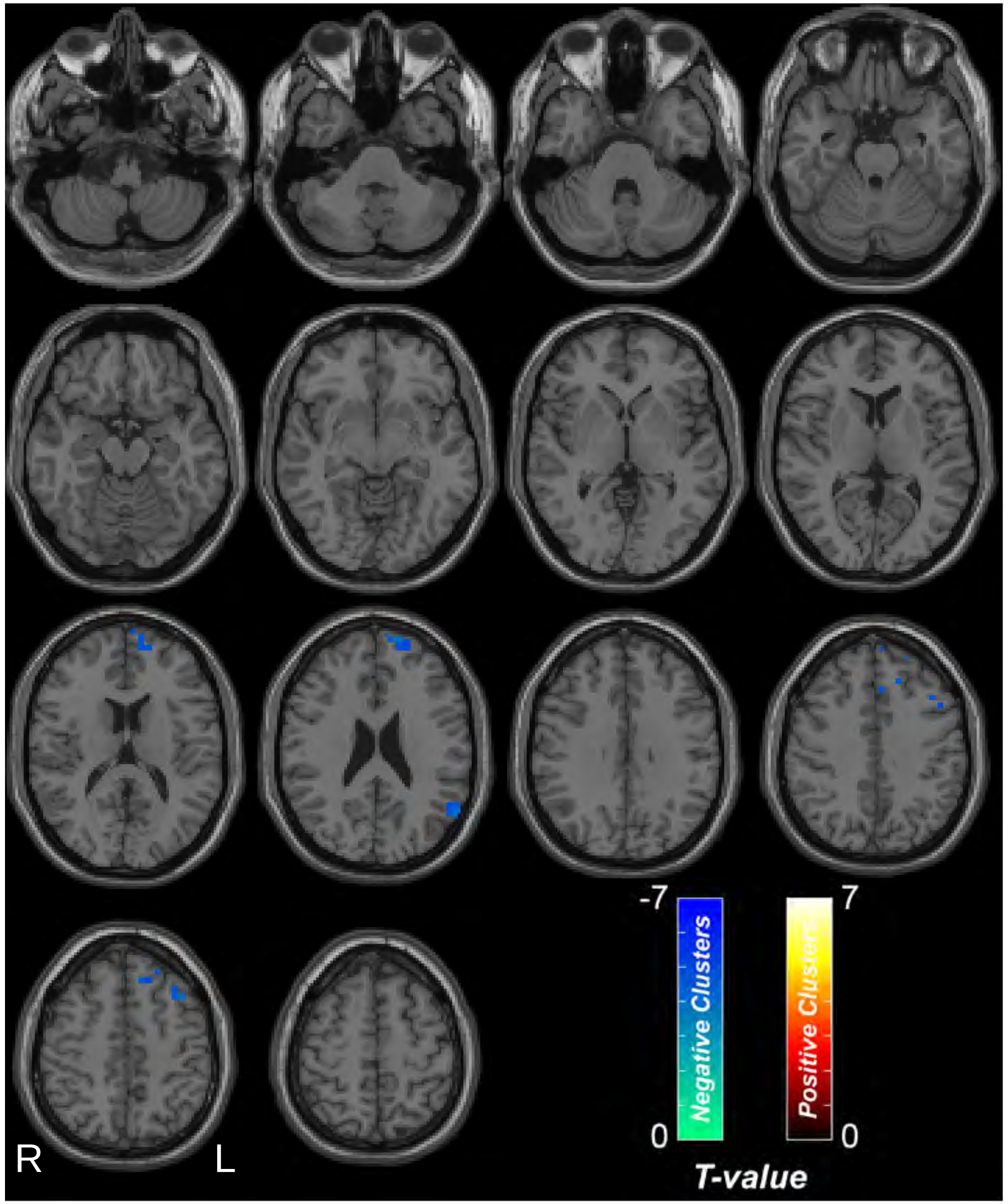


Kronemer et al., 2021

### 18s pre-stimulus

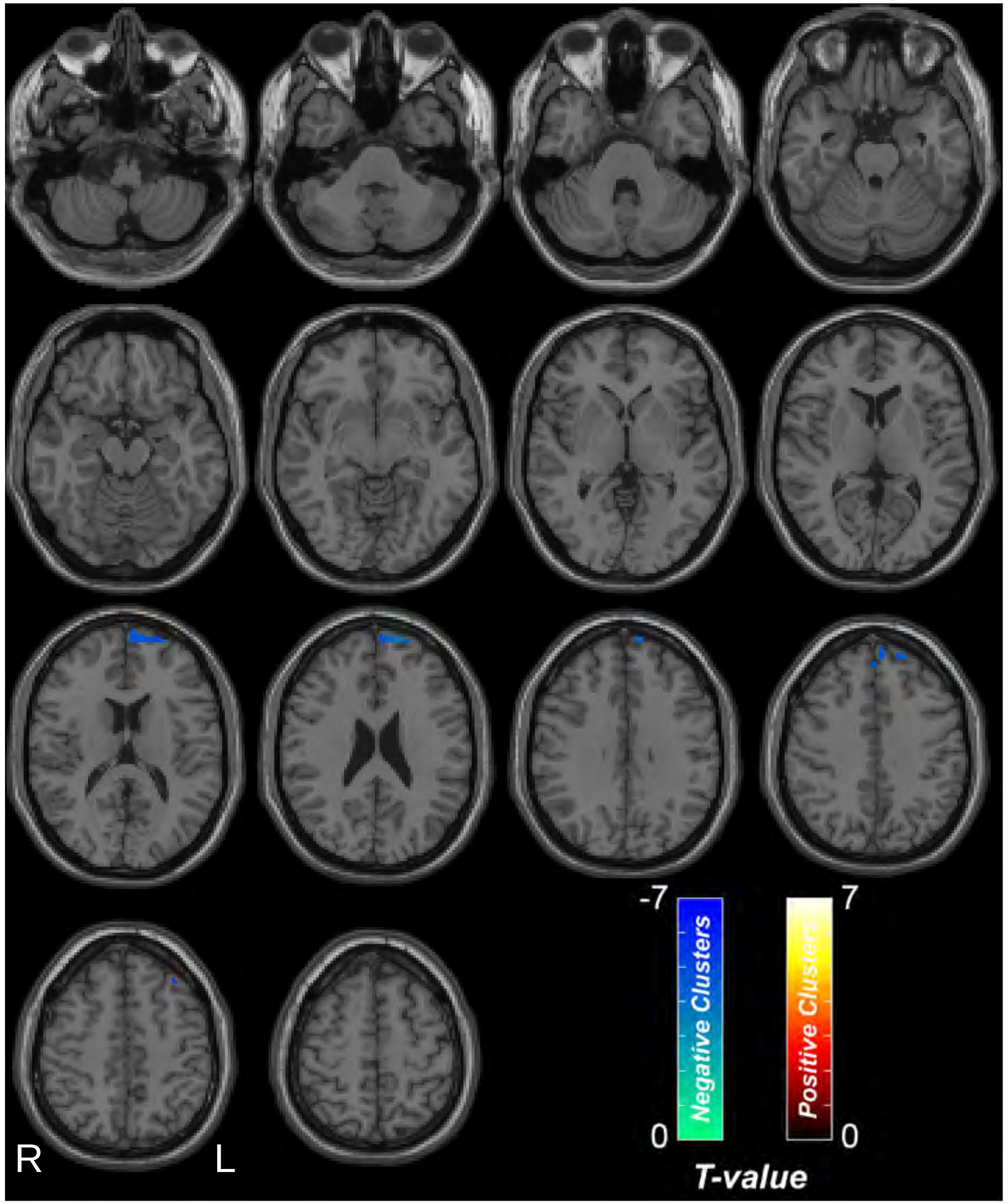


Kronemer et al., 2021

### 17s pre-stimulus

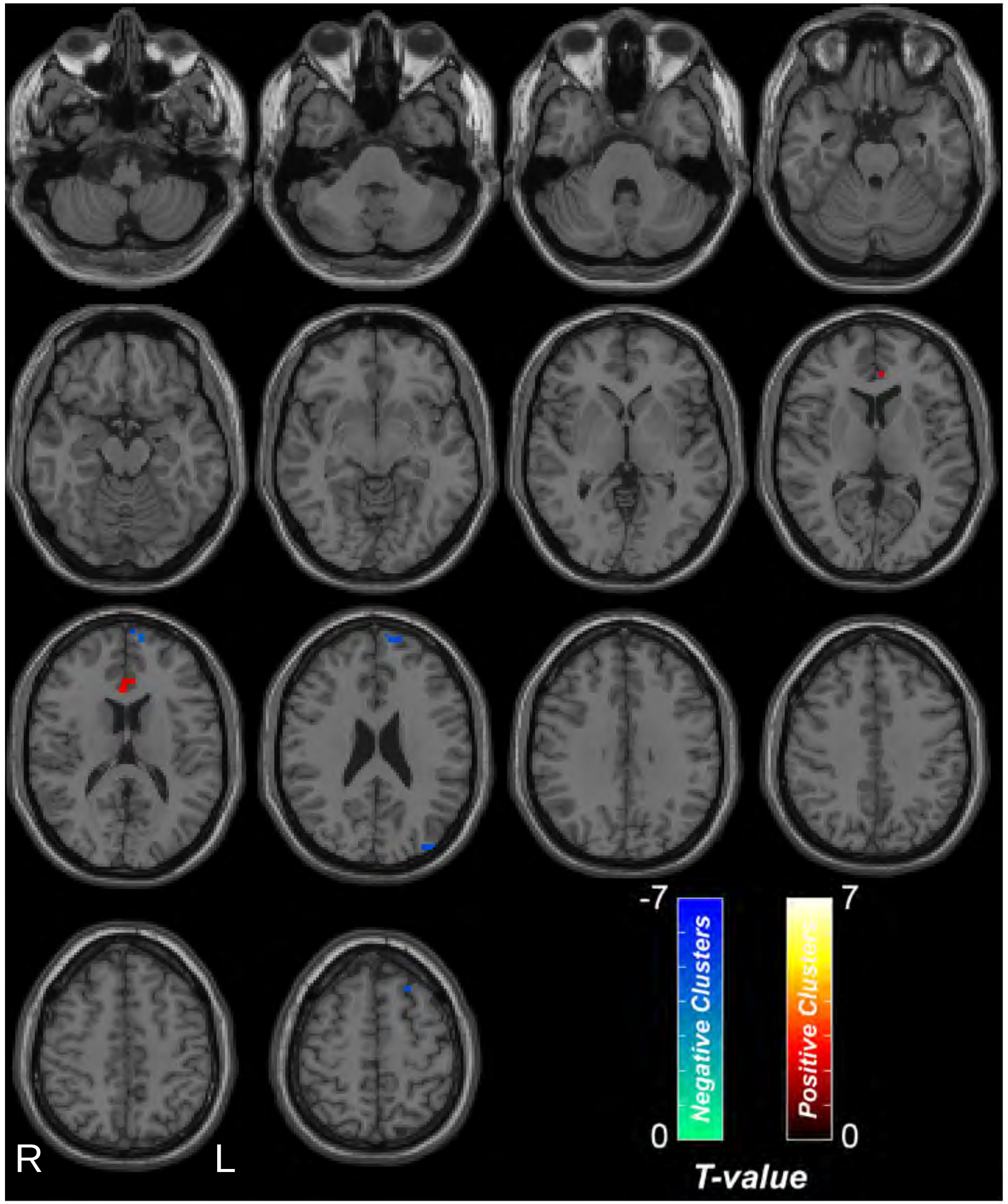


Kronemer et al., 2021

### 16s pre-stimulus

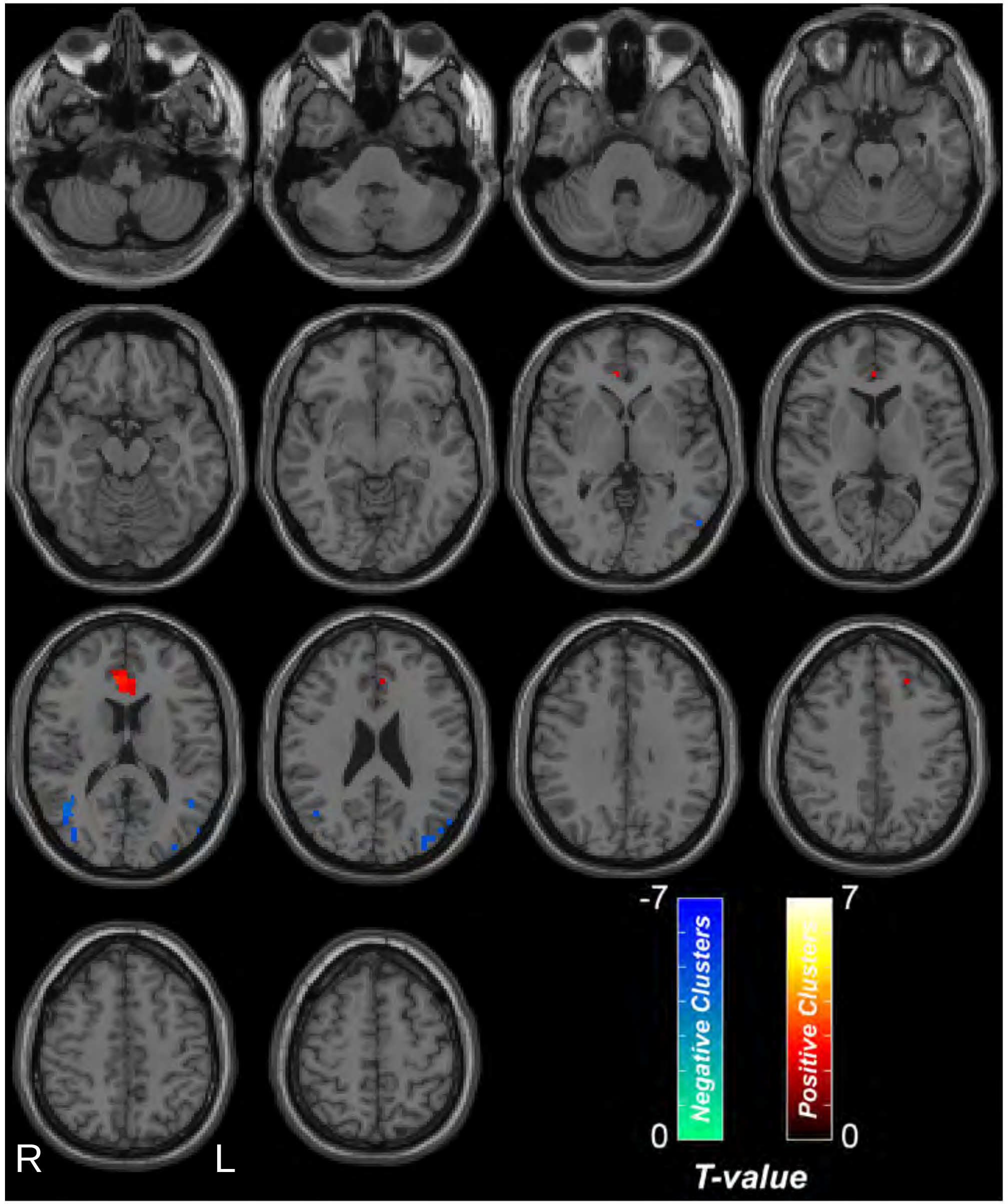


Kronemer et al., 2021

### 15s pre-stimulus

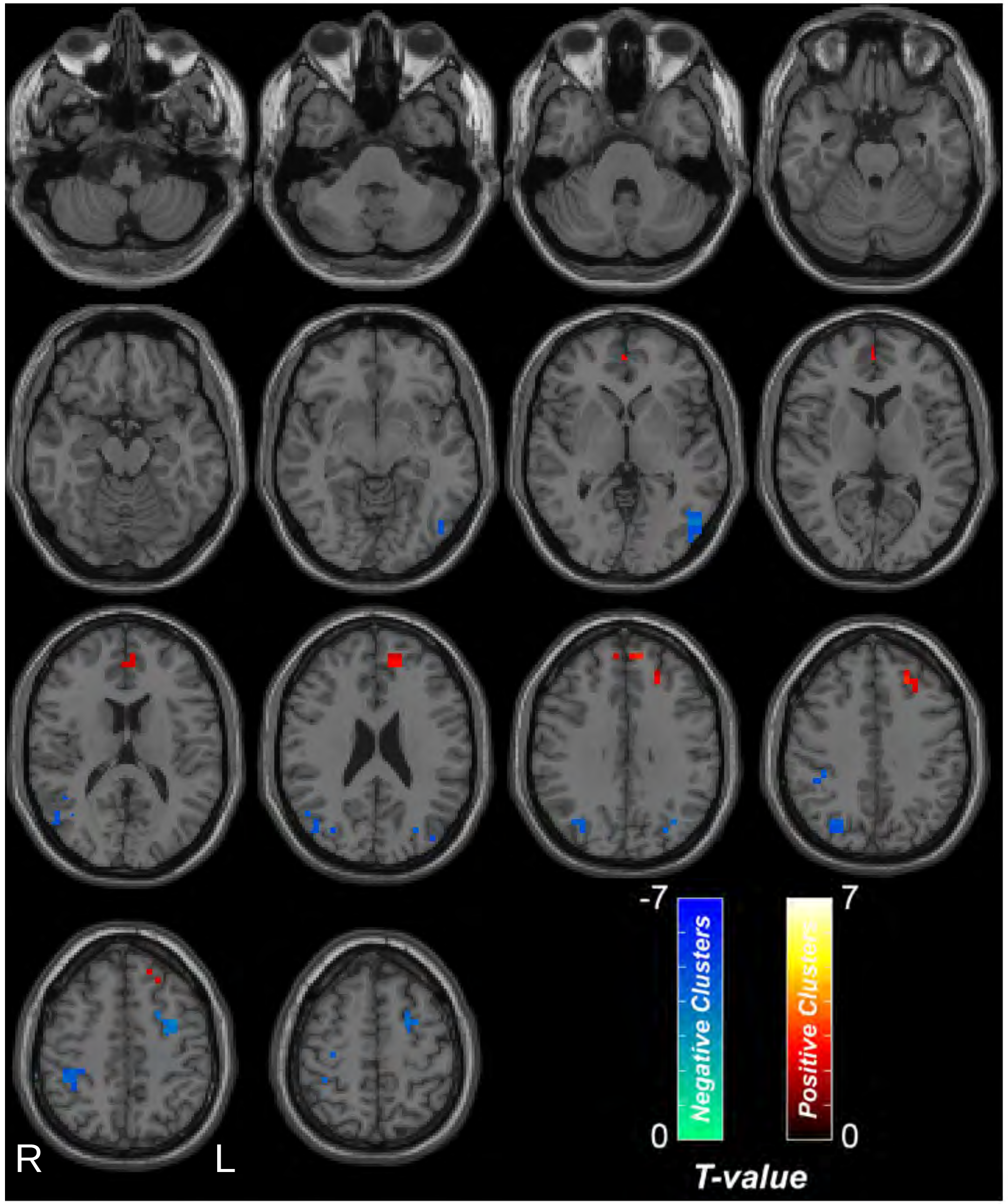


Kronemer et al., 2021

### 14s pre-stimulus

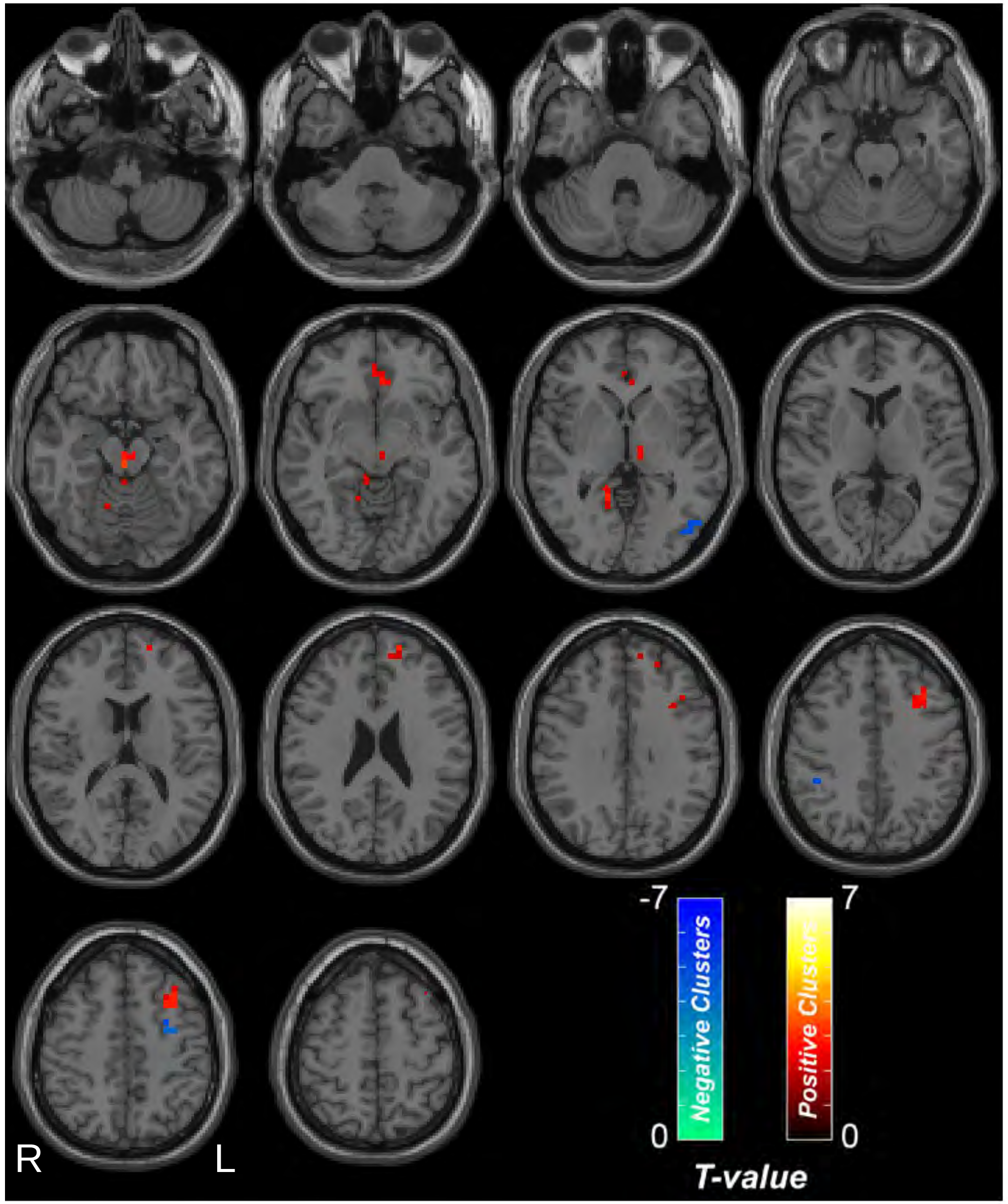


Kronemer et al., 2021

### 13s pre-stimulus

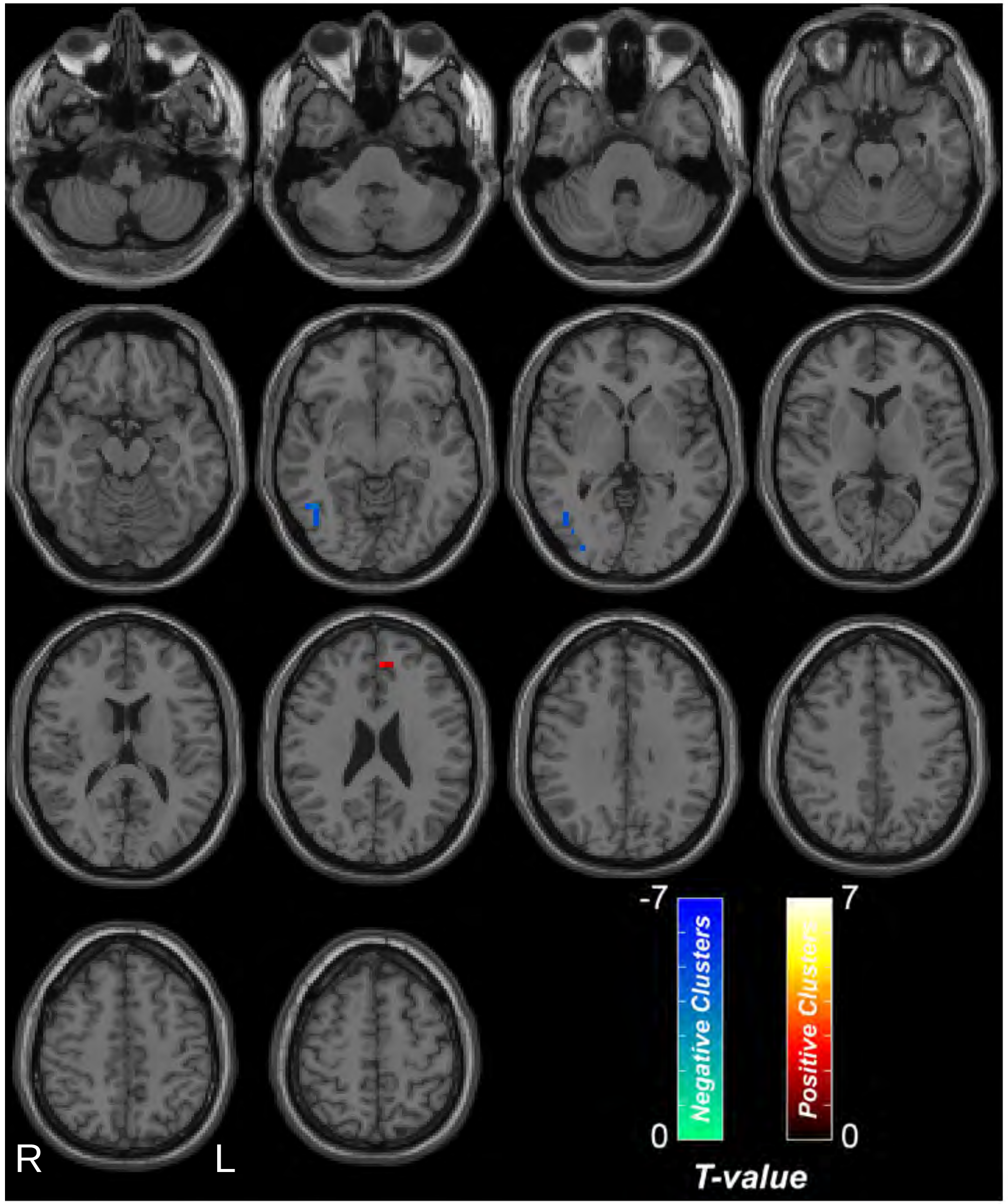


Kronemer et al., 2021

### 12s pre-stimulus

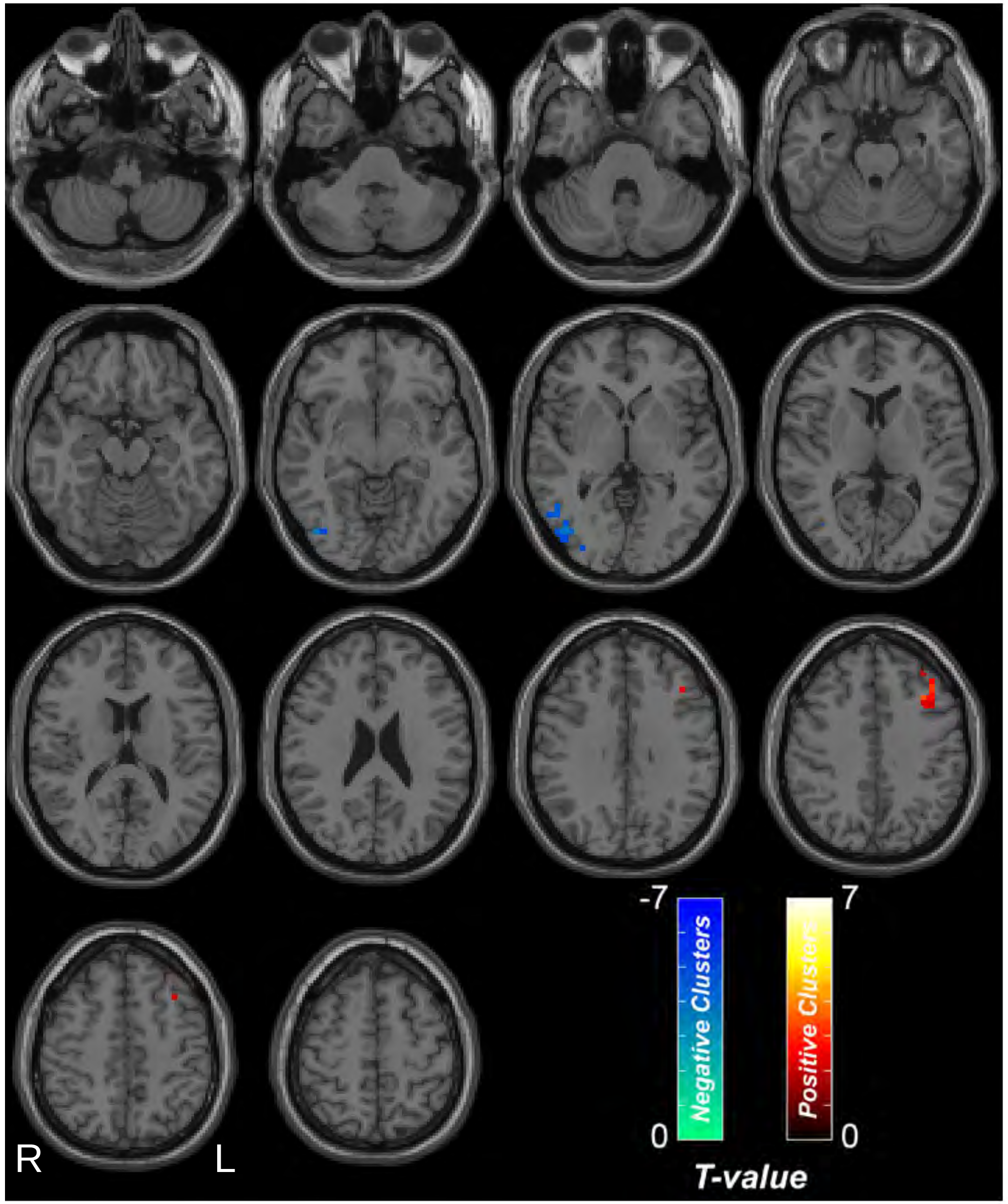


Kronemer et al., 2021

### 11s pre-stimulus

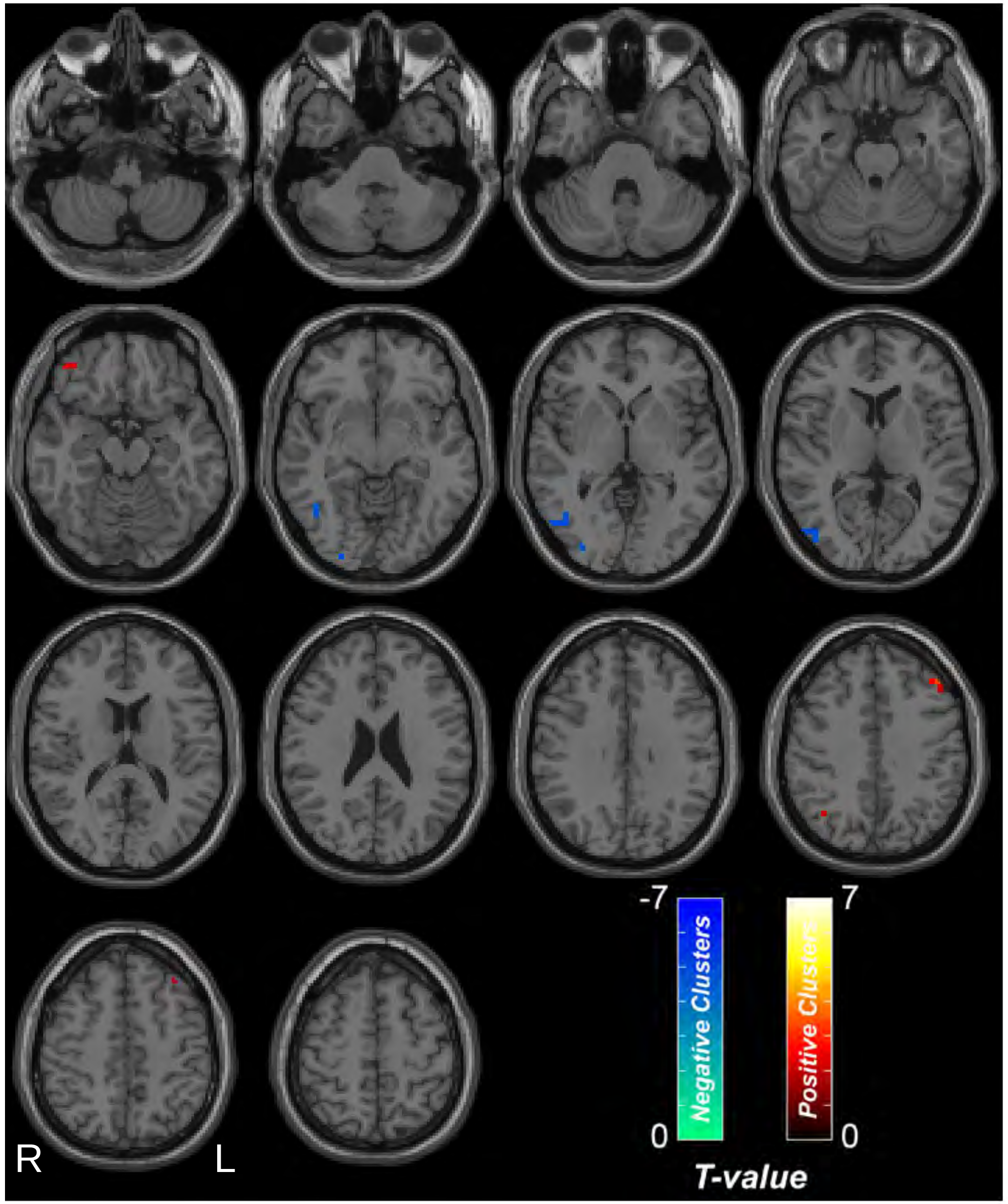


Kronemer et al., 2021

### 10s pre-stimulus

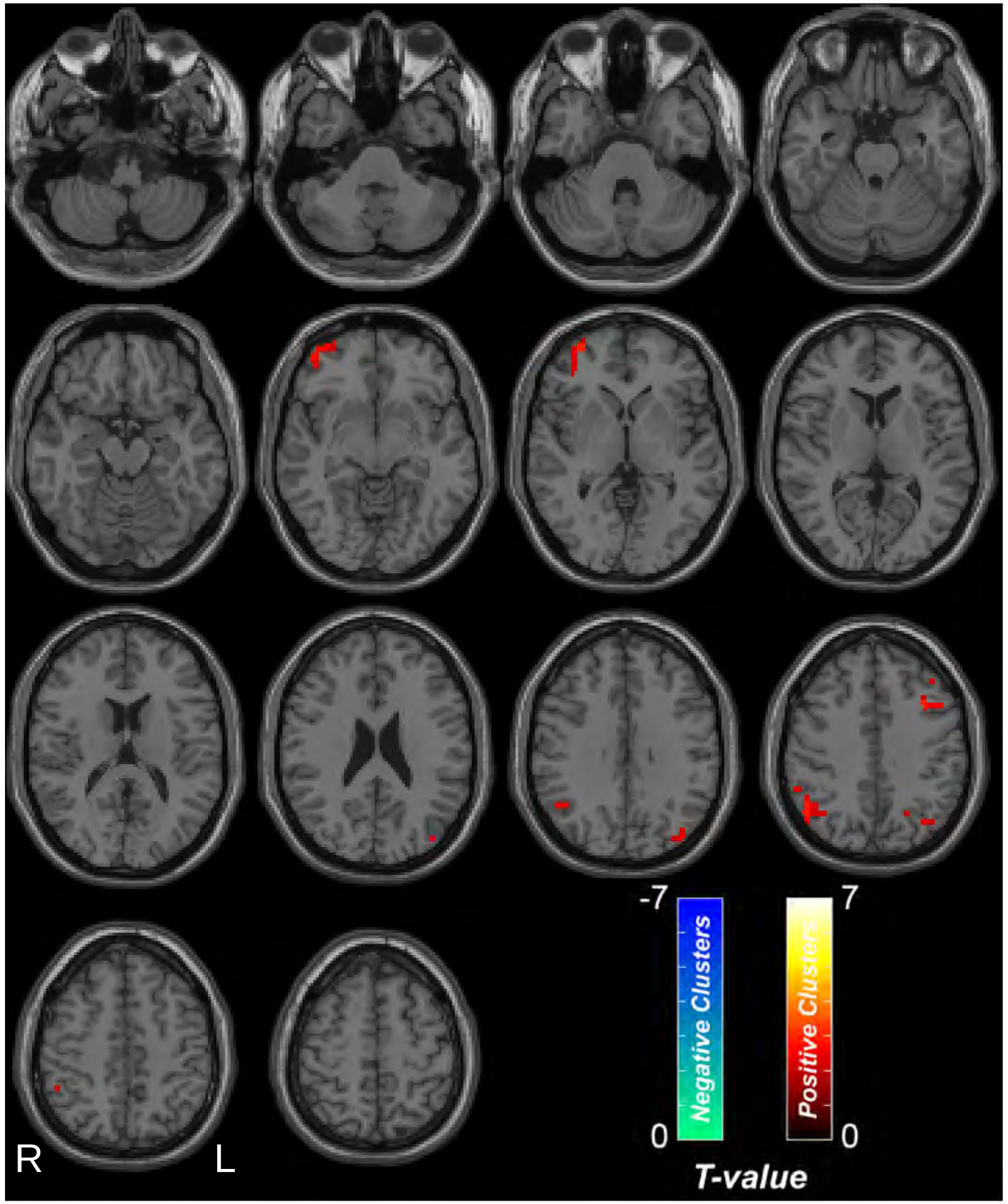


Kronemer et al., 2021

### 9s pre-stimulus

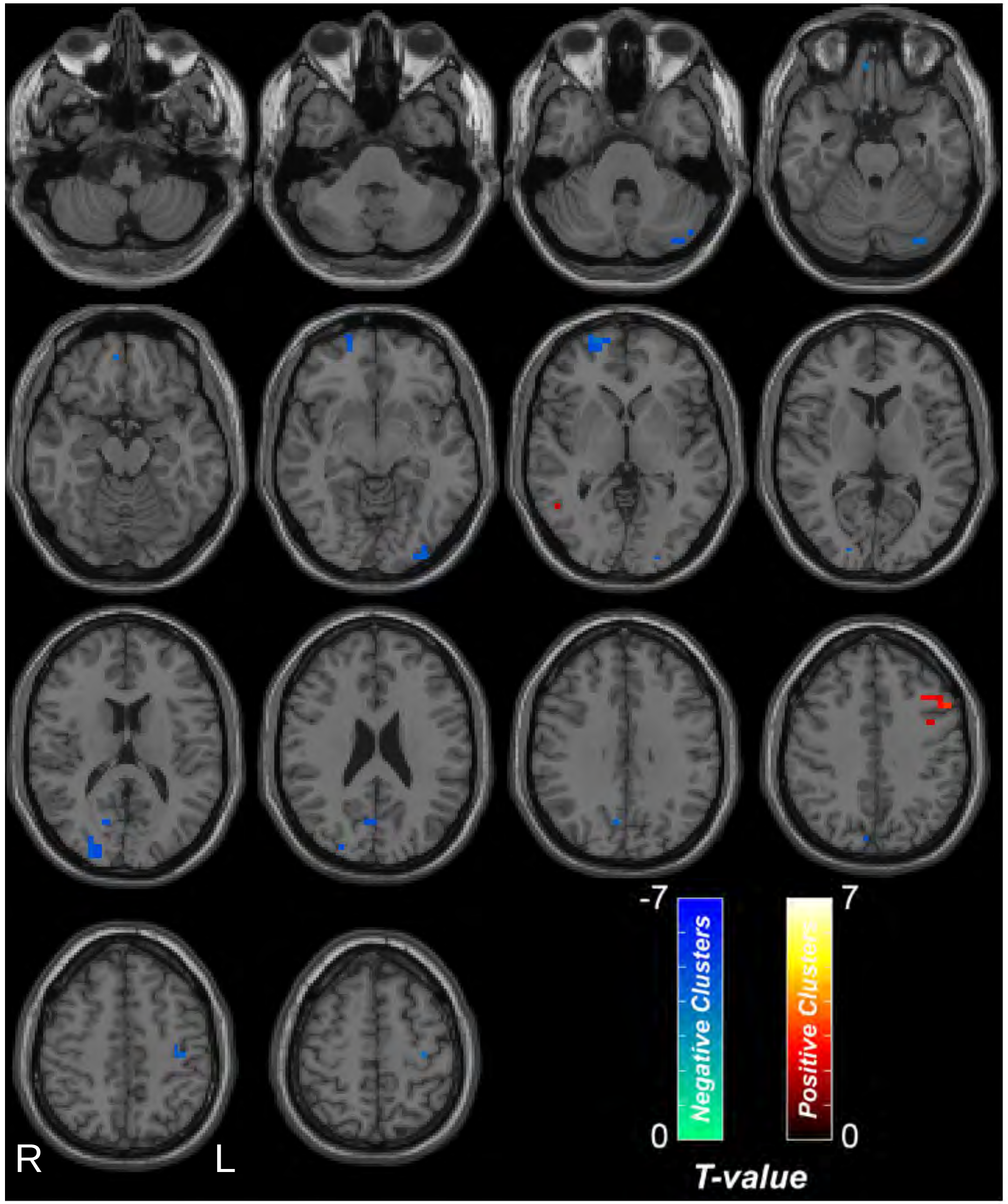


Kronemer et al., 2021

### 8s pre-stimulus

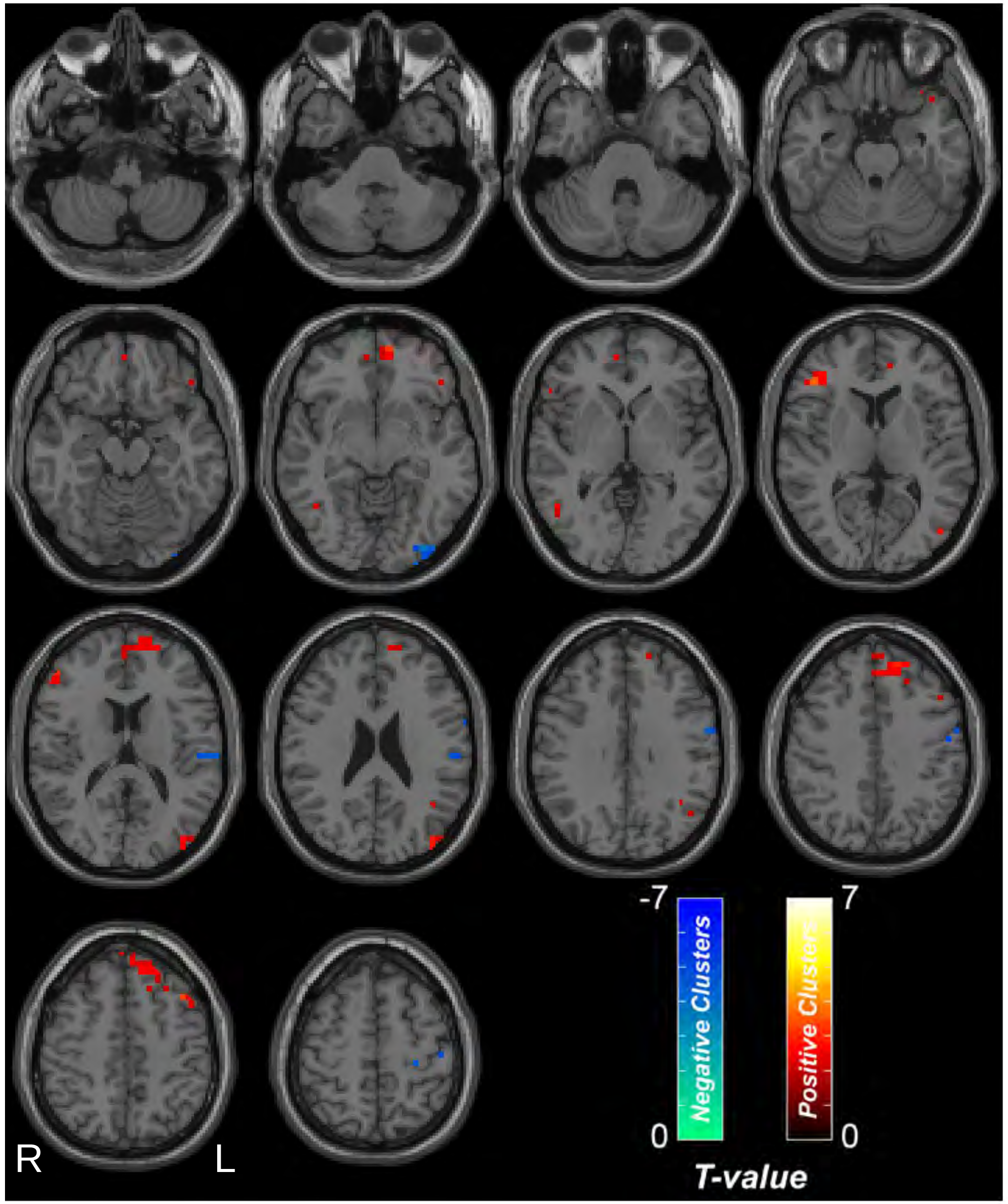


Kronemer et al., 2021

### 7s pre-stimulus

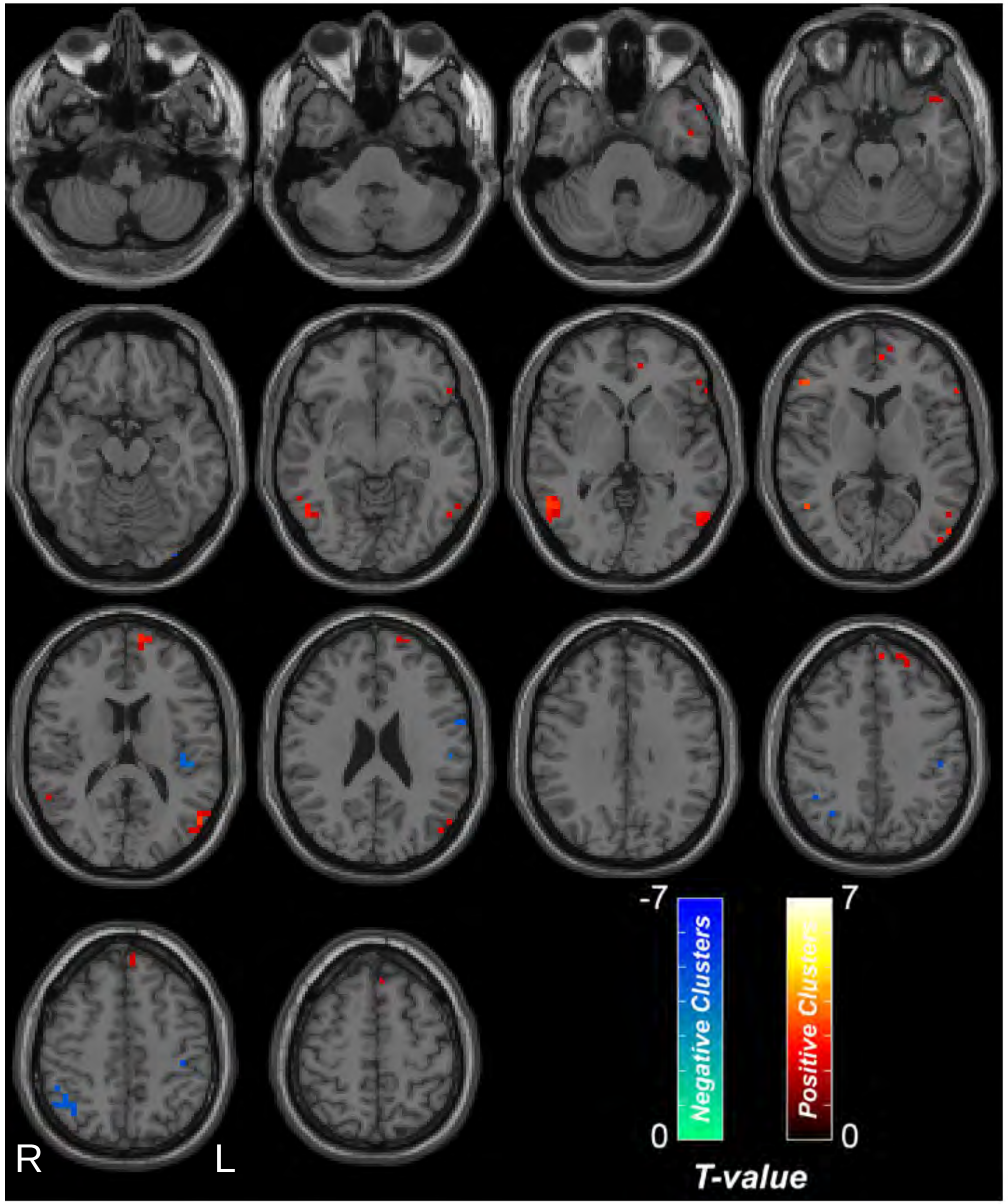


Kronemer et al., 2021

### 6s pre-stimulus

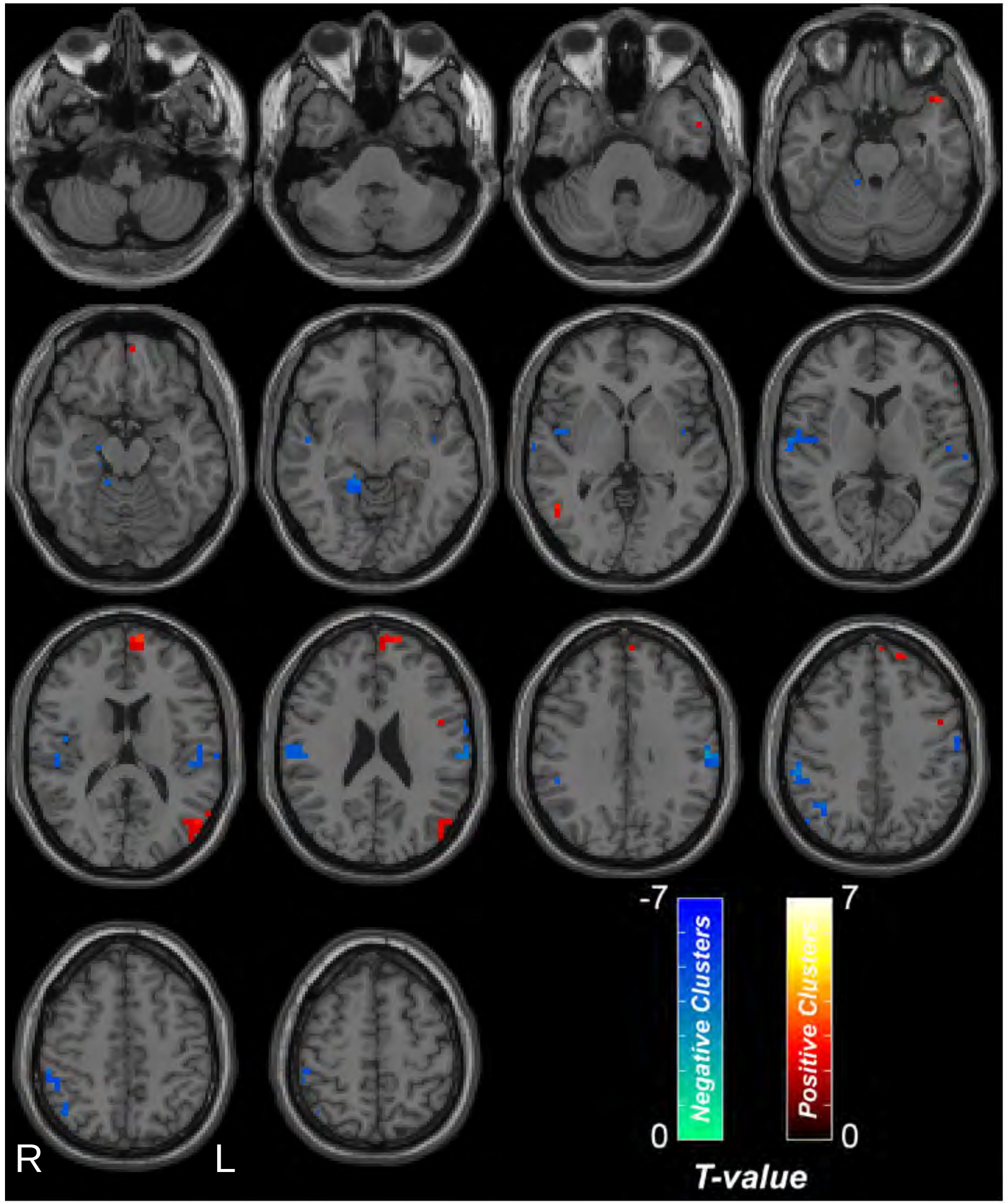


Kronemer et al., 2021

### 5s pre-stimulus

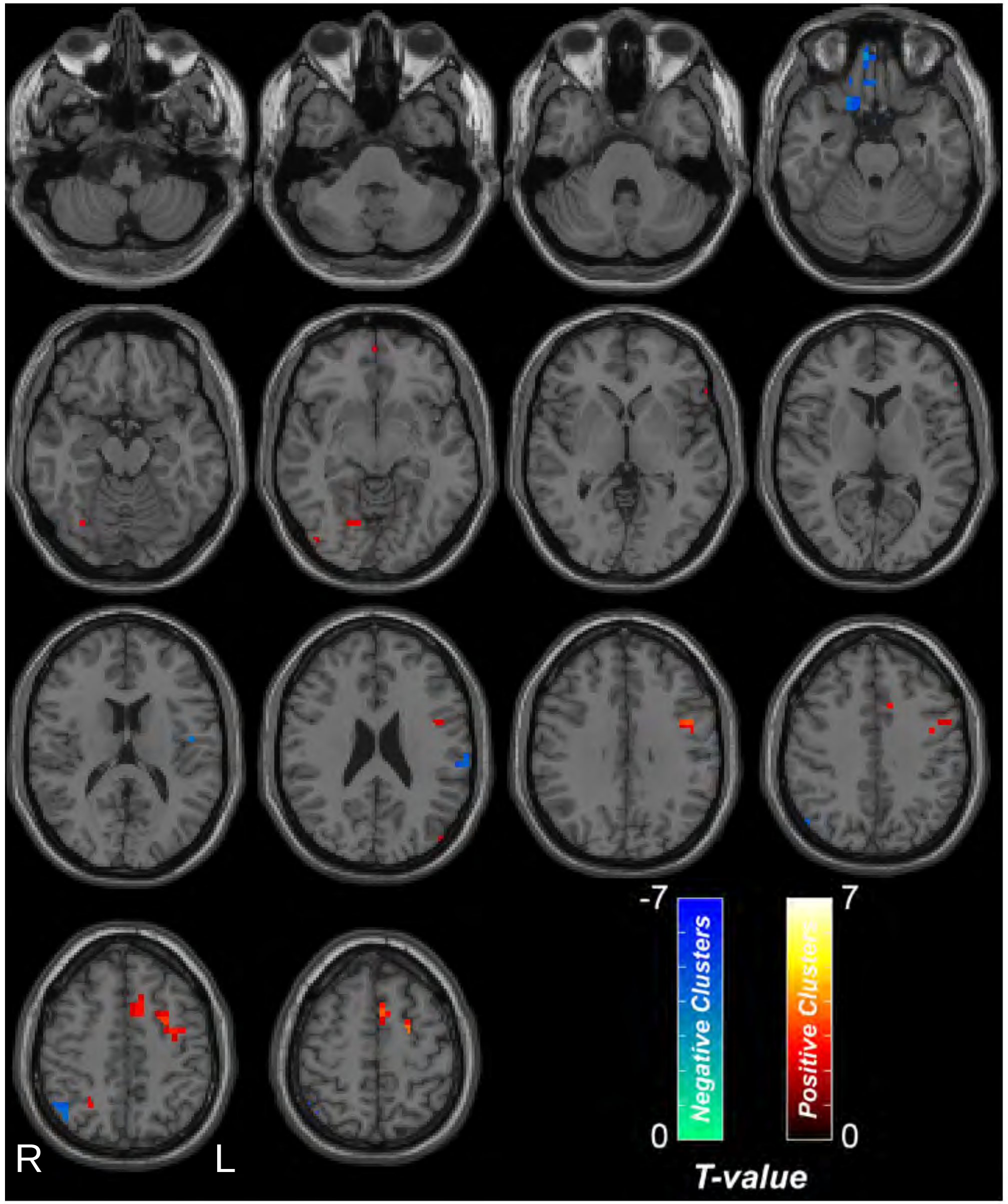


Kronemer et al., 2021

### 4s pre-stimulus

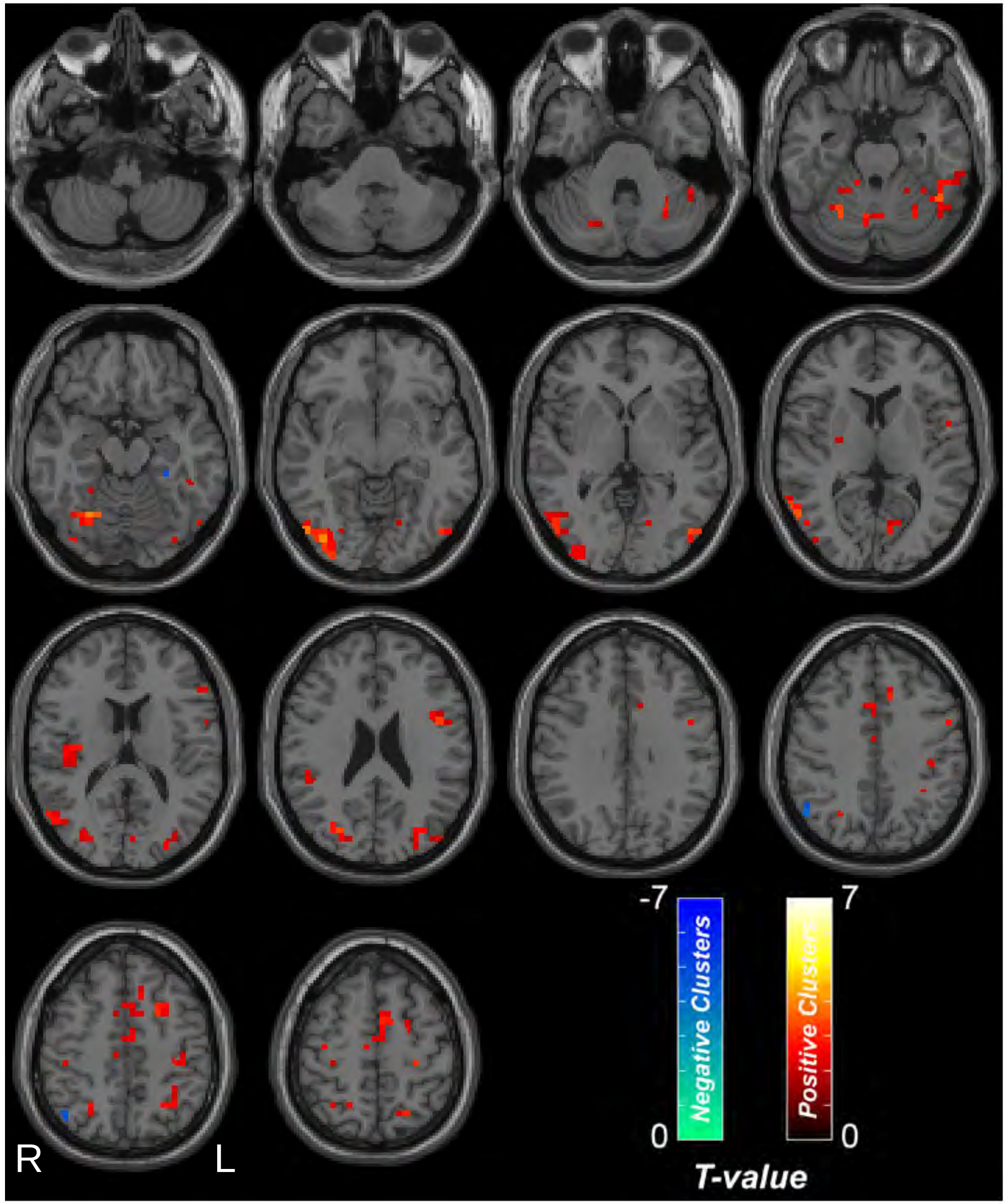


Kronemer et al., 2021

### 3s pre-stimulus

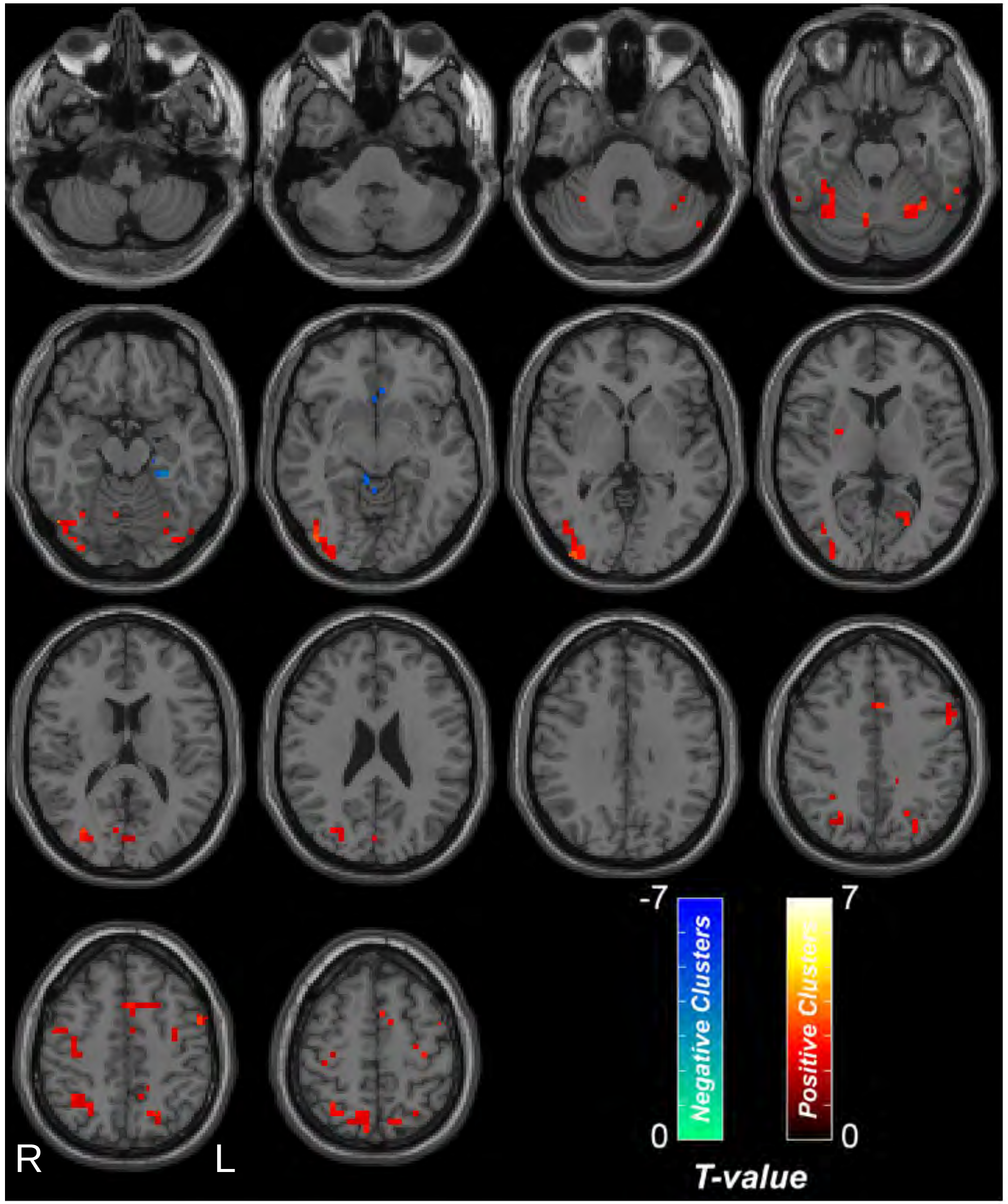


Kronemer et al., 2021

### 2s pre-stimulus

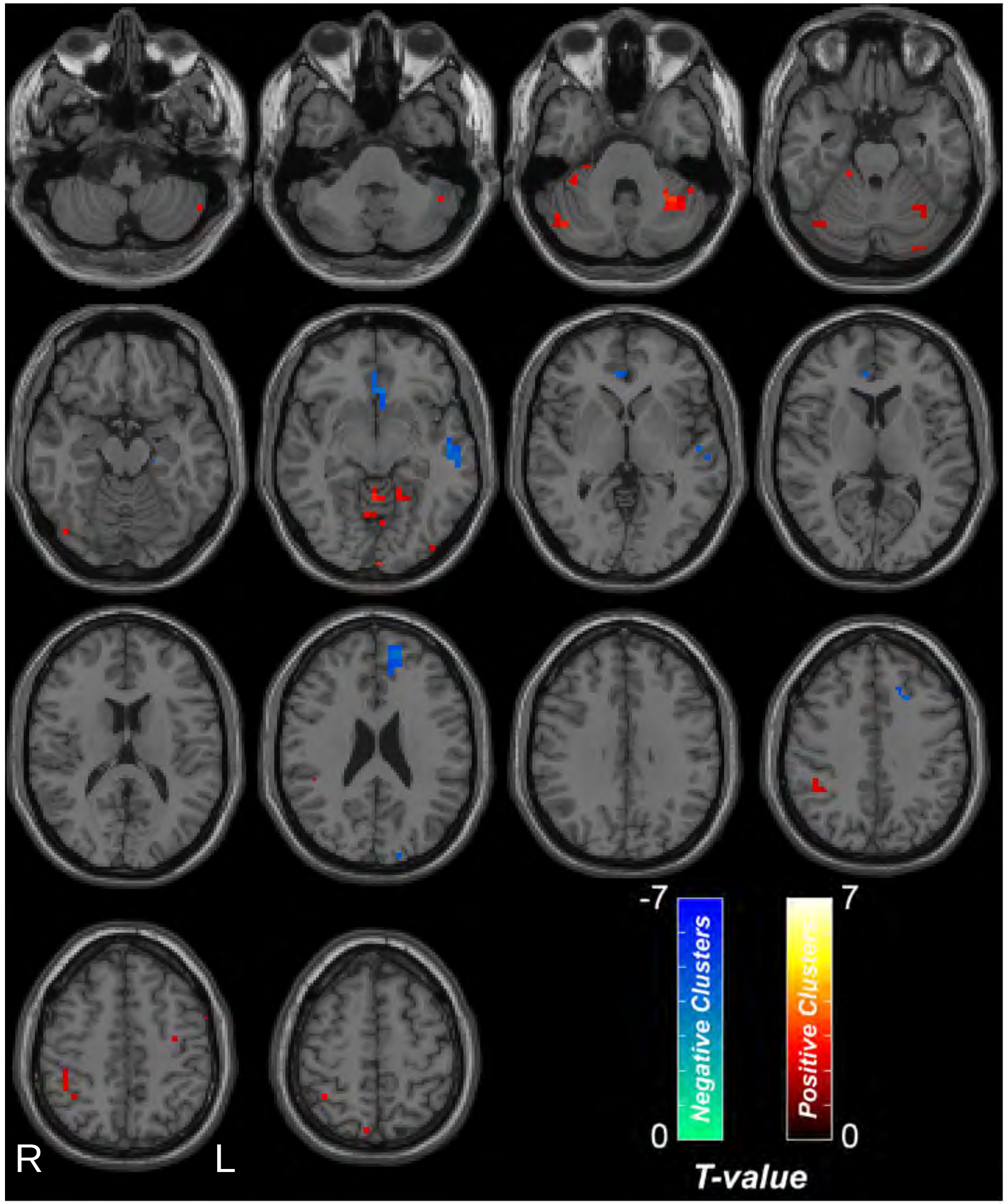


Kronemer et al., 2021

### 1s pre-stimulus

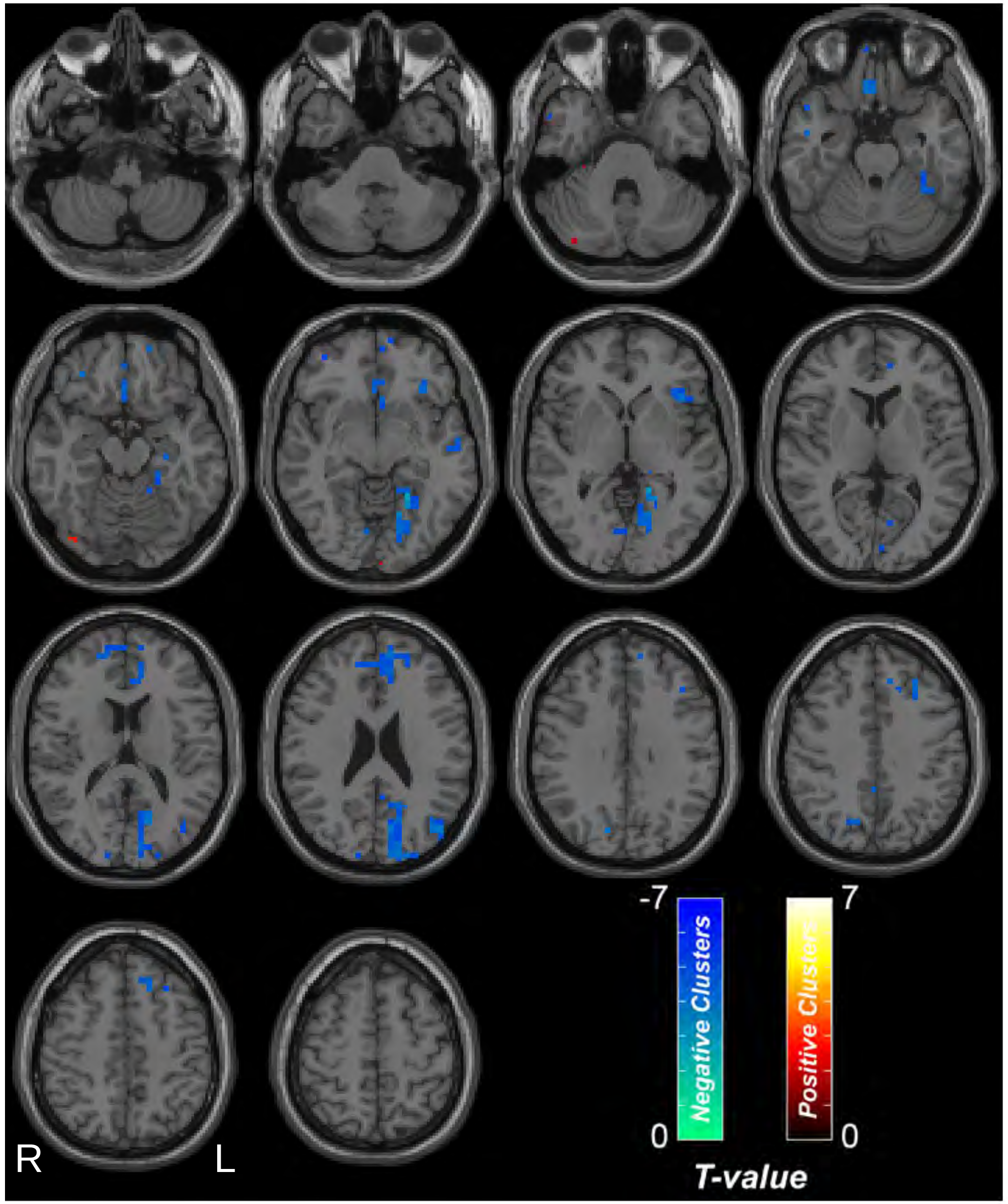


Kronemer et al., 2021

### Stimulus Presentation

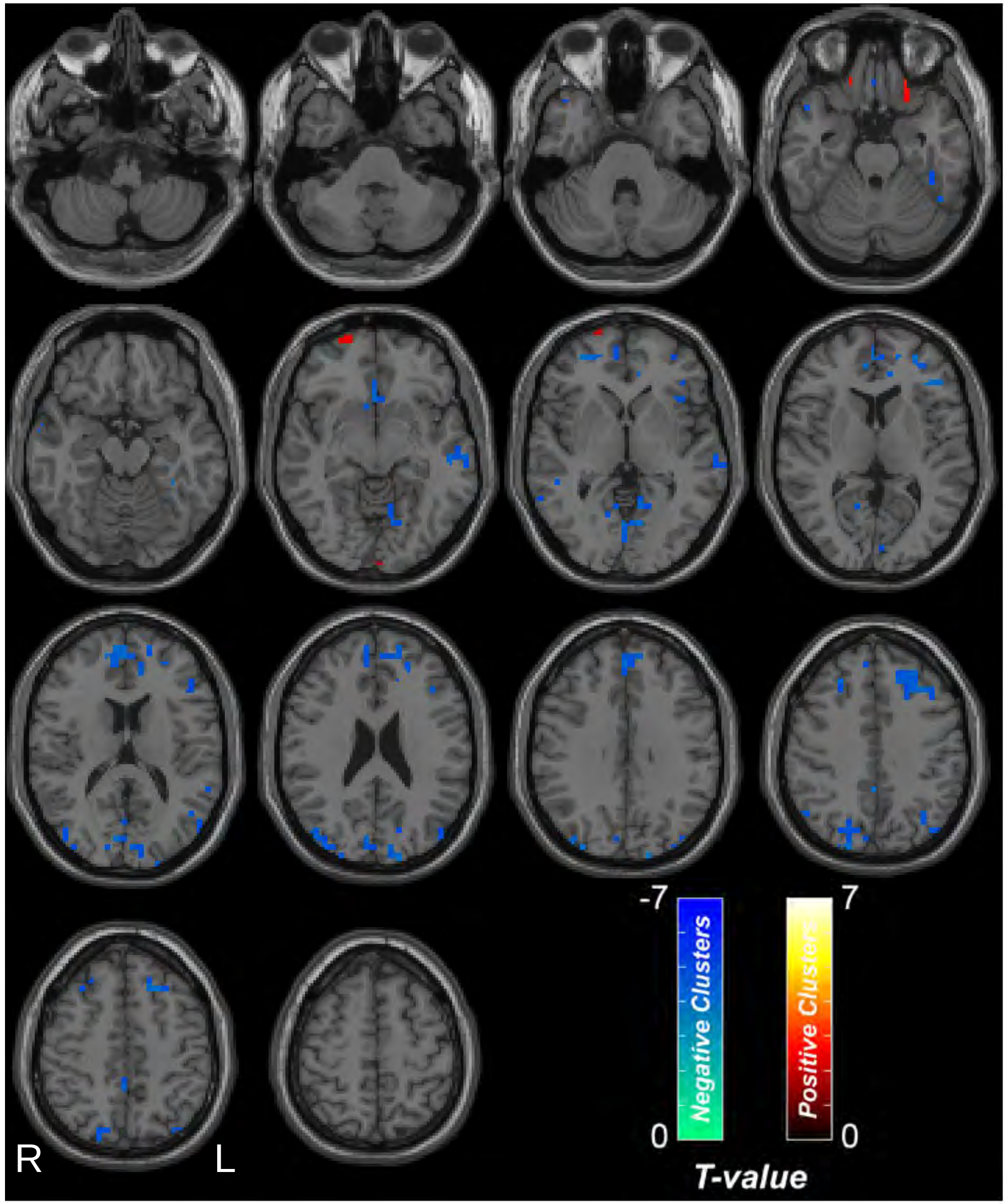


Kronemer et al., 2021

### 1s post-stimulus

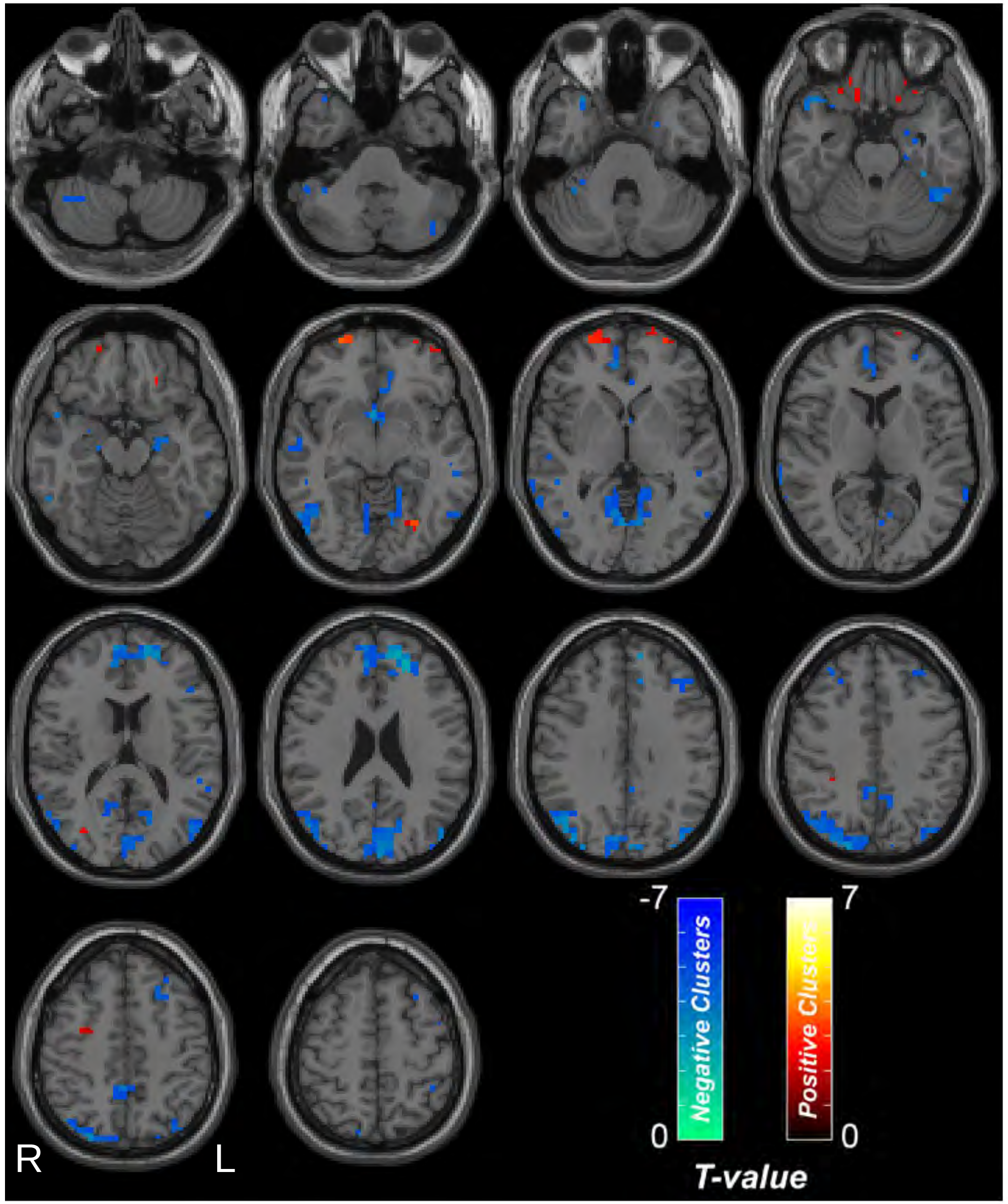


Kronemer et al., 2021

### 2s post-stimulus

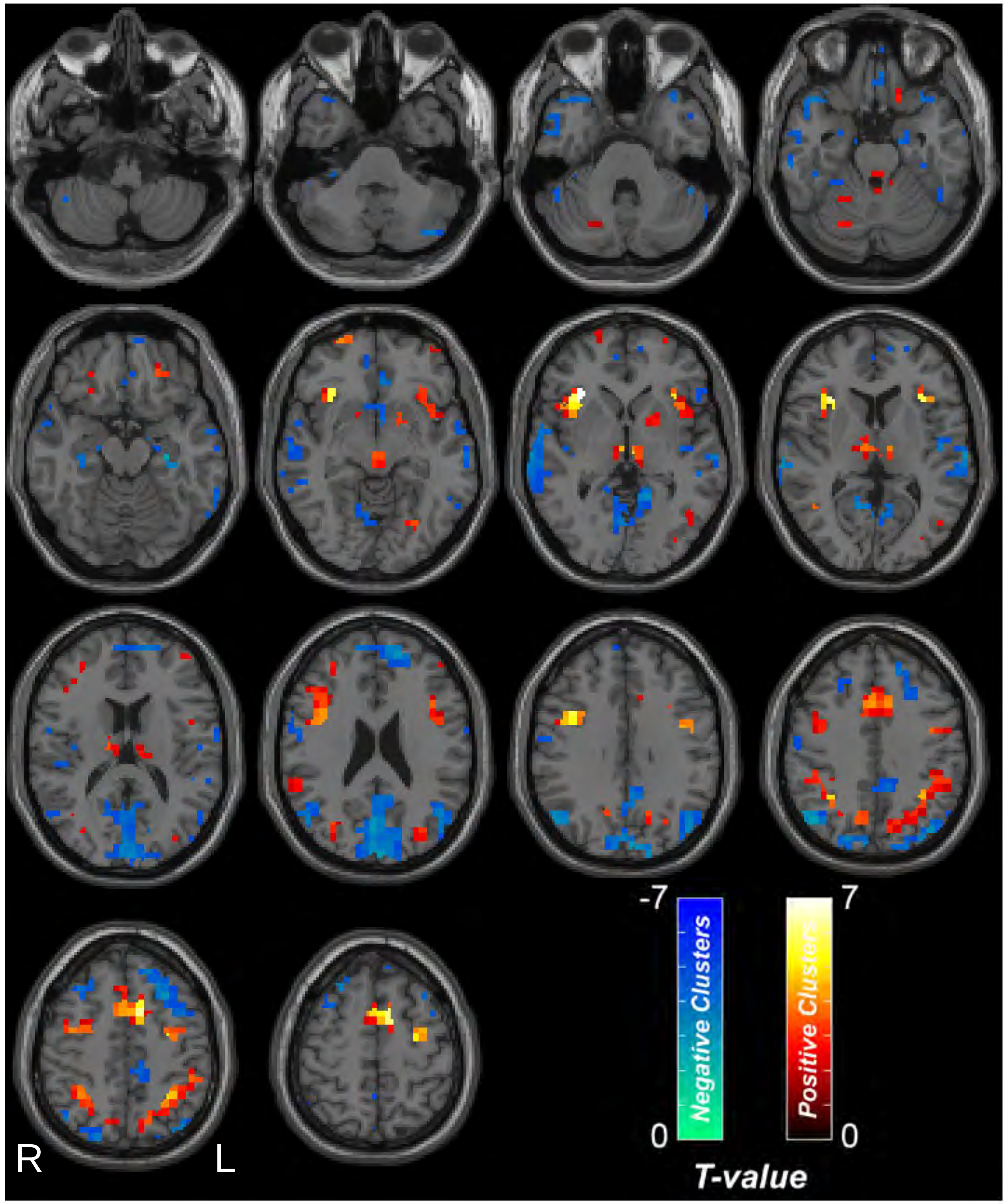


Kronemer et al., 2021

### 3s post-stimulus

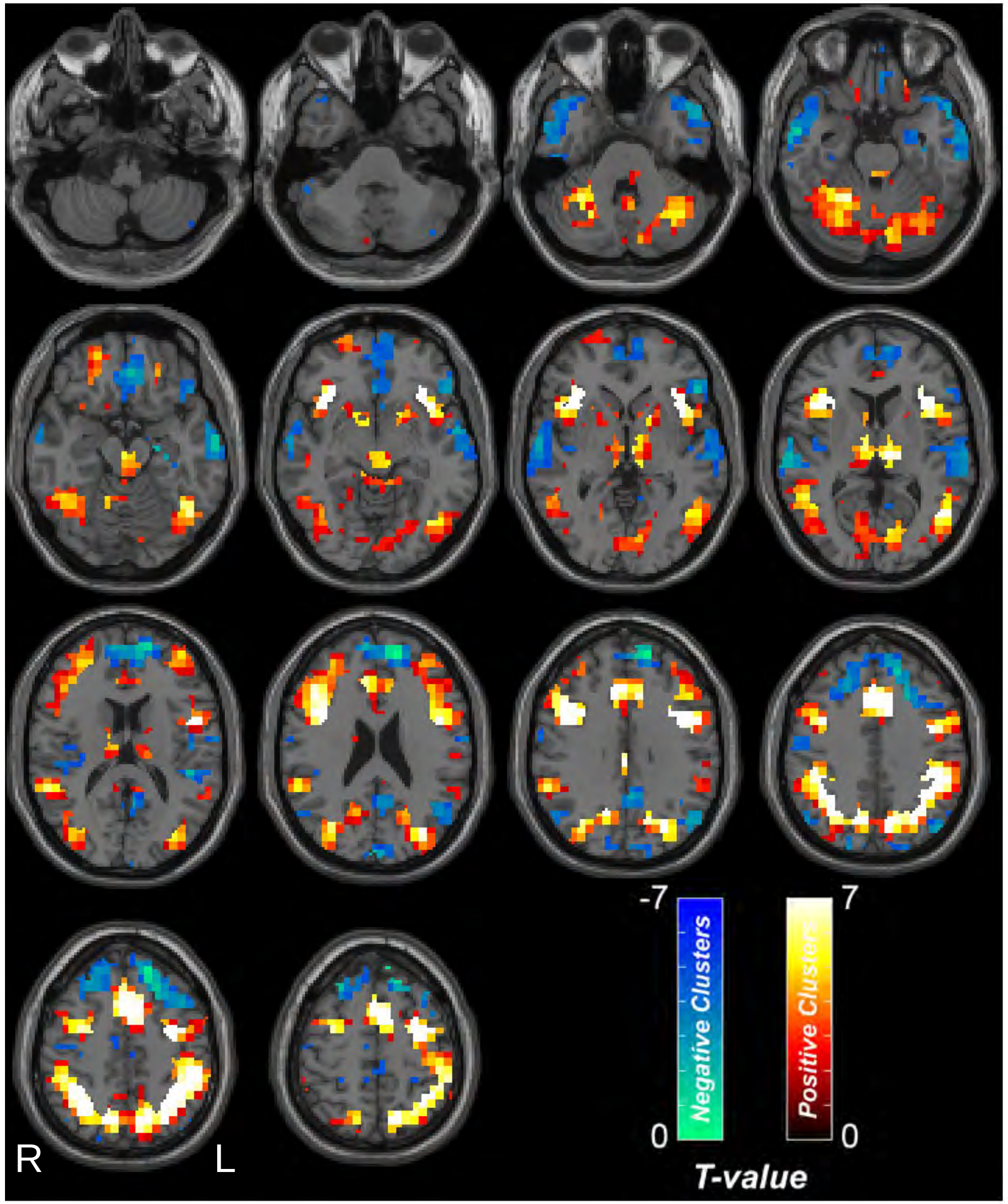


Kronemer et al., 2021

### 4s post-stimulus

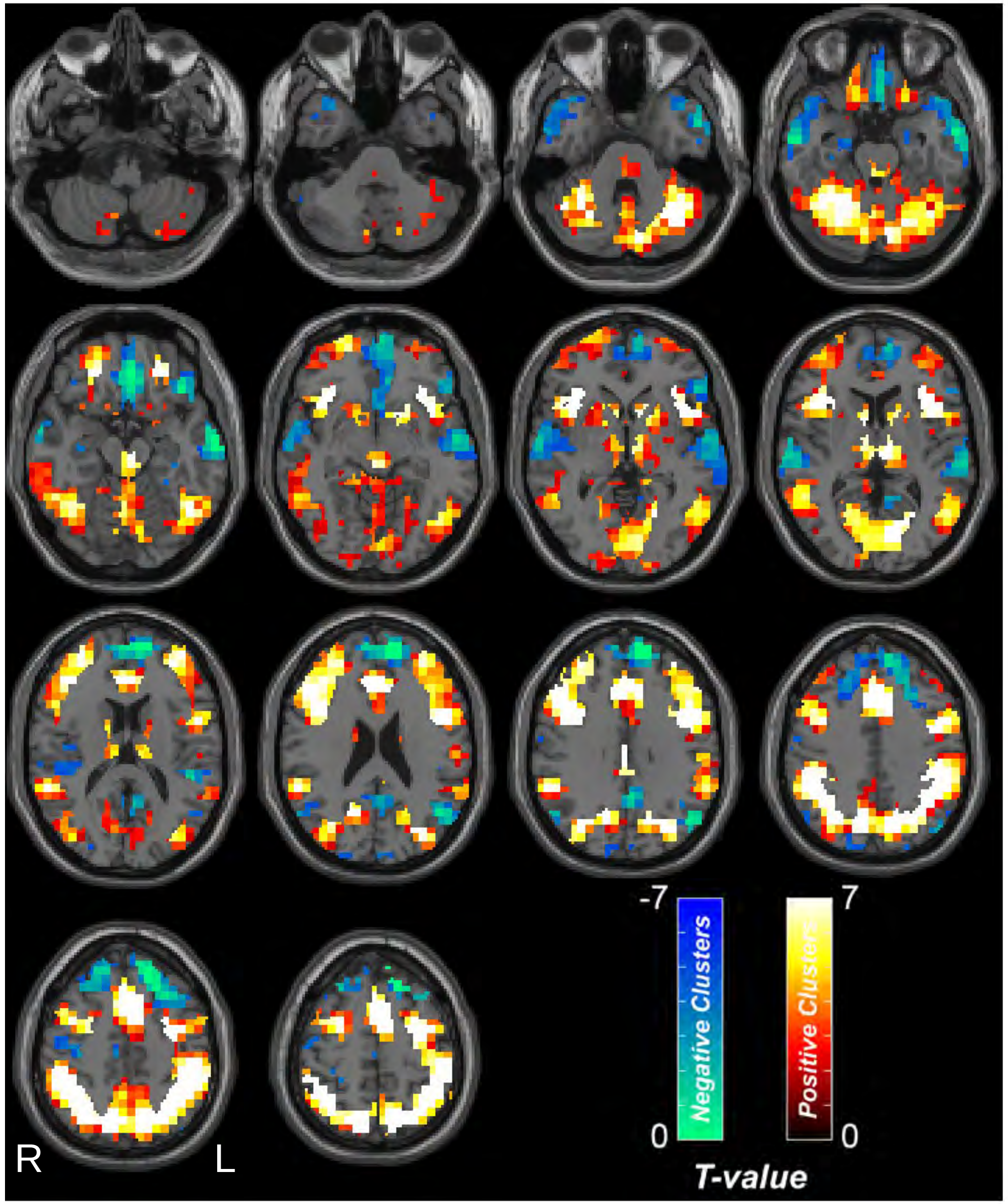


Kronemer et al., 2021

### 5s post-stimulus

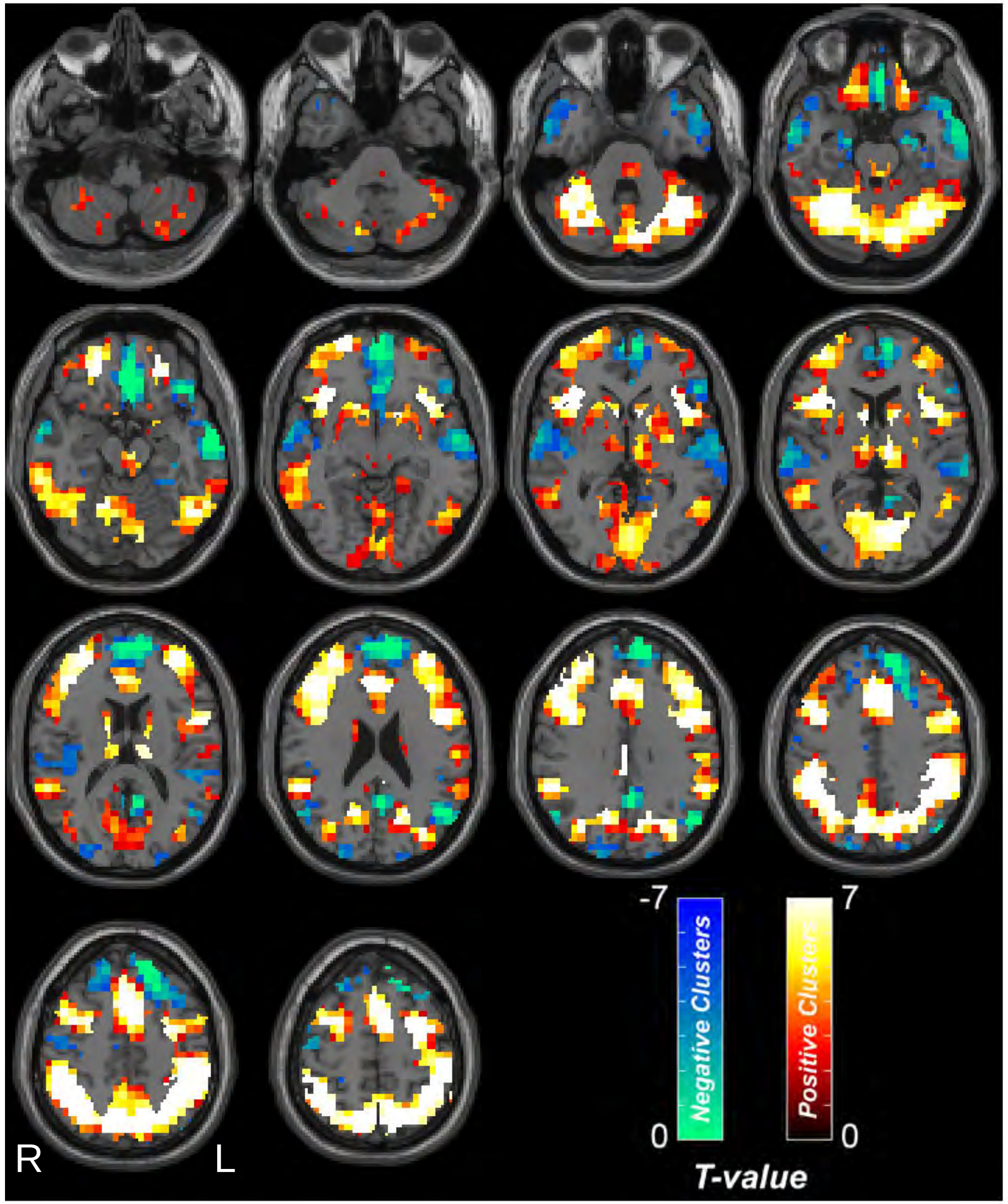


Kronemer et al., 2021

### 6s post-stimulus

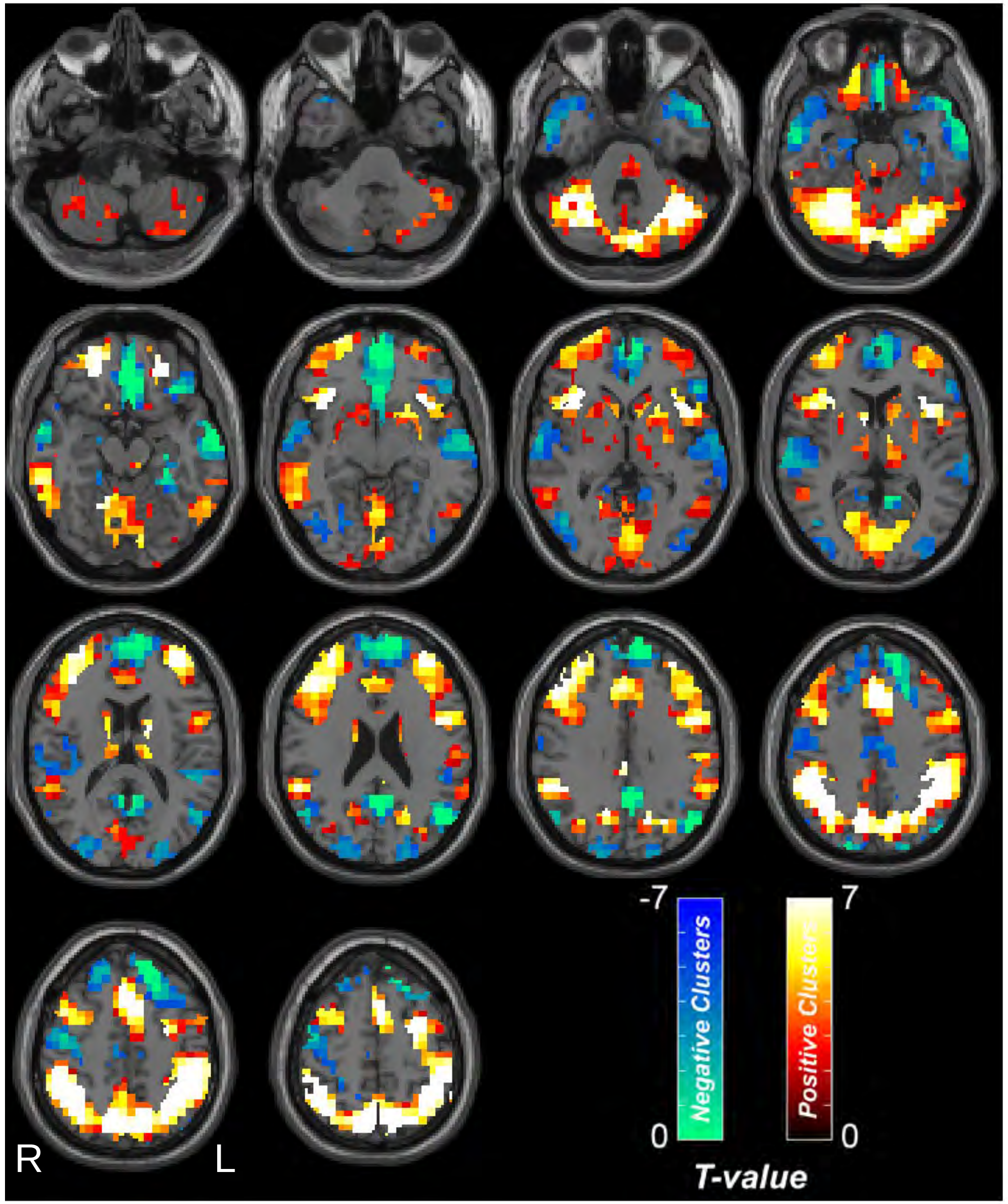


Kronemer et al., 2021

### 7s post-stimulus

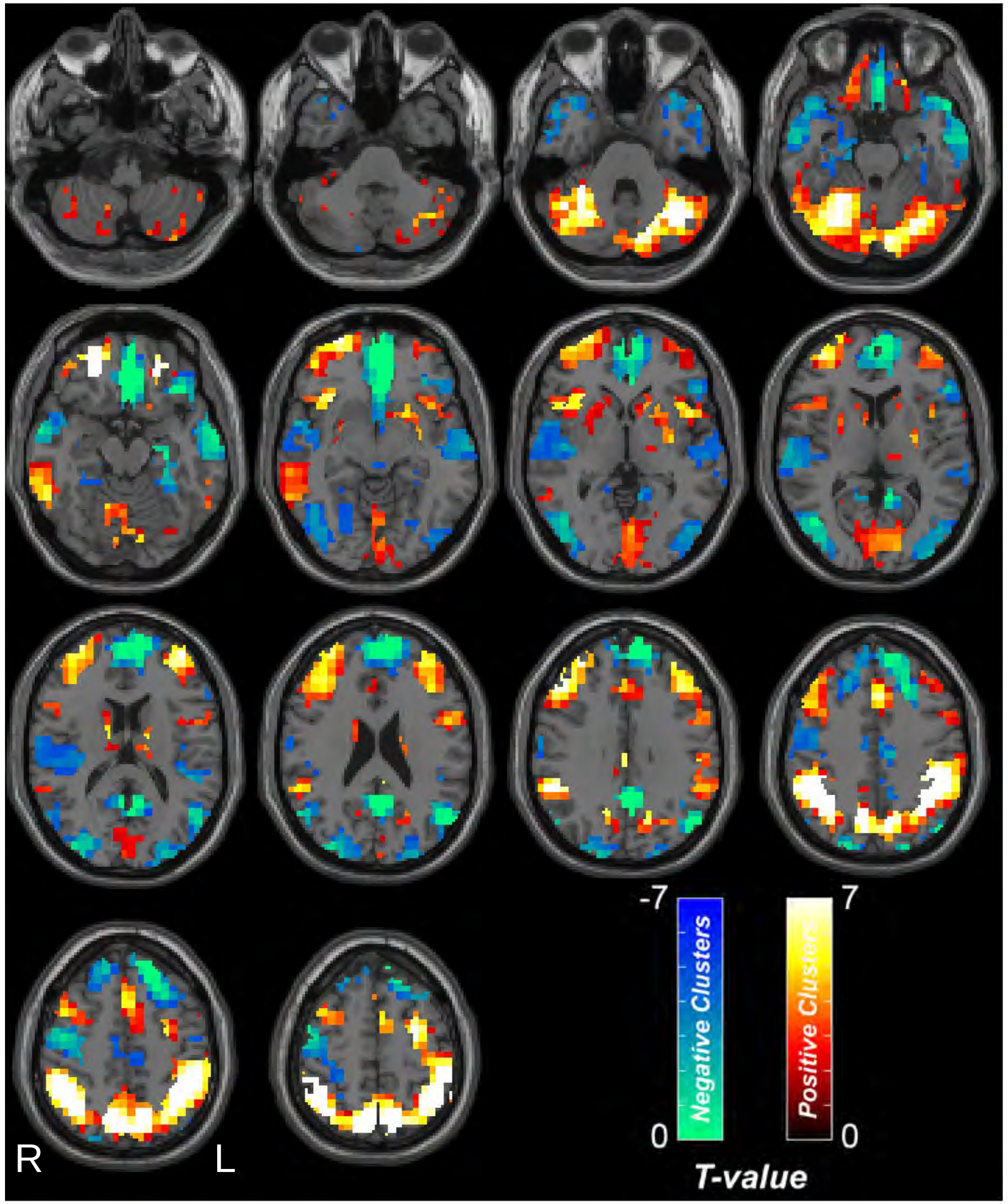


Kronemer et al., 2021

### 8s post-stimulus

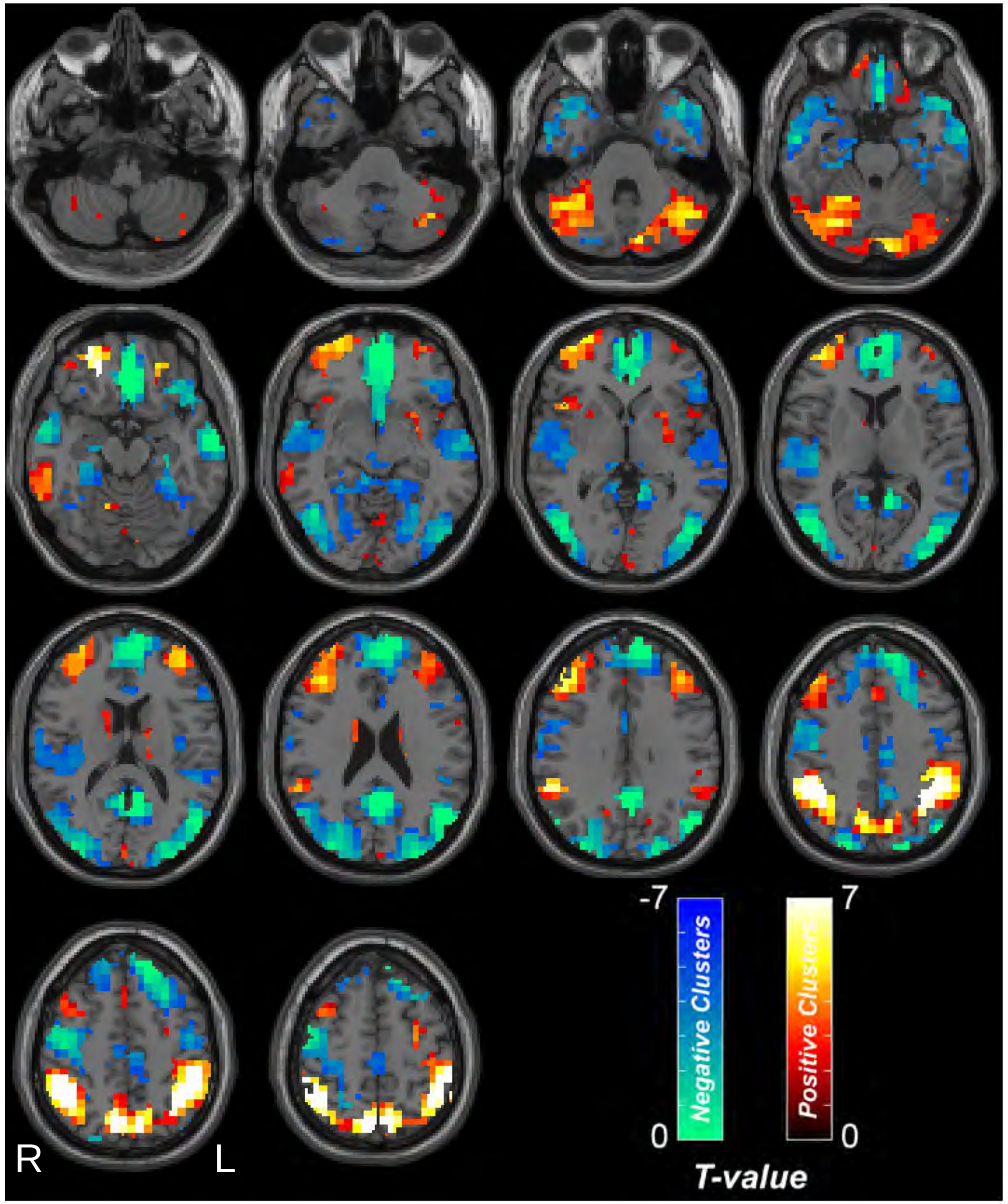


Kronemer et al., 2021

### 9s post-stimulus

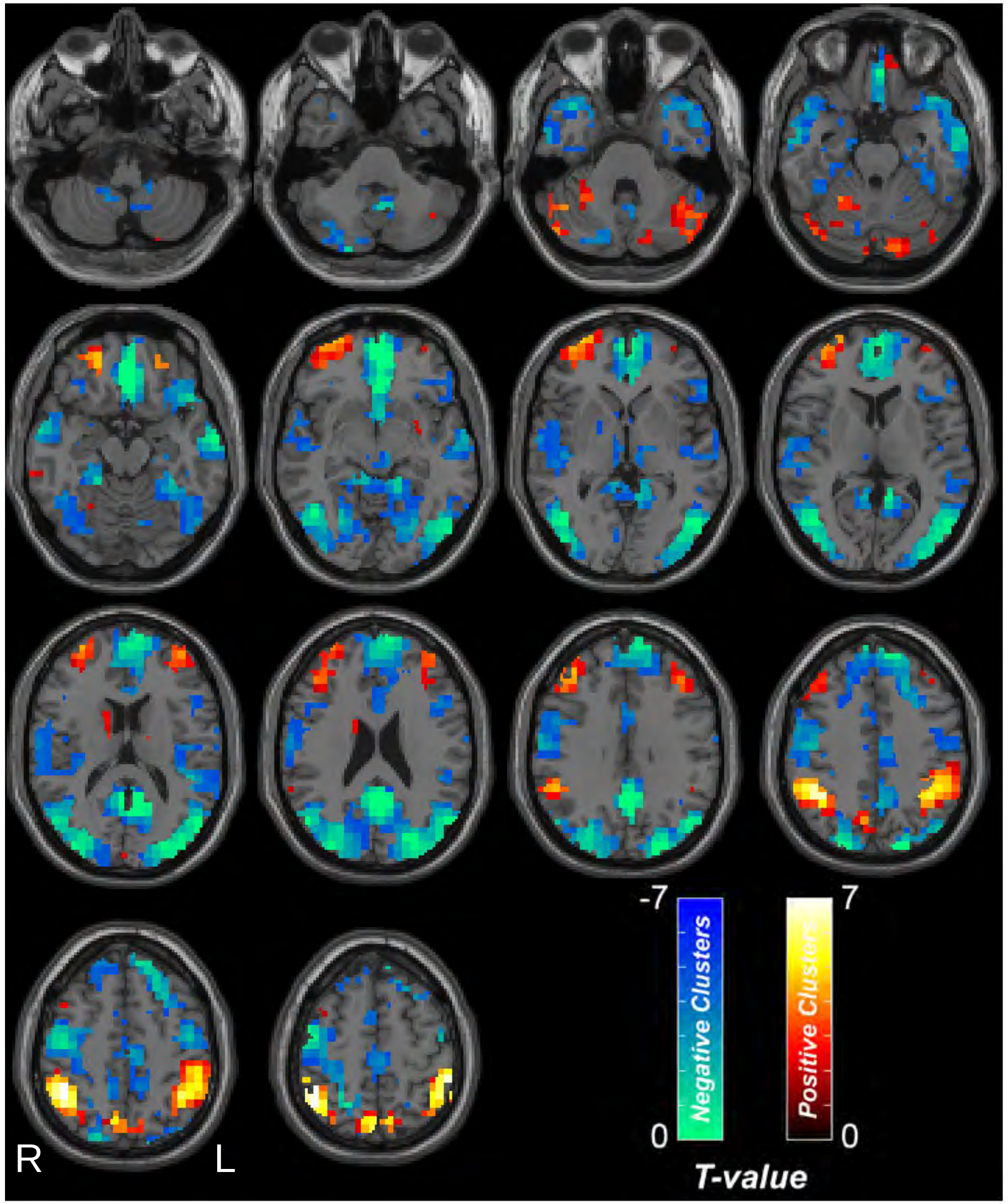


Kronemer et al., 2021

### 10s post-stimulus

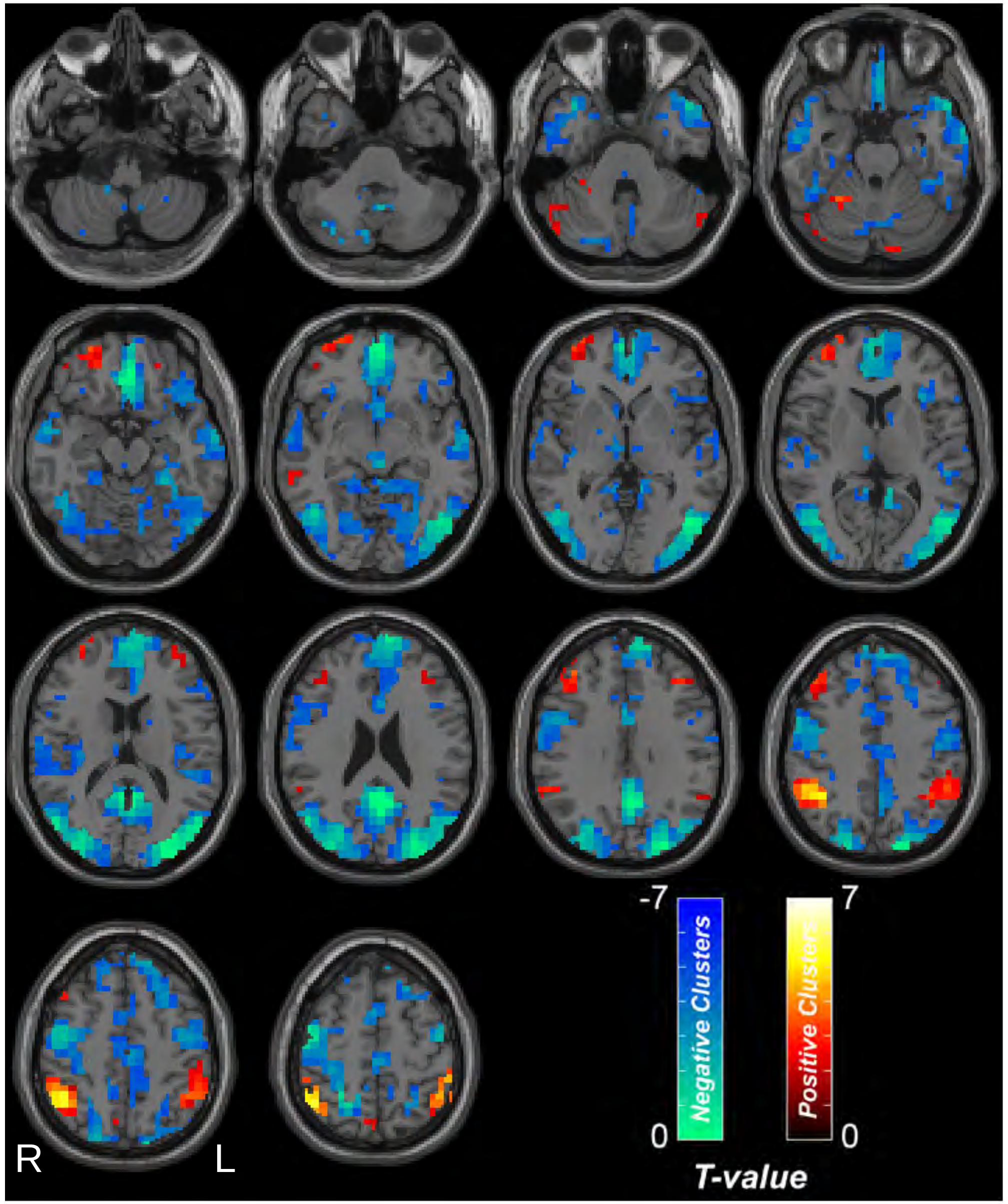


Kronemer et al., 2021

### 11s post-stimulus

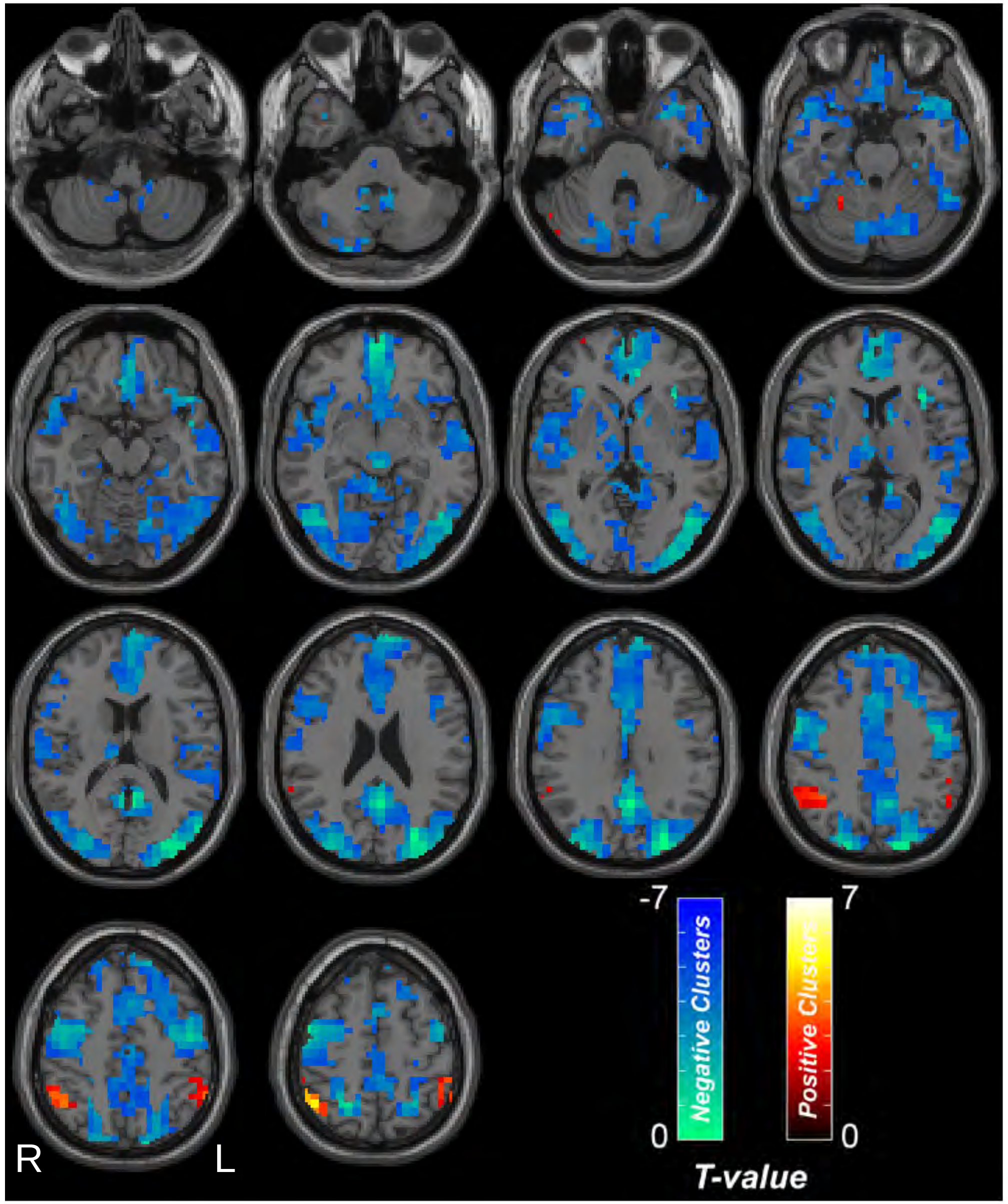


Kronemer et al., 2021

### 12s post-stimulus

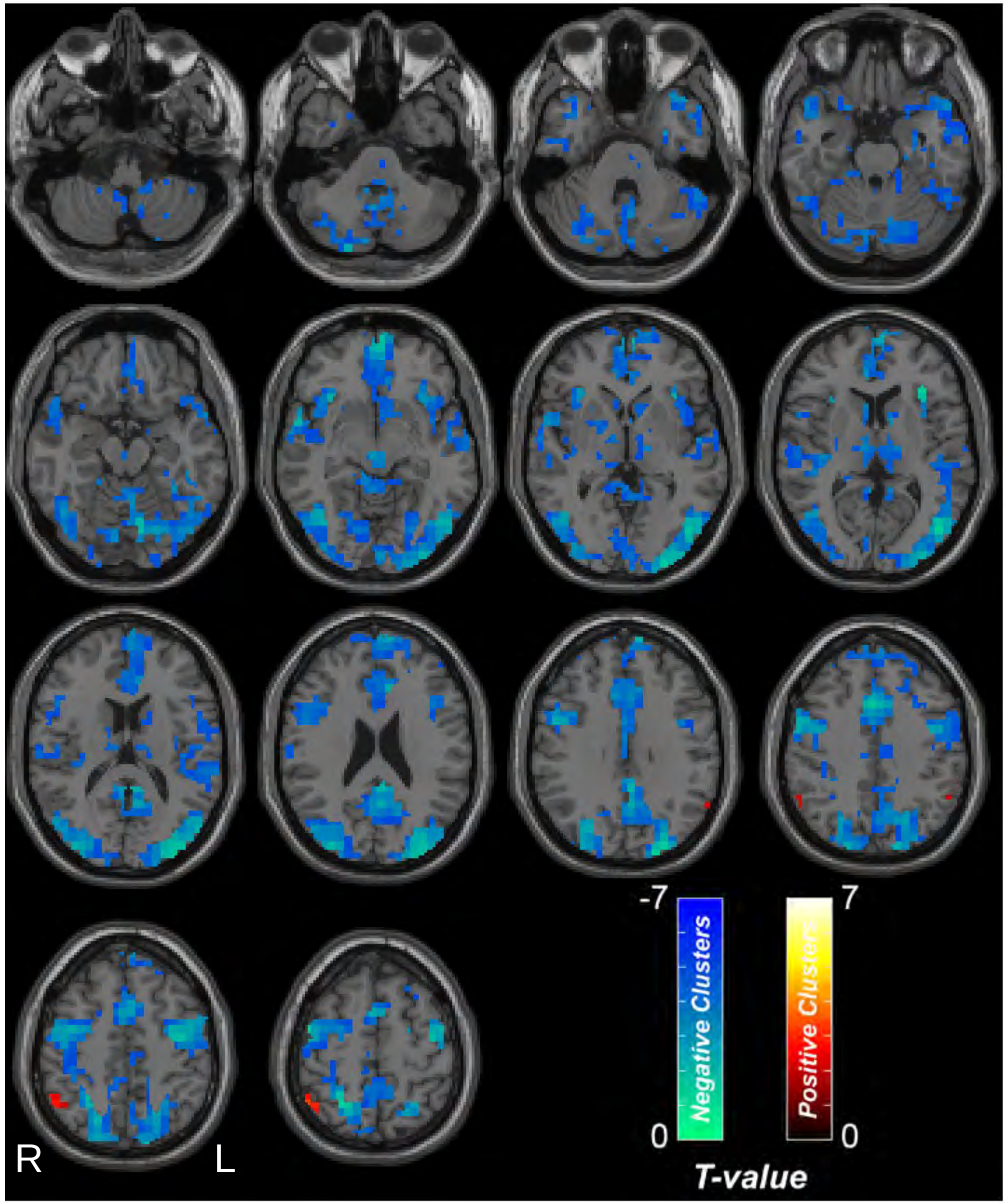


Kronemer et al., 2021

### 13s post-stimulus

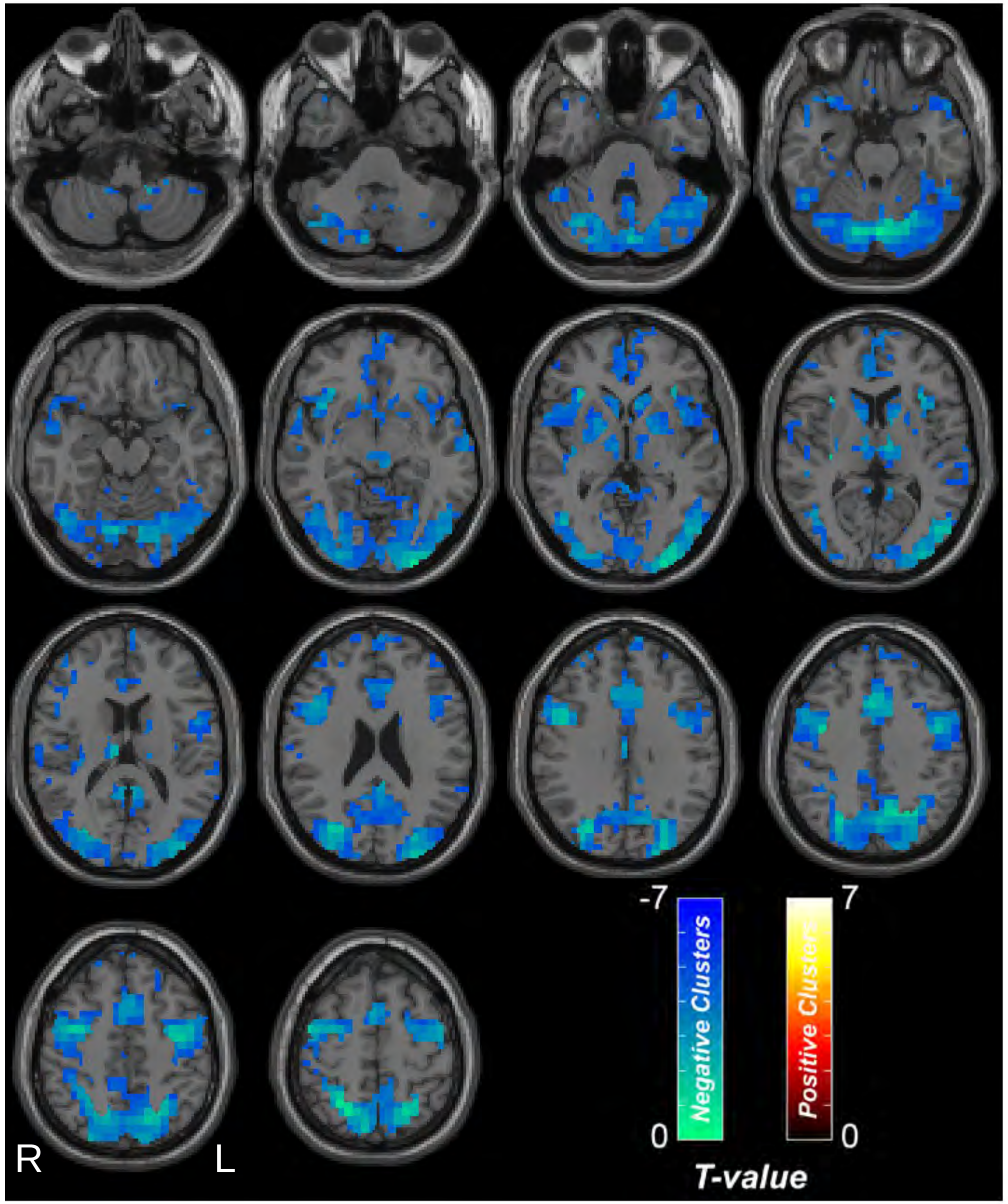


Kronemer et al., 2021

### 14s post-stimulus

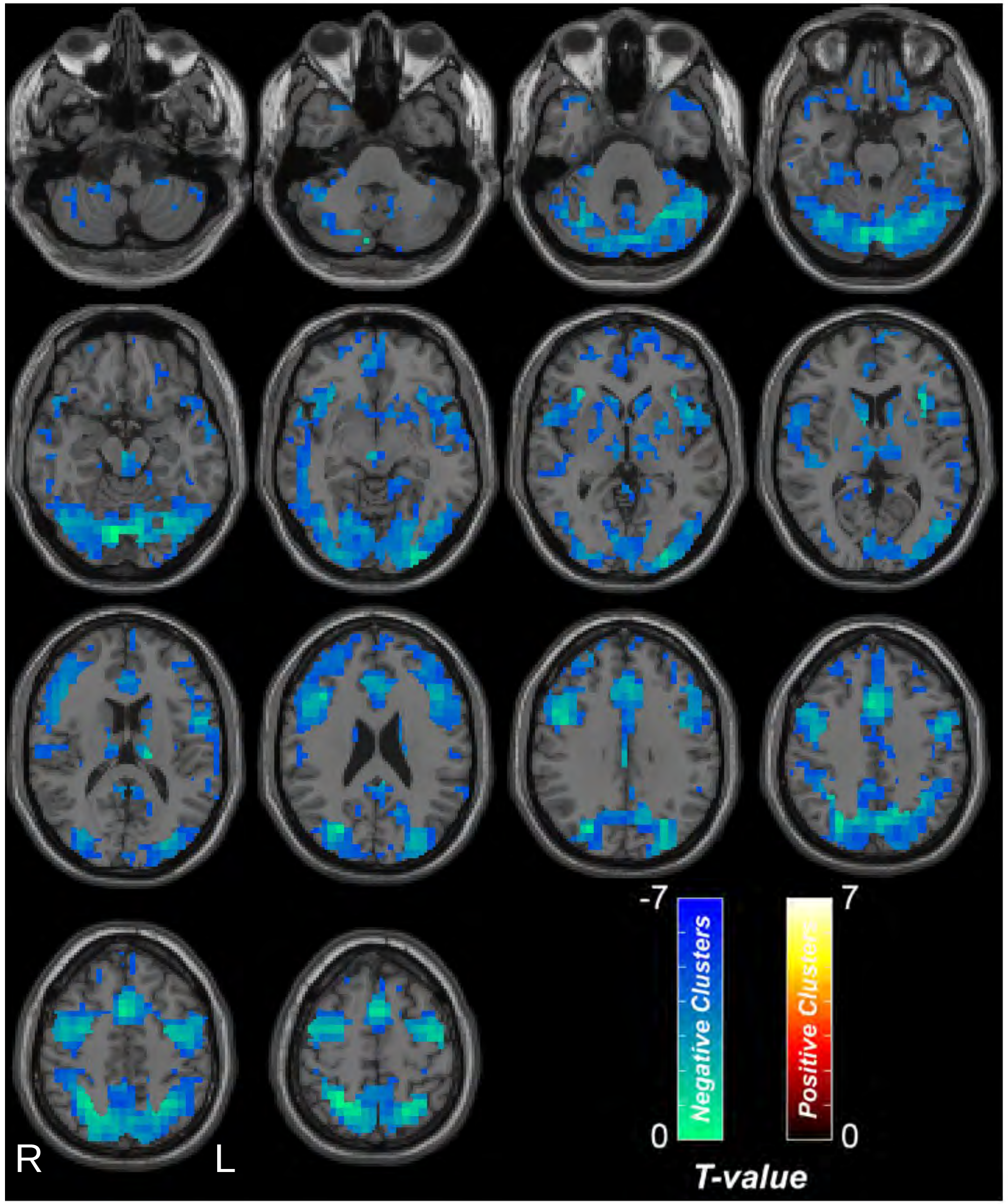


Kronemer et al., 2021

### 15s post-stimulus

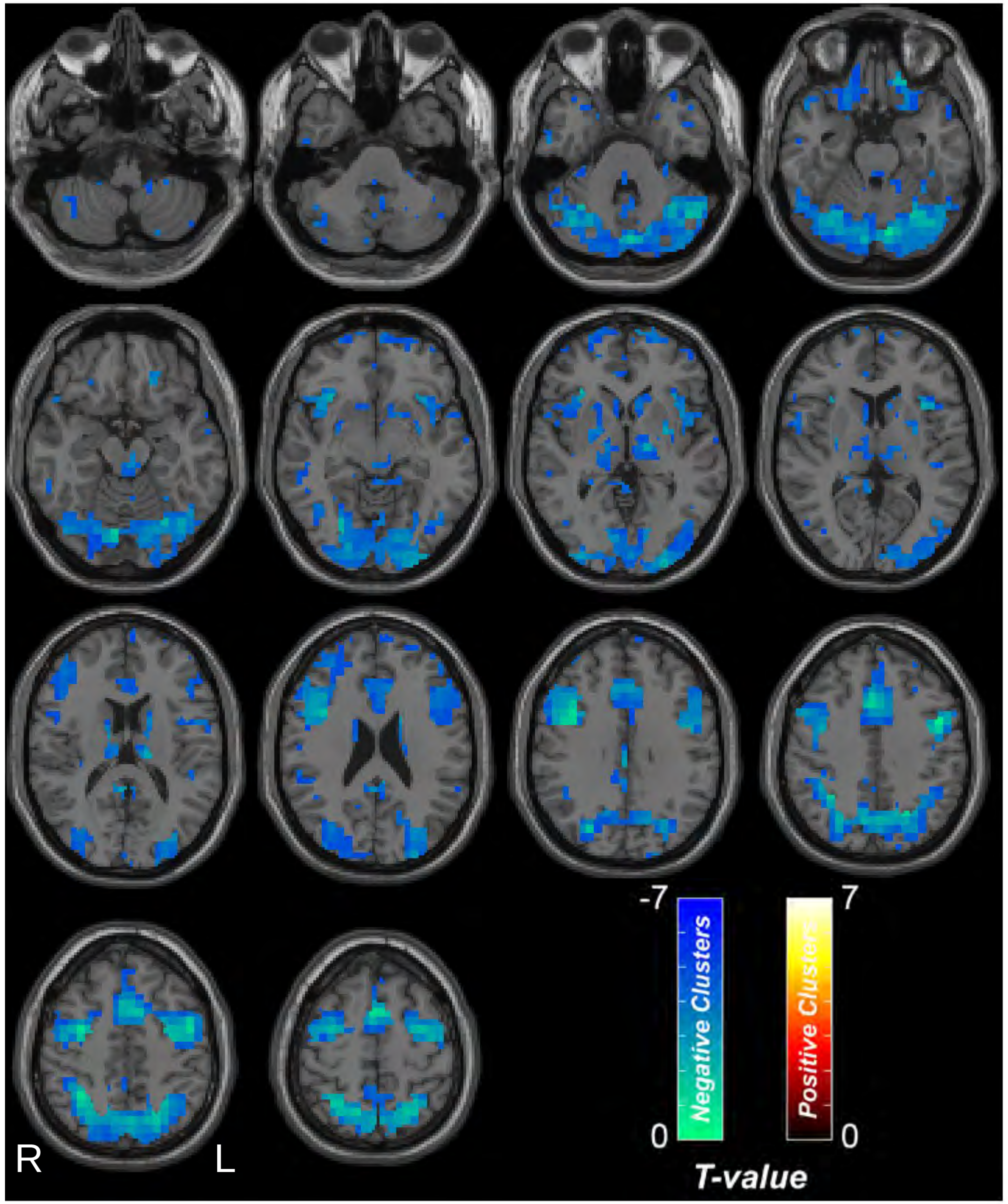


Kronemer et al., 2021

### 16s post-stimulus

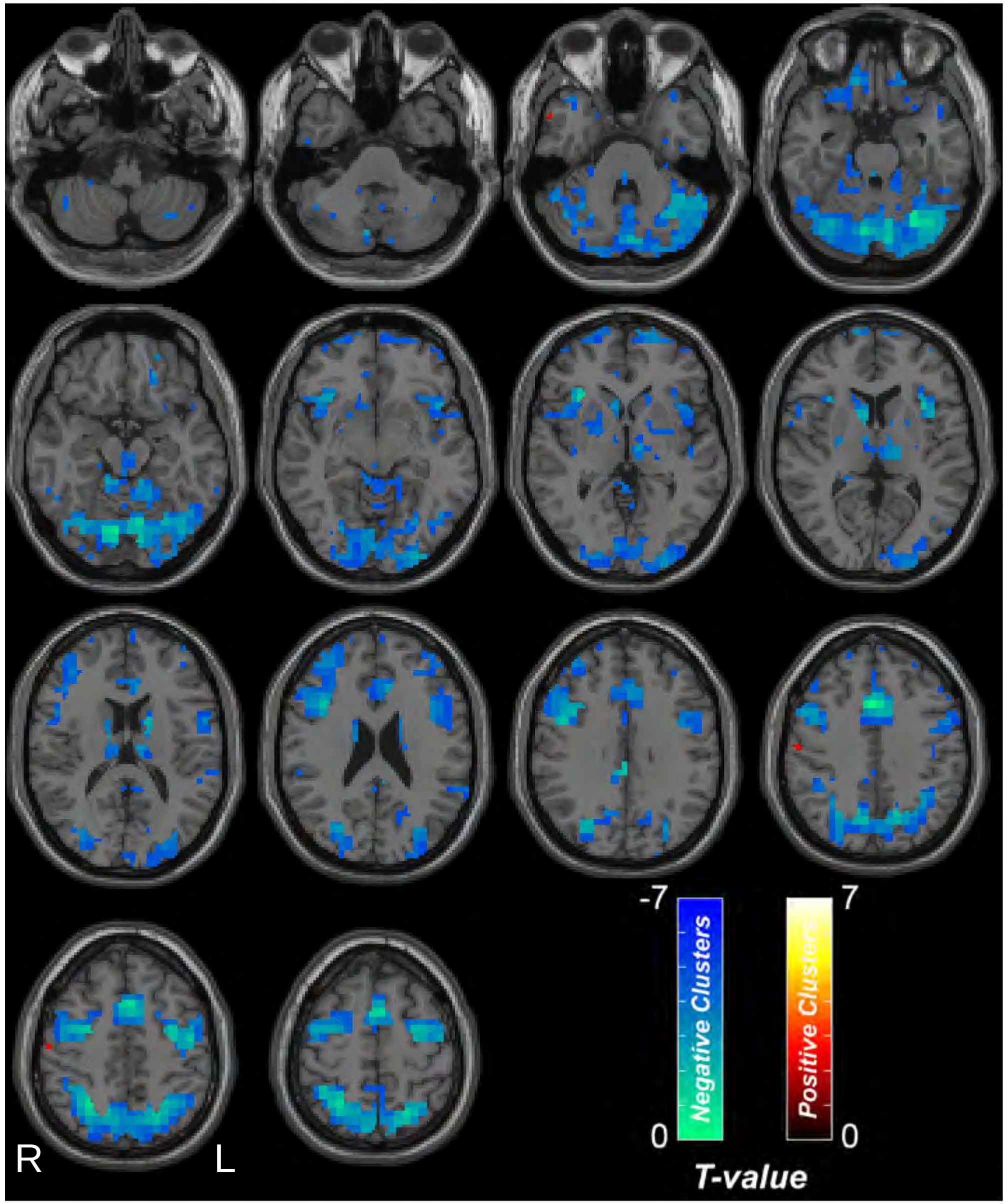


Kronemer et al., 2021

### 17s post-stimulus

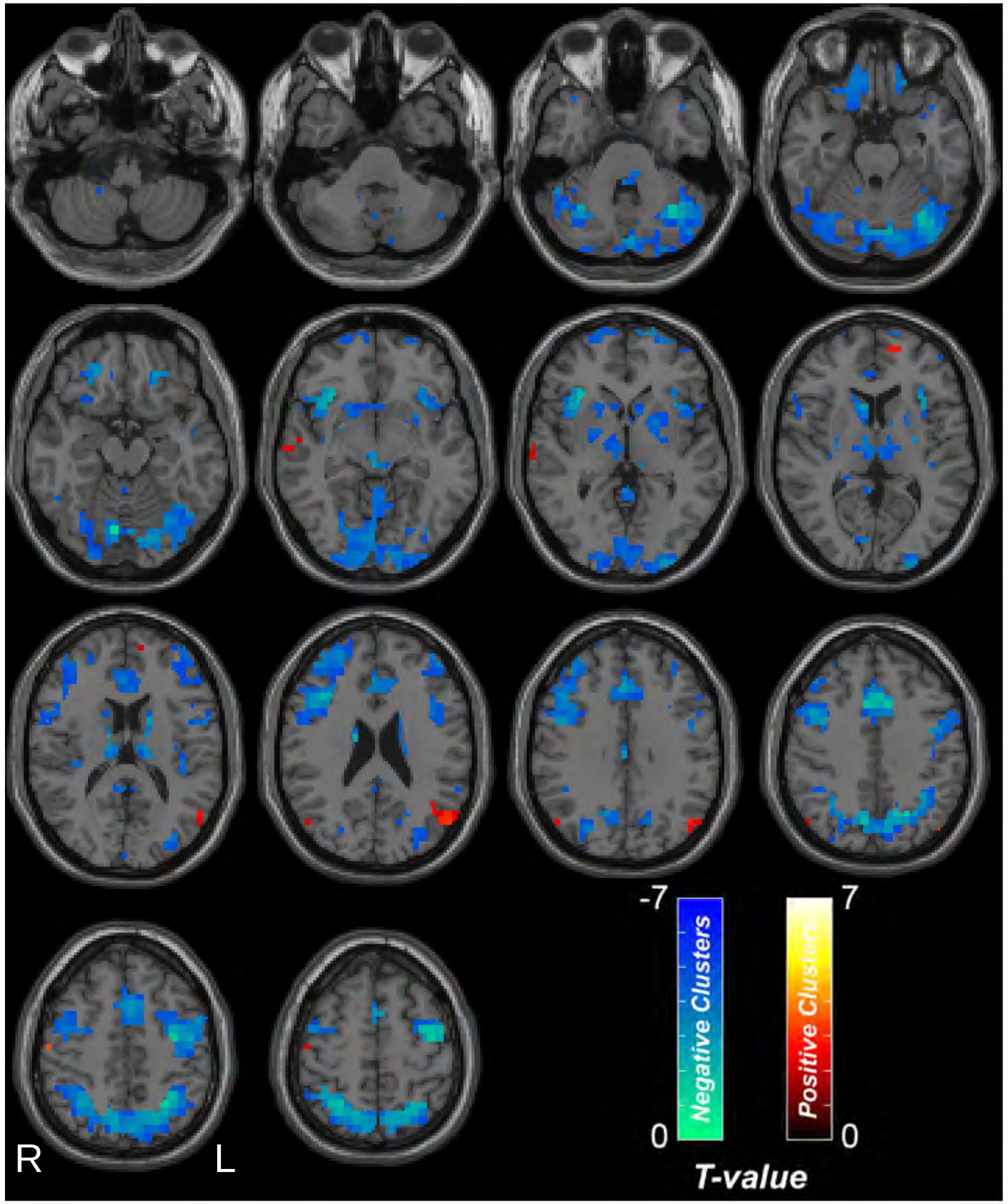


Kronemer et al., 2021

### 18s post-stimulus

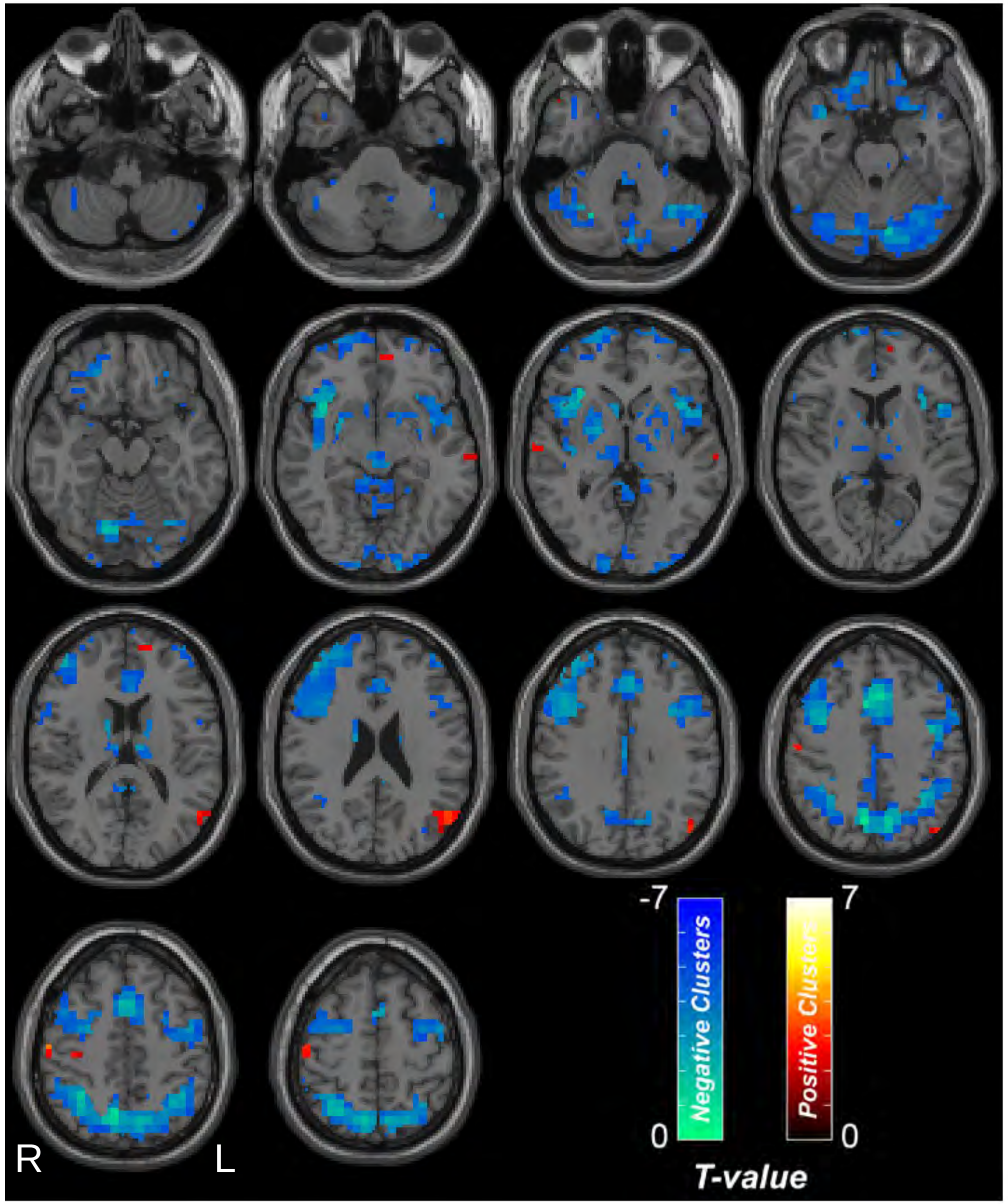


Kronemer et al., 2021

### 19s post-stimulus

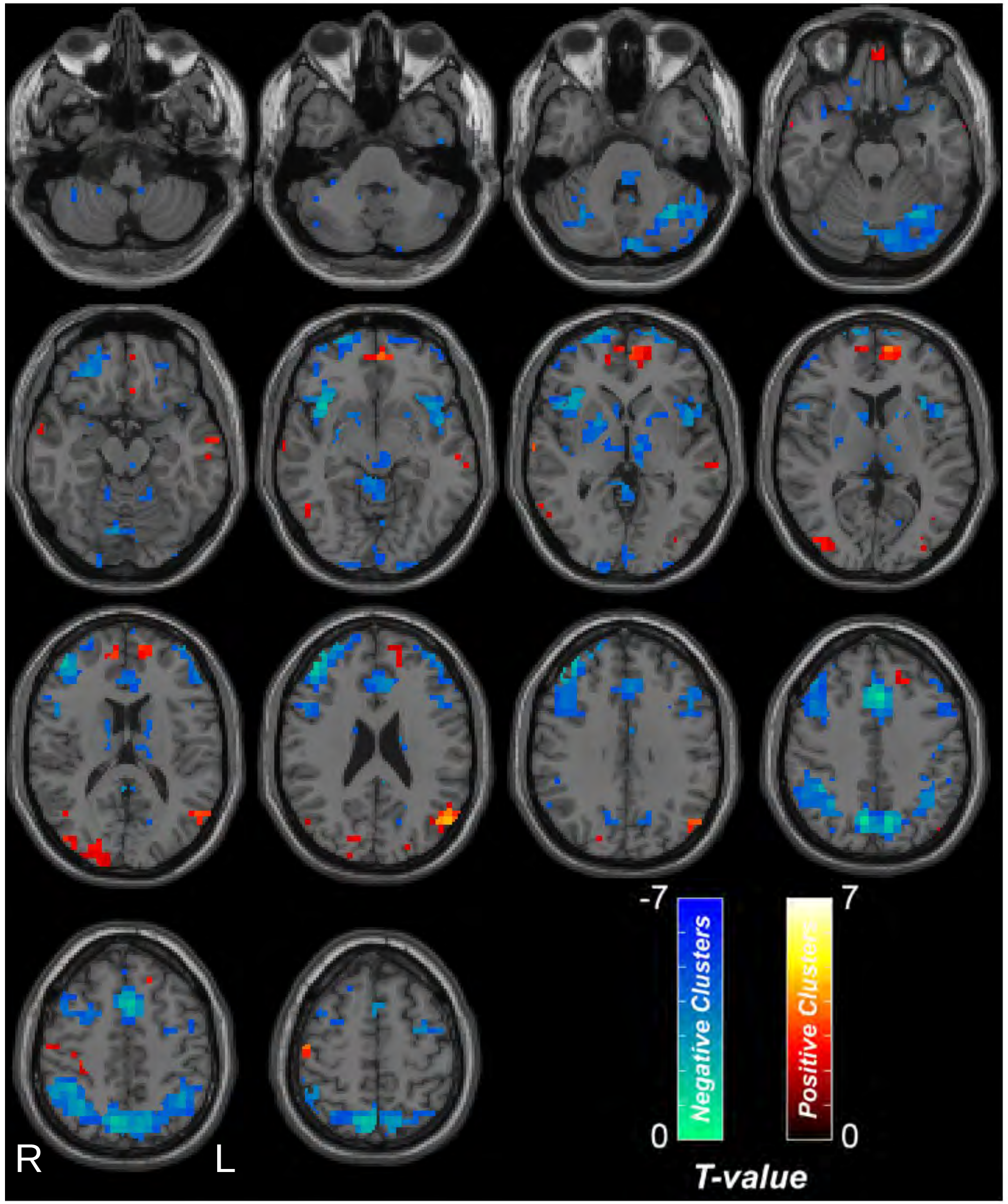


Kronemer et al., 2021

### 20s post-stimulus

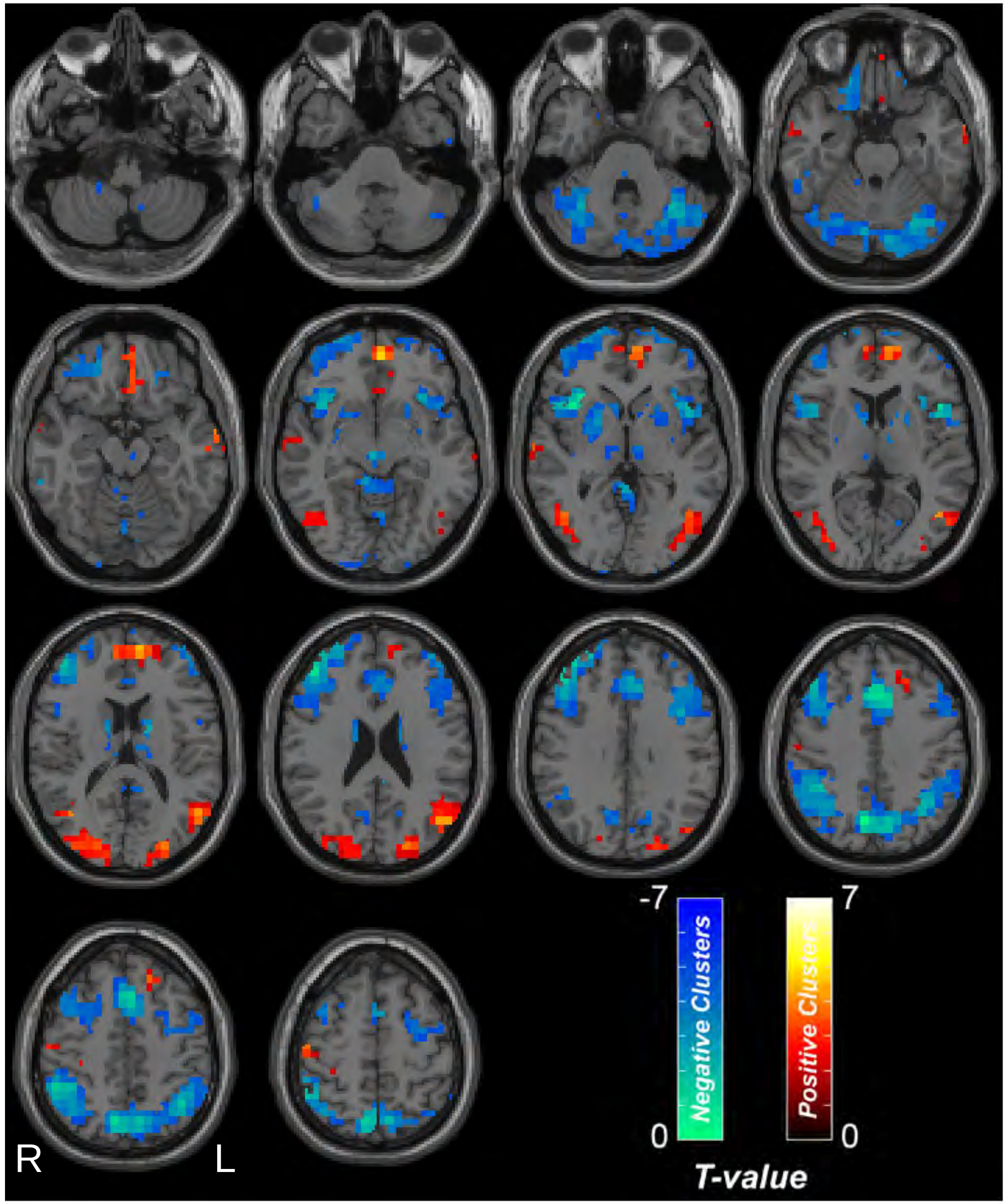


Kronemer et al., 2021
